# Evolutionary dynamics of the arthropod moulting machinery

**DOI:** 10.64898/2025.12.24.696370

**Authors:** Giulia Campli, Sagane Joye, Olga Volovych, Asya Novikov, Ariel D. Chipman, Marc Robinson-Rechavi, Robert M. Waterhouse

## Abstract

Exoskeletons define arthropods, providing support for segmented bodies and appendages while protecting against environmental stress and predation. Although ubiquitous, this evolutionarily variable feature has enabled arthropods to occupy diverse lifestyles and ecological niches, contributing to their unrivalled diversity. Because the chitinous cuticle is rigid, exoskeletons must be periodically shed and replaced as animals grow. Arthropods therefore develop through discrete moults, with conserved phases that progress from pre-moult preparation to ecdysis and post-moult maturation. These are tightly regulated developmental transitions controlled by a molecular toolkit comprising neuropeptides, hormone-synthesising enzymes, receptors, and the early, fate, and late gene sets that activate and execute the moulting process. Although genetic studies in model species have identified many components, major knowledge gaps remain, especially in non-insect arthropods. Advances in genome sequencing now enable comparative genomic analyses across diverse, previously understudied arthropod lineages to begin to address these gaps. We present a comprehensive comparative genomic survey of arthropods, sampling all four subphyla: Chelicerata, Myriapoda, Crustacea, and Hexapoda. Orthology inference and gene copy-number analyses contrast stable and dynamic components of the moulting machinery across the phylum. Ancestral state reconstructions and phylogenetic reconciliations reveal gene duplication and loss dynamics and the evolutionary histories of key moulting gene families. The inclusion of newly generated myriapod genomes addresses a major taxonomic gap and enables inferences of gene repertoire changes in the Mandibulata ancestor. The broad taxonomic representation enables a phylum-wide assessment to systematically evaluate, refine, and revise current understanding of the evolutionary dynamics of the entire moulting genetic toolkit.

## Introduction

Development via periodic moulting represents a hallmark of all extant arthropod subphyla and their extinct ancestors. Their exoskeletons provide both physical support and protection, but need to be replaced to allow for normal growth and maturation. Arthropod life histories are therefore punctuated by ecdysis events. When internal pressure opens fractures along sutural lines in the rigid chitin, the individual can escape from the former exoskeleton and replace it with a new one. Synthesis of the new, larger exoskeleton starts well before this ecdysis event. It is part of a broader process that also includes a preparatory phase before the exit from the exuviae, and a terminal phase encompassing the maturation of the newly formed sheath. Across species, these phases vary in time span, and the number of moults occurring during the entire life cycle also varies substantially. For example, a fruit fly undergoes only three moults over the course of a few days, while horseshoe crabs normally moult yearly over decades (Tetlie et al. 2008). Large variations are also found in the length of the intermoult period, where the arthropod feeds and grows until reaching a size where the exoskeleton constrains growth (Nijhout et al. 2014). Remarkably, moulting allows growth not only by an increase in body dimensions but also by the postembryonic addition of new segments (Fusco and Minelli 2021; Chipman 2025). This process of anamorphosis characterises the development of several lineages, both extinct, such as Trilobita and other arthropod stem-groups, and extant, such as the early branching pycnogonid chelicerates, all diplopod myriapods and many centipedes, and several crustaceans (Fusco and Minelli 2021). In contrast, epimorphic development proceeds with no changes in the number of segments after hatching. Moulting is therefore both a ubiquitous yet highly variable process. This has enabled arthropods to adopt a wide variety of lifestyles and to exploit different ecological niches, for example in holometabolous insects it mediates the metamorphic transition to adulthood via a pupal stage.

Moulting events are carefully orchestrated transitionary processes in normal growth and development. These are controlled by a toolkit of molecular components including neuropeptides, hormone-synthesising enzymes, and receptors, as well as the so-called early, late, and fate genes (Campli et al. 2024). Studies have shown that the ecdysteroid and sesquiterpenoid hormone receptors are the main moulting signal transducers, activated by hormones triggered by moult-inducing environmental or endogenous stimuli (Qu et al. 2015). Such hormonal pathways lead to the activation of regulatory networks of transcription factors that control the expression of sets of genes coding for structural components of the exoskeleton. Finally, declining hormonal titres initiate irreversible neuroresponses, culminating in ecdysis and post-ecdysis exoskeleton maturation. Characterisation of these molecular components has advanced through research leveraging well-established genetic techniques for experimental manipulations of a few tractable model species. This means that there remain substantial gaps in our understanding of the molecular mechanisms underlying the moulting process (Campli et al. 2024). Nevertheless, new opportunities to begin to address these gaps are growing rapidly as sequencing technologies enable the generation of large amounts of genomics data. Such resources facilitate the exploration of understudied arthropod lineages (Feron and Waterhouse 2022a), and the investigation of gene evolutionary histories exploiting an increasing number of species. These histories are governed by mechanisms of gene evolution including duplications, losses, and more rarely *de novo* gene birth, on which evolutionary forces act resulting in modifications to the repertoire of organismal functions (Thomas et al. 2020). Phenotypic innovations can arise not only from gene family expansions and contractions, but also from changes in the functional units of proteins, i.e. the gain, loss, and rearrangement of domains within the protein (Lees et al. 2016). Assessing such dynamics has provided evolutionary insights into processes such as immunity, digestion and metabolism, chemosensation, and reproduction (Palmer and Jiggins 2015; Thomas et al. 2020; Feuda et al. 2021; Ruzzante et al. 2022).

Here we present the results from conducting an extensive comparative genomic survey of putative moulting toolkit components employing a large set of species with representatives of all arthropod subphyla: Chelicerata, Myriapoda, Crustacea, and Hexapoda, where hexapods and the paraphyletic crustaceans together form the monophyletic Pancrustacea. Through comprehensive orthology delineation and gene copy-number profiling, we conduct a phylum-wide assessment of the stable and dynamic components of the moulting machinery functional modules. Complemented by ancestral state reconstructions and phylogenetic reconciliations, we investigate the evolutionary histories of moulting gene families across the arthropod phylogeny. Furthermore, *de novo* genome sequencing of myriapods contributes to the knowledge of this heretofore poorly represented subphylum, as well as allowing for the pinpointing of gene repertoire changes to the Mandibulata ancestor. Our results extend, in a systematic manner, the taxonomic breadth of interrogated lineages, and serve to confirm, challenge, and revise reports from the literature on the presence and putative functionality of moulting genetic toolkit components across Arthropoda.

## Results

### Broad taxonomic representation enables a phylum-wide assessment of the moulting machinery

Assessing and collating available and newly generated annotated genomes resulted in a dataset of 145 species spanning the four subphyla: Chelicerata, Myriapoda, Crustacea, and Hexapoda, comprising 35 orders, 57% of which are represented by multiple species (Figure 1). Sampling from publicly available genomic resources aimed to build a representative dataset that included arthropod species from a range of different orders. Selecting the best quality assemblies and annotations from over-represented clades (mainly insects) aimed to balance taxonomic representation across the phylum. To include Myriapoda representatives in the dataset, genome sequencing, assembly, and annotation were performed for three species: *Scolopendra cingulata* (Mediterranean banded centipede) and *Scutigera coleoptrata* (house centipede) from the class Chilopoda, and *Archispirostreptus syriacus* (black millipede) from the class Diplopoda. To investigate the evolutionary histories of genes implicated in moulting processes, this phylum-wide dataset was used to define species and order-level phylogenies (Figure 1A) and to delineate orthologous groups (OGs) at the last common ancestor (LCA) of Arthropoda (Figure 1B). Species tree estimation from single-copy orthologues resulted in a phylogeny congruent with previous large-scale phylogenomics findings (Misof et al. 2014; Schwentner et al. 2018; Howard et al. 2020). Graph-based orthology delineation resulted in the clustering of 80% of 2’047’671 annotated protein-coding genes into 74’214 OGs, each representing all genes descended from a single gene in the LCA. Together, the time-calibrated species phylogeny and orthology data (Supplementary Figure 1) provide the basis for reconstructing gene family evolutionary histories including duplication and loss events over the last ∼600 million years of arthropod evolution.

**Figure 1.**
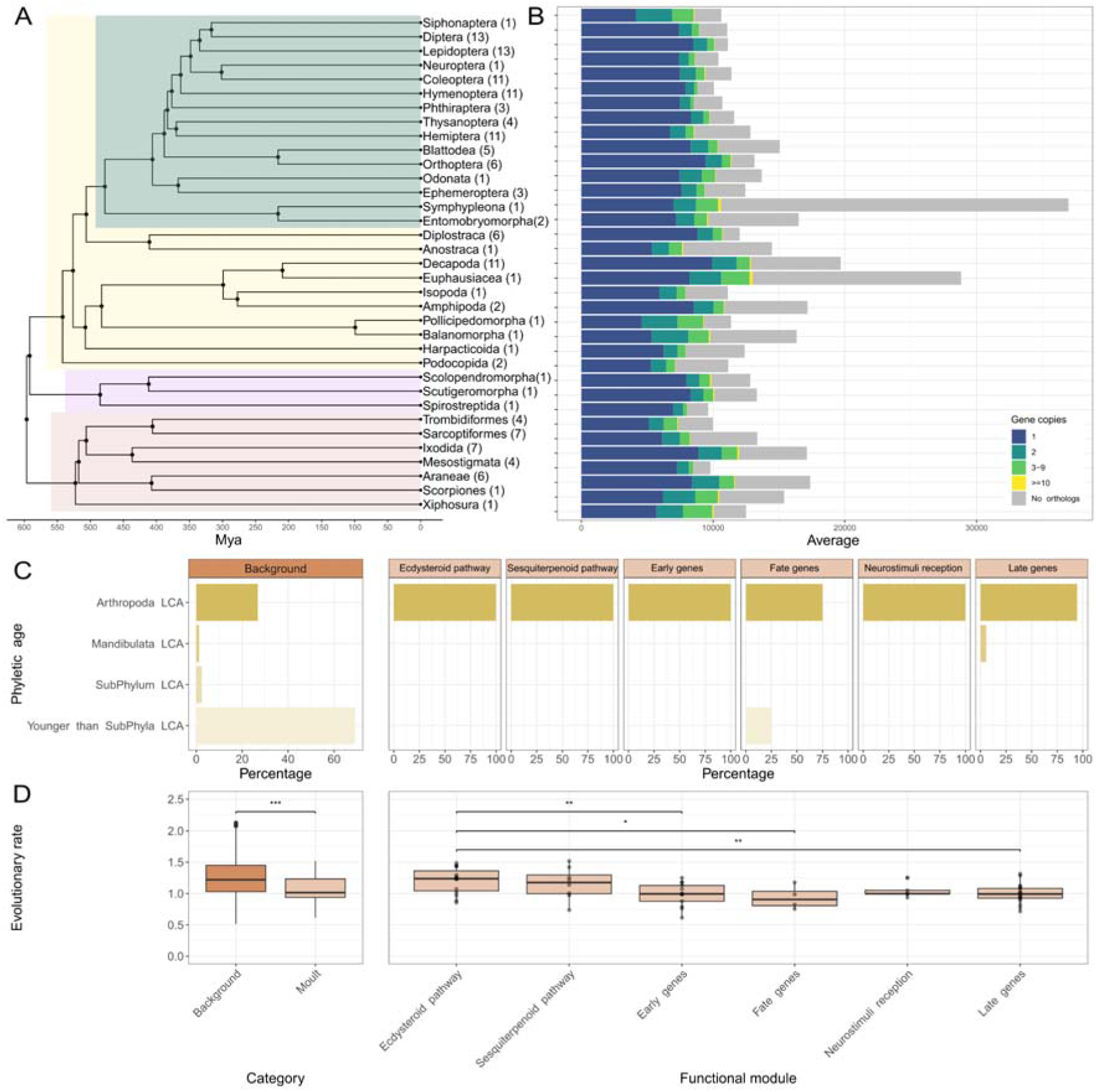
Phylum-wide phylogeny reconstruction and orthology delineation. **(A)** Relationships and estimated divergence times of the 35 orders from the complete dataset of 145 arthropod species. The number of species in each order is shown in parentheses at each leaf of the tree. The four arthropod subphyla are highlighted: Chelicerata (pink), Myriapoda (purple), Crustacea (yellow), and Hexapoda (green). The time-calibrated phylogeny was estimated using a super-alignment of protein sequences from single-copy orthologues. **(B)** Averaged across the species within each order, counts of genes in orthologous groups (OGs) according to copy number, and count of genes for which no orthologues could be identified. **(C)** Phyletic ages - defined as the last common ancestor (LCA) of all the species included in each OG - of all OGs (Background), and for OGs that comprise the six functional modules of moulting machinery gene families. **(D)** Distribution comparisons of OG average inter-species pairwise protein sequence divergence, normalised to the average of single-copy orthologues. Left: between all OGs and of the union of functional modules of moulting machinery gene families. Right: amongst the six functional modules of moulting machinery gene families. Asterisks indicate Wilcoxon test p-values of *** p<0.0005; ** p<0.005, and *p<0.05. The boxplots show the median, first and third quartiles, and lower and upper extremes of the distribution (1.5 × Interquartile range).

Querying the orthology dataset with curated lists of moulting machinery genes, *i.e.* components of the functional modules of moulting processes known from emerging and model systems, resulted in the identification of 63 OGs containing a total of 19’015 genes. The lists were based on extensive reviews of the accumulated knowledge of the genetic toolkit governing arthropod moulting, comprising 122 genes with evidence of involvement in various moulting processes. They represent the moulting-related gene families of interest, *i.e.* genes identified in one or more species as playing a role in moulting as well as their identifiable orthologues and paralogues from across Arthropoda (Campli et al. 2024). Based on the roles they play, these gene families can be grouped into six functional modules: the ecdysteroid and sesquiterpenoid pathways for hormone synthesis and signalling, the early gene and fate gene sets for activation of the moulting-induced programme, the toolkit of receptors of neuropeptides switching on the moulting process, and downstream late gene effectors for exoskeleton renewal. These classifications enable comparisons of the evolutionary histories of these functional modules, and the background of all other OGs, in terms of their phyletic ages (Figure 1C) and levels of protein sequence divergence (Figure 1D).

Phyletic ages are defined as the LCA of all the species included in the OG, simplified to stratifications of Arthropoda, Mandibulata, the four subphyla, and younger than subphylum level. The 145-species dataset allows for the designation of OGs spanning the Chelicerata-Mandibulata root to be inferred at least as old as the arthropod ancestor, *i.e.* OGs assigned a phyletic age of Arthropoda LCA may be older. All but two of the moulting machinery gene families were traceable to the arthropod ancestor, reflecting the ancient origins of genes involved in moulting processes (Figure 1C). This contrasts with the background of all other gene families, for which only 26% of OGs were traceable to the arthropod LCA and more than half emerged after the divergence of the four subphyla (Figure 1C). Evolution of new gene families, *i.e.* genes for which orthology can be traced to an internal node of the phylogeny but not to the arthropod root, therefore appears to contribute little to the evolution of the moulting machinery. However, functional studies in the many less well studied arthropod lineages may yet reveal novel innovations, whose origins can then be traced using comparative analyses across the phylum.

Evolutionary rates quantify sequence-level dynamics in terms of the levels of protein sequence divergence amongst OG member genes. Compared to other OGs, the moulting machinery gene families show overall significantly lower evolutionary rates, consistent with expected constraints associated with such an ancient and developmentally critical process (Figure 1D). While each moulting functional module shows some variation in evolutionary rates, the ecdysteroid pathway is overall the most divergent and is significantly higher than the early, late, and fate gene modules. The sesquiterpenoid pathway shows the same trend as the ecdysteroid pathway, but it is not significantly more divergent than the others. Genes implicated in moulting processes therefore generally show lower levels of sequence divergence, but members of the ecdysteroid and sesquiterpenoid pathways have elevated evolutionary rates compared to the other modules.

### Gene copy-number profiling reveals stable and dynamic components of the moulting machinery

Phylogenetic profiling across the 145-species dataset partitioned by the 35 orders and six functional modules identifies gene gains and losses across all modules and nearly all orders (Figure 2). These modules differ in their overall patterns, with the sesquiterpenoid pathway and early genes modules exhibiting mainly single-copy orthologues, the ecdysteroid pathway and late genes showing elevated proportions of duplications, and the neurostimuli module displaying the lowest proportion of single-copy orthologues. While detection of orthologues provides evidence of the maintenance of a gene family, orthologue absences may represent true gene losses or the failure to detect them. Asserting a loss from an individual genome therefore requires follow-up investigations to support the loss hypothesis. However, focussing on profiling results where absences characterise multiple species within a lineage provides support to assert ancestral loss events. This profiling analysis reveals broad trends across the 35 orders, which vary greatly amongst their member genes (Supplementary Figures 2-7), detailed here for each functional module.

**Figure 2.**
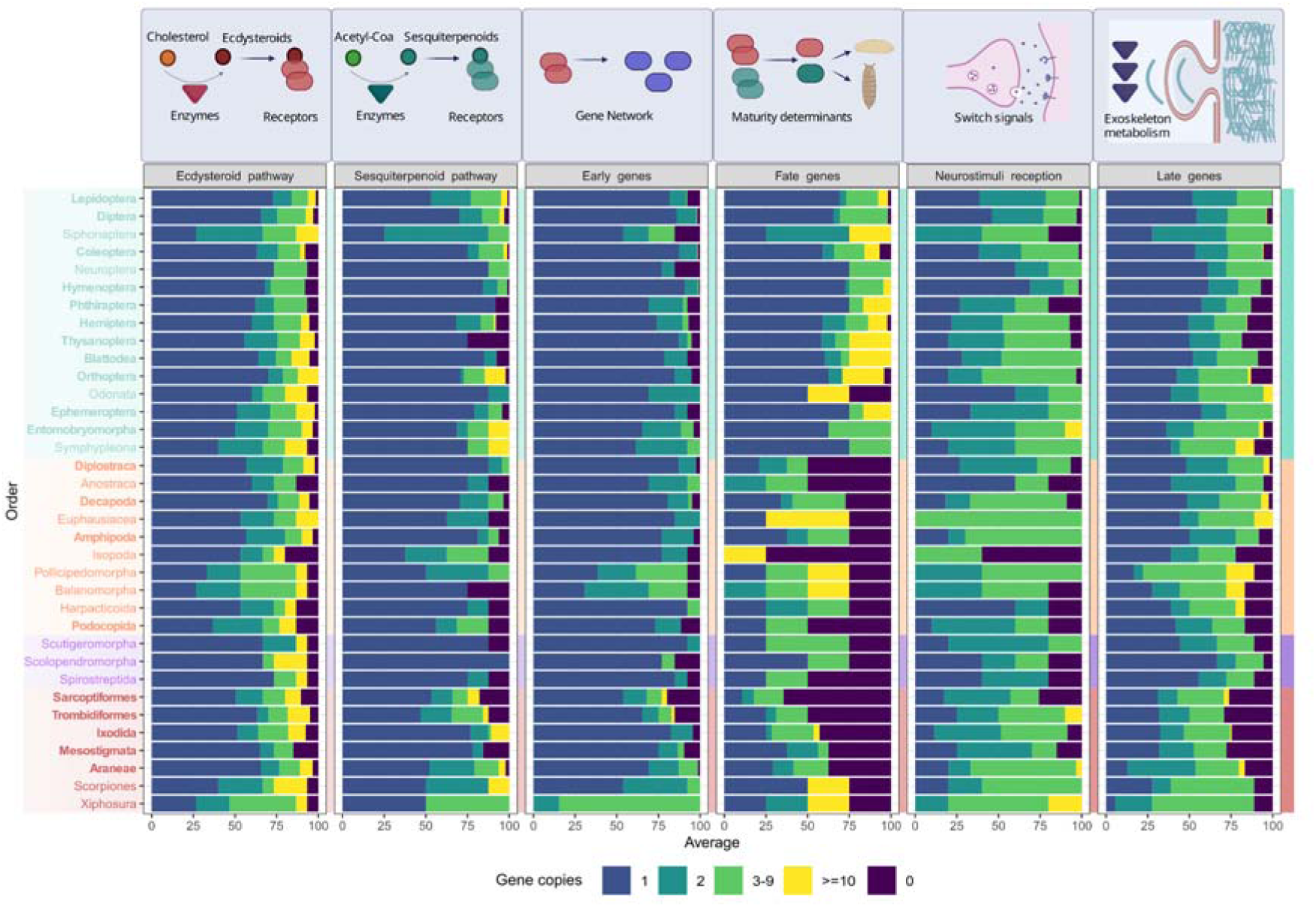
Gene family copy-number profiling of the moulting machinery. Phylogenetic profiling of gene copy numbers of families across the 145-species dataset, partitioned by the 35 represented orders (rows) and six functional modules of the moulting machinery (columns). The cartoons above each functional module summarise the roles they play in moulting processes. The four arthropod subphyla are coloured: Chelicerata (brick red), Myriapoda (lilac purple), Crustacea (peach orange), and Hexapoda (water green). Orders with more than one representative species are shown in bold. For each order, the bars show averaged proportions across the species within each order, counts of genes in orthologous groups (OGs) according to copy number, and count of genes for which no orthologues could be identified. The variability of the individual member gene families of each module is detailed in Supplementary Figures 2-7.

Of the two pathways for hormone synthesis and signalling, the ecdysteroid pathway exhibits an overall lower proportion of gene families with single-copy orthologues across the phylum than the sesquiterpenoid pathway (Figure 2). This is driven mainly by duplications of genes encoding cytochrome enzymes involved in ecdysteroid biosynthesis, as well as transporters for ecdysteroid trafficking, with other members showing more conservative evolutionary histories (Supplementary Figure 2). The cytochrome P450 monooxygenase (CYP) genes include *spook* (*spo*, cyp307a1), *spookier* (*spok*, *cyp307a2*), *phantom* (*phm*, *cyp306a1*), *disembodied* (*dib*, *cyp302a1*), *shadow* (*sad*, *cyp315a1*), *shade* (*shd*, *cyp314a1*), and *cyp18a1* (Pan et al. 2021). With two gene copies in most insects and some crustaceans, the profiling results agree with previous observations indicating that *spook* was present in the arthropod ancestor while its paralogue emerged in the pancrustacean ancestor but not the Mandibulata ancestor (Schumann et al. 2018). The *disembodied*, *shadow*, and *shade* families are generally found as single-copy orthologues, but apparent absences of *disembodied* –apart from in barnacles– are not supported by evidence from the Arthropod P450 Enchiridion (Charamis et al. 2025). In contrast, the family containing *phantom* and *cyp18a1* is highly expanded –most orders have 10 or more genes– and includes the cyp15 epoxidase cytochromes. Co-clustering of *phantom* and *cyp18a1* is consistent with the emergence of *phantom* in the mandibulate ancestor (Schumann et al. 2018), however, the cyp15 members should cluster separately. CYP2 clan P450s that are abundant, especially in non-insect arthropods, so the complexities of lineage-specific expansions of CYP clans mean that homology-based inferences may be unreliable. Hypotheses on gene function thus require biochemical experiments to understand which of these enzymes in each species is responsible for each biosynthetic reaction (Campli et al. 2024).

The families of transporters for ecdysteroid trafficking exhibiting expansions include the ATP-binding cassette (ABC) transporter expressed in trachea (Atet) and the Ecdysone Importer (Ecl). Atet is responsible for loading ecdysone into secretory vesicles for subsequent release into the circulatory system (Pan et al. 2021). It belongs to subfamily G of the ABC transporter superfamily, which has substantially expanded in insects, and has lower gene counts in Mesostigmata species (Denecke et al. 2021). Across the 35-order dataset, large family sizes predominate across the insect orders, in several lineages of Acari, and in two of the three myriapod species (Supplementary Figure 2). The family that includes EcI shows generally modest expansions compared to Atet, with 3-9 genes across most orders, and with the largest counts found in non-Hexapod orders (Supplementary Figure 2). EcI is also known as Organic anion transporting polypeptide 74D (Oatp74D), and mediates ecdysone uptake by target cells in peripheral tissues (Okamoto and Yamanaka 2021; Samantsidis et al. 2022; Wellmeyer et al. 2023). As with the cytochromes, identifying functionally equivalent genes involved in ecdysteroid trafficking from amongst the Atet and EcI family expansions would require experimental investigations.

Amongst other ecdysteroid pathway gene families, we report here, to the best of our knowledge, the first identification of *shroud* (*sro*, also named *non-molting-glossy*) beyond model insects. *Shroud* encodes a short-chain dehydrogenase/reductase that functions in the ‘Black Box’ of the ecdysteroid biosynthesis pathway (Niwa et al. 2010), found across all arthropod orders in the orthology dataset except in Isopoda (Supplementary Figure 2). The sterol regulatory element binding protein (SREBP) is known in insects as an important lipid sensor, while in the shrimp *Litopenaeus vannamei* it is involved in regulating ecdysteroidogenesis (X. Zhang et al. 2019; Jing and Behmer 2020; Zheng et al. 2022). The results extend the breadth of previous observations in insects and a few decapod species to chelicerates and thus to the arthropod ancestor. Additionally, the results indicate that a single gene coding for the membrane steroid binding protein (MSBP) was present in the arthropod ancestor (Supplementary Figure 2). In *D. melanogaster* cell lines, ecdysone sequestration by MSBP suppresses ecdysone signalling (Fujii-Taira et al. 2009). Finally, the Ecdysone Receptor (EcR) has likely duplicated in daphnids from the order Diplostraca, and its partner Ultraspiracle (Usp, also known as Retinoid X Receptor, RXR) is found in two copies in roughly 75% of Araneae, Mesostigmata, Ixodida, and Sarcoptiformes species (Supplementary Figure 2).

The sesquiterpenoid pathway, which produces hormones such as juvenile hormone (JH) and methyl farnesoate (MF), exhibits much higher proportions of single-copy orthologues in most orders compared to the ecdysteroid pathway (Figure 2). The exceptions to this are the JH epoxide hydrolase (JHEH) and JH acid methyl-transferase (JHAMT) families, with multi-copy orthologues in a majority of orders (Supplementary Figure 3). In insects, JHEH and JHAMT share JH acid as a substrate, but they mediate reactions in opposite directions; while JHAMT can catalyse the formation of JH to stimulate JH-induced biological responses, JHEH activity leads to JH diol acid and irreversible inactivation of JH (Qu et al. 2015; Smykal and Dolezel 2023; Campli et al. 2024). Both show expansions mainly in Hexapoda, with more than 50% of the species in multi-species orders having at least two copies. A similar pattern of duplications is seen in Chelicerata (Supplementary Figure 3). In Myriapoda and Crustacea they are mostly found as single-copy orthologues, except for some lineages within Malacostraca, where JHAMT might have been lost and JHEH duplicated.

JH synthesis is considered an insect innovation, so the study of sesquiterpenoid pathway components is biased towards holometabolan models, with genomic analyses limited to modest datasets comprising only a few non-insect representatives (Li et al. 2007; Qu et al. 2015; Sin et al. 2015; Wang et al. 2022; Campli et al. 2024). Yet the profiling results support the presence of all key components of the JH pathway in the arthropod ancestor (Supplementary Figure 3). This is interesting given that most of their characterised roles are based on the study of JH in insects, with little exploration of how they might interact with MF or farnesoic acid (FA) in other arthropods. The presence of these components in the arthropod ancestor suggests they were ancestrally involved in sesquiterpenoid processing, an idea supported by the known ability of methoprene-tolerant (Met) to bind MF (Miyakawa et al. 2013). In insects, JH-mediated signalling FK506-binding protein 39kD (FKBP39) and calponin homology domain protein 64 (Chd64) bind to JH-responsive DNA elements; JH binding protein (JHBP) and JH esterase binding protein (JHEBP) are known for their roles in JH sequestration; Met and its partner Taiman (tai) are known as JH receptors. All of these JH-associated proteins could have ancestrally targeted other hormonal forms or intermediates. While not as widespread or large as for JHEH and JHAMT, duplications are also observed for Met. One duplication has been documented in the fruit fly and tsetse fly with the paralogue germ cell-expressed (*gce*) exhibiting partially overlapping but not fully redundant functions. Two copies have also been found in the silkmoth *Bombyx mori* (Daimon et al. 2015; Jindra et al. 2015). The profiling results extend identified putative *met* paralogues also in Blattodea, Ephemeroptera, Diplostraca, Decapoda, Podocopida, Trombidiformes, and Araneae (Supplementary Figure 3), suggesting that multiple copies for fine regulation of immature phenotype gating might also be found beyond Holometabola.

The early genes, transcription factors activated just after hormone stimulation, are the module with the most single-copy orthologues across the orders (Figure 2). A notable exception is the Xiphosura horseshoe crab, which has multiple copies for each gene family (Supplementary Figure 4). This is based on only a single species, but is consistent with three rounds of whole genome duplications (WGDs) in horseshoe crabs (Nong et al. 2021; Castellano et al. 2025). Interestingly, these genes have been maintained despite apparently strong constraints to avoid duplication across the rest of the phylum, which is consistent with a general pattern of constrained genes being retained in duplicate after WGDs (Makino and McLysaght 2010; Singh et al. 2012; Roux et al. 2017; Aase-Remedios et al. 2025). In the orthology dataset the Ecdysone-inducible genes 75 (*E75*) and 78 (*E78*) are clustered in the same OG, however these were likely both present in the ecdysozoan ancestor (Schumann et al. 2018). In this otherwise well-maintained set of gene families a plausible loss could have occurred of the gene encoding Hormone Receptor 96 (HR96), a nuclear receptor that is inducible by 20-hydroxyecdysone (20E) and binds cholesterol (Fisk and Thummel 1995; Jing and Behmer 2020). It was previously reported missing from the aphid *Acyrthosiphon pisum* (Bonneton and Laudet 2012), and appears to be absent from more than half the Hemiptera species in the orthology dataset, as well as in all of the mites of the order Mesostigmata (Supplementary Figure 4). Together with the potential losses of the neverland (nvd) enzyme for cholesterol processing and the Atet transporter, these results favour the hypothesis that novel pathways might have emerged in mites in relation to parasitic lifestyles and synchronisation to host metabolism (Cabrera et al. 2015; Aurori et al. 2021).

The small set of fate genes is characterised by a notable pattern of missing genes beyond hexapods (Figure 2). This is mainly due to the presumed emergence of the larval determinant transcription factor, chronologically inappropriate morphogenesis (*chinmo*), in the hexapod ancestor. Chinmo is essential for maintaining the nymphal phenotype in *O. fasciatus* and the larva in *Drosophila*, in opposition to two other fate genes, *broad* and *ecdysone-inducible gene 93* (*E93*) (Truman and Riddiford 2022; Nagata and Suzuki 2025). Another immaturity determinant, Kruppel homolog 1 (*Kr-h1*), appears to have been repeatedly lost in chelicerates, since no orthologues were detected in at least 50% of species from Araneae and Acari orders, and in any of Trombidiformes, while several other chelicerates do have identifiable orthologues (Supplementary Figure 5). Sarcoptiformes appears to have the most incomplete set, with 86% of the surveyed species also missing the adulthood specifier *E93*. The *broad* gene family exhibits a complex history, with many BTB/POZ transcription factors clustering together resulting in more than three copies throughout Arthropoda (Supplementary Figure 5).

The neurostimuli module exhibits the lowest proportion of single-copy orthologues, with multiple copies of the genes coding for neuropeptide receptors triggering moulting in all four subphyla. Between 60% and 80% of pathway components are duplicated, often with multiple copies (Figure 2). This is particularly striking for the eclosion hormone receptor (EHR) and Rickets (rk) gene families (Supplementary Figure 6), suggesting the presence of multiple copies in the Arthropoda ancestor. In insects, ecdysis triggering hormone receptor (ETHR) is known to have multiple isoforms that might have acquired different specialisations (Campli et al. 2024), thus isoform sequence evolution and lack of accuracy in isoform annotation might inflate the number of putative paralogues (Supplementary Figure 6). Although the ecdysis neuromotor signal reception components are complete in other chelicerates, they are missing from more than 50% of Sarcoptiformes and Mesostigmata species (Supplementary Figure 6). Previous investigations reported conflicting results for these components, especially for the existence of the crustacean cardioactive peptide receptor (CCAP-R) signal (Campli et al. 2024).

The late genes exhibit elevated proportions of duplications, similar to the ecdysteroid pathway (Figure 2). This module comprises genes involved in exoskeleton assembly such as the enzymatic effectors necessary for synthesis, remodelling, spatial arrangement, and cross-linking the building blocks of chitin and cuticle proteins (CPs). The superfamily of CPs is not included in the profiling analysis due to its large and complex nature (Willis 2010) and its largely structural role in moulting processes. Across the multi-species orders in the dataset, from 31% (Hymenoptera) to 70% (Araneae) of the late effector genes were detected at least in double-copy (Figure 2). Amongst the chitinases, Cht2, Cht4, Cht8, and Cht9 cluster together in a large multi-copy group spanning all orders, in contrast to the most conservative Cht7 that is almost universally single-copy (Supplementary Figure 7). A key effector step is cuticle hardening, achieved in insects mainly by cross-linking CPs with oxidised catechol derivatives, as in the pathway for melanin synthesis (Campli et al. 2024). This oxidation is mediated by the laccase enzyme straw (stw), paralogue of the multicopper oxidase1 (mco1), from a large superfamily of multicopper oxidases (Asano et al. 2019; Janusz et al. 2020). The profiling analyses suggest the absence of *stw* or *mco1* orthologues in chelicerates (Supplementary Figure 7), although related multicopper oxidases may be present. No studies to-date have fully characterised the mechanisms for cuticle hardening in chelicerates, in fact discovery of melanin itself is relatively recent (Hsiung et al. 2015), and the absence of *mco* has been preliminarily explored only in four chelicerates (Asano et al. 2019). Therefore, in lineages where post-ecdysis hardening of the exoskeleton is poorly understood, mechanisms based on under-explored cross-linking substrates such as resilin are of particular interest. Resilin is a matrix protein that cross polymerises by tyrosine-mediated bonds and thus gives elastic resistance to joints, tendons, or wings, in locusts, dragonflies, and flies. Surprisingly, it has been found at high concentration in a copepod ‘tooth’ specialised to break hard diatom shells (Michels et al. 2012; Lerch et al. 2020; Lerch et al. 2022). The profiling results reveal more than three copies of resilin-like genes across different arthropod orders and at least ten copies in 67% of Araneae species and in some crustaceans, a few Podocopida, a copepod, and a barnacle (Supplementary Figure 7), indicating a putative role for diverse resilins in many arthropods.

### Moulting toolkit insights from three new myriapod genomes

Mandibulata comprises a monophyletic clade including Pancrustacea and Myriapoda (Chipman 2024). Genomic sampling of Myriapoda remains limited relative to the other subphyla, therefore, sampling additional myriapod species helps to build a better characterisation of this group, and to investigate changes occurring in the mandibulate ancestor. Genome assemblies were produced from Pacific Biosciences HiFi sequencing reads with estimated coverage of 40x to 47x resulting in genomes of 1.5 Gbp (600 contigs), 2.2 Gbp (344 contigs), and 1.5 Gbp (3’887 contigs) for *S. cingulata*, *S. coleoptrata*, and *A. syriacus*, respectively (see Materials and Methods). Genome annotation employed RNA sequencing data to predict gene models for each assembly, resulting in sets of 17’995, 18’157, and 11’437 protein-coding genes for *S. cingulata*, *S. coleoptrata*, and *A. syriacus*, respectively, with completeness scores of between 95% and 98%.

Inspection of the annotated genomes of the two centipedes and the millipede revealed several potentially relevant lineage-specific peculiarities in hormone synthesis and response, with an otherwise strong conservation of the rest of the arthropod moulting toolkit (Figure 3). In the three species, 41% of gene families were recovered as single-copy orthologues in both Chilopoda and Diplopoda, and 13% as duplicated with at least two copies in both classes. Duplication and loss events in myriapods generally reflect the observed arthropod-wide patterns. These include a propensity for duplications of the *phm* and *cyp15* cytochromes (Supplementary Figure 2), as well as frequent detection of multi-copy families for EcI (Supplementary Figure 2), Broad (Supplementary Figure 5), EHR (Supplementary Figure 6), chitinases and chitin deacetylases (Supplementary Figure 7). The analyses of the myriapod genomes further indicated that the duplication of spo-like *cyp307* genes does not extend beyond Pancrustacea (Supplementary Figure 2), and that *chinmo* was not present in the mandibulate ancestor (Supplementary Figure 5).

**Figure 3.**
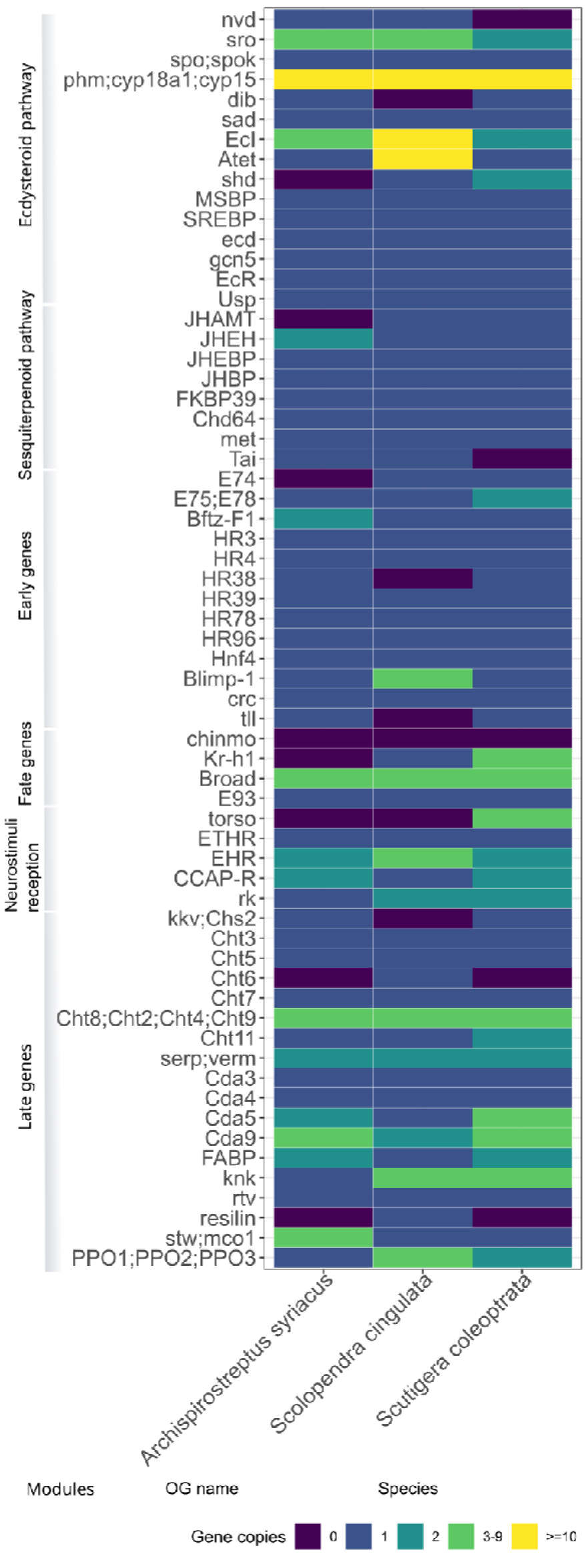
Moulting gene family copy-number profiling in myriapods. Phylogenetic profiling of gene copy numbers of families involved in moulting across the three myriapod species sequenced and annotated in this study, namely the centipedes *S. cingulata* and *S. coleoptrata*, and the millipede *A. syriacus*. Orthologues are partitioned into six functional modules of the moulting machinery (named along the side of the figure). Cells show the counts of genes in orthologous groups (OGs) classified as single-copy (blue) or multi-copy (green, yellow) orthologues, or for which no orthologues could be identified (deep purple). Gains and losses of genes involved in moulting are evident across all modules and all three species: the ecdysteroid pathway and late genes show elevated proportions of duplications/gains, while in contrast the sesquiterpenoid pathway and early genes modules exhibit mainly single-copy orthologues. Full gene names are provided in Supplementary Figures 2-7.

The absence of a JHAMT orthologue from the millepede *A. syriacus* (Figure 3) is congruent with the hypothesis that millipedes, unlike centipedes, use farnesoic acid (FA) as their main moulting hormone, since they are not able to further transform it into methyl farnesoate (So et al. 2022). It is worth noting that millipedes might also have alternative strategies for ecdysteroid stimulation, suggested by the apparent absence of a *shd* orthologue and thus, the lack of the last step for catalysis of ecdysone into 20E, which is generally accepted as its biologically active derivative (Campli et al. 2024). Beyond the variations in hormone synthesis pathways, the main response mechanisms to hormonal signalling might be even further remodelled. *A. syriacus* also appears to lack orthologues of the transcription factor encoding *Ecdysone-inducible protein 74* (E74) gene, which is activated early in the moulting cascade by 20E stimulation even in chelicerates. Additionally, *Kr-h1*, an important JH transducer, is also putatively missing from the millipede genome. The responsiveness of *Kr-h1* to sesquiterpenoids has only been studied in Hexapoda, but the gene is also detected in some chelicerates (Campli et al. 2024). The future availability of additional annotated genomes for millipedes will help to clarify these initial observations from *A. syriacus*.

The centipede *S. coleoptrata* displays another putative metabolic difference, with the putative loss of the *nvd* orthologue (Figure 3). Since *nvd* encodes the enzyme catalysing the first step in the biosynthetic pathway beginning with cholesterol, its absence suggests the possibility of novel solutions for cholesterol intake. If confirmed, this would be specific to the subclass Notostigmophora, since *nvd* is found in the genomes of the centipedes *S. cingulata* and *Strigamia maritima*, which both belong to the sister subclass Pleurostigmophora (Schumann et al. 2018). Additionally, four putative copies of the torso receptor were found in *S. coleoptrata*, but no homologues were detected in *S. cingulata* or *A. syriacus*, as previously observed in the centipede *S. maritima* (Chipman et al. 2014). Its absence might indicate deep differences not only for the regulation of the moulting process, but also for embryonic patterning, since torso in insects binds both the paralogues prothoracicotropic hormone and the trunk ligand as part of the terminal patterning pathway (Rewitz et al. 2009; Duncan et al. 2013; Weisbrod et al. 2013; de Oliveira et al. 2019; Campli et al. 2024).

### Ancestral state reconstructions map gene gain and loss events and protein domain rearrangements

While copy-number profiling characterises evolutionary dynamics from the perspective of extant species’ gene repertoires, understanding where and when key changes may have occurred requires reconstructing gene evolutionary histories. Employing ancestral state reconstructions of gene counts and ancestral domain architectures, the orthology dataset was analysed to trace gene family expansions and contractions across the arthropod phylogeny, and to infer domain architecture rearrangements that occurred throughout the evolution of Mandibulata.

Mapping gene gains and losses across Arthropoda was performed on a subset of 97 species after reducing the number of insects and selecting one representative for genera that included more than one species in the 145-species dataset. In total, the analysed gene families involved in moulting were predicted to have undergone 1’114 expansions and 964 contractions. Comparing across all gene families of the same age, 16.7% of ancestral nodes showed a greater proportion of expansions amongst families involved in moulting processes (Figure 4A). That is, the odds of experiencing expansions rather than no-change or contraction events were greater for moulting-related genes than other gene families at these 16 ancestral nodes. This expansion excess amongst moulting-related gene families is evident at the LCA of each subphylum except Myriapoda, and occurs at other ancient nodes such as the Collembola LCA, the Acariformes LCA, and the ancestor of Araneae and Scorpiones (Arachnopulmonata). These observations point towards gene duplications playing a role in shaping the evolutionary trajectories of members of the moulting toolkit during the early diversification of the main Chelicerata lineages and the early pancrustacean radiations. No expansion excess was detected for Myriapoda, which could suggest a more stable evolutionary history in this subphylum, however, better taxonomic representation is required to confirm this trend. Notably, across the represented eumetabolous insects (Holometabola and Paraneoptera) none of the 35 ancestral nodes show any significant expansion excess (Figure 4A), suggesting that gene duplications of the moulting machinery were limited following the transition towards insect metamorphosis.

**Figure 4.**
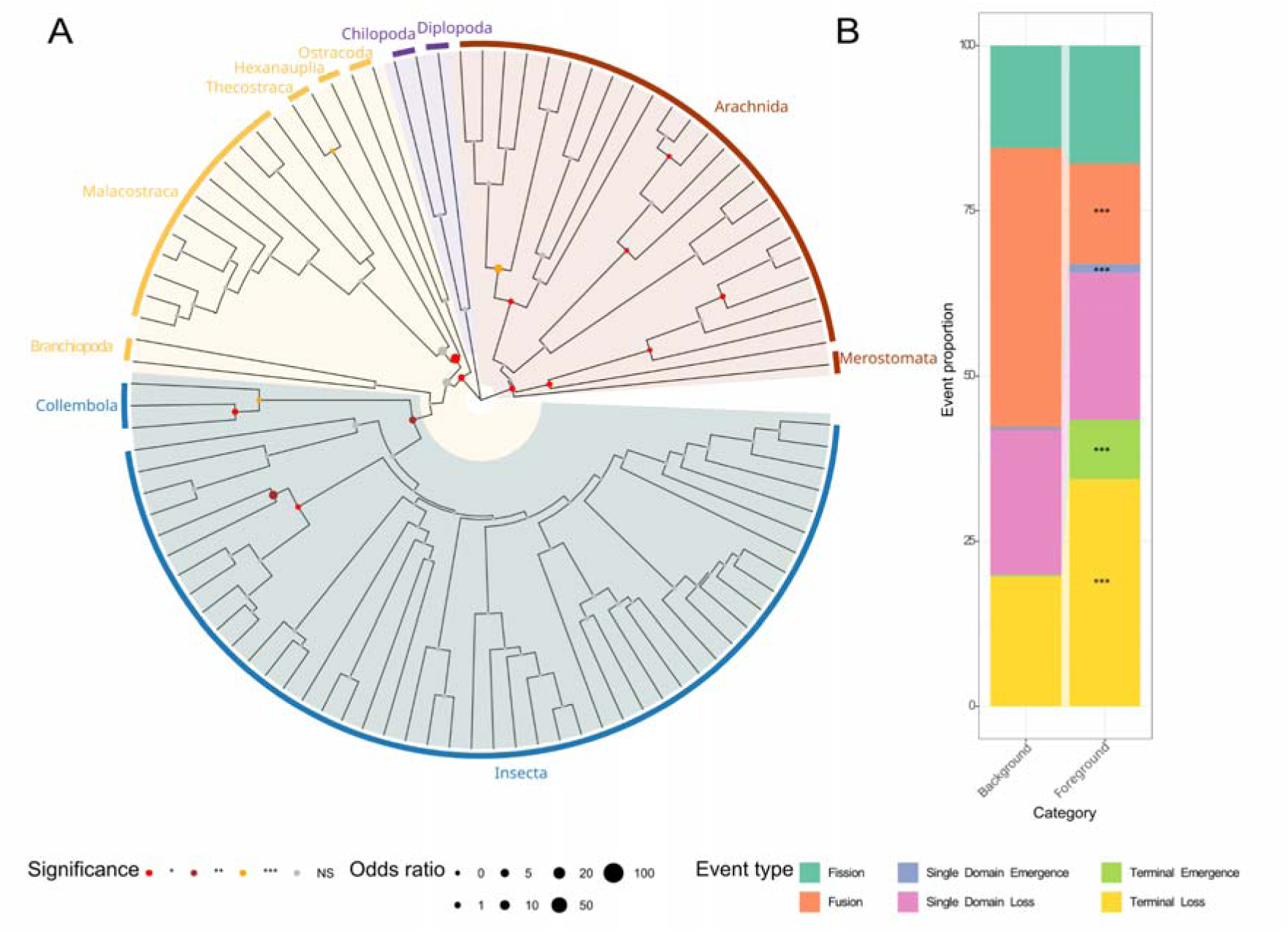
Gene family and protein domain architecture dynamics of moulting gene families. **(A)** Ancestral state count reconstructions of gene family dynamics of moulting gene families. Gene counts at internal nodes are reconstructed from the observed gene copy numbers of the leaves to infer changes in family sizes at each ancestral node. The phylogeny shows the four arthropod subphyla highlighted: Chelicerata (pink), Myriapoda (purple), Crustacea (yellow), and Hexapoda (green). Nodes are annotated with a solid circle reporting the odds ratio (size) and strength of corrected p-values (colours, p.adj < 0.05 *, p.adj < 0.005 **, p.adj < 0.0005 ***, NS not significant) of excess gene family expansions detected amongst moulting-related gene families compared to other gene families (Fisher’s exact test). **(B)** Proportions of identified protein domain rearrangements explained with exact and non-ambiguous solutions for classification as fissions, fusions, single domain emergence or loss events, and terminal emergence or loss events. Rearrangement events in domain architectures were inferred by ancestral reconstructions comparing the moulting gene families set (foreground) and the background gene set of all families as old as the Arthropoda LCA (asterisks indicate Fisher’s exact test p.adj < 0.005 **, p.adj < 0.0005 ***).

Domain architecture analysis of proteins from the 145-species Arthropoda LCA orthology dataset annotated with PfamScan catalogued an expected majority of 90% stable and maintained arrangements (Supplementary Figure 8). Amongst the identified rearrangements, 459 (44%) can be explained with exact and non-ambiguous solutions for classification as fissions, fusions, single domain emergence or loss events, and terminal emergence or loss events (Figure 4B). Compared to the events resolved across all other genes families, the moulting gene set experienced a similar proportion of fission events alongside a greatly reduced proportion of domain fusion events (15.25%, odds ratio = 0.3), *i.e.* a tendency to avoid creating new combinations of domains throughout their evolutionary histories. While the proportions of single domain losses are similar, the emergence of single domains is significantly higher amongst the moulting gene set members (1.31%, odds ratio = 4.6), albeit representing a small fraction of the total events. Terminal loss and emergence events are both significantly enriched for the moulting machinery with a notably large increase in the proportion of terminal emergences (terminal losses: 34.42%, odds ratio = 1.6; terminal emergences: 8.93%, odds ratio = 25). The over-representation of gains or losses of domains at either end of an ancestral domain architecture amongst the moulting gene set members suggests a flexibility to elaborate on a conserved core, with likely functional consequences in fine-tuning conserved functions across different arthropod lineages.

### Reconciliations pinpoint ancient duplications and genome screening corroborates lineage-wide losses

The copy-number profiling and ancestral state reconstructions highlight several components that either retained ancient gene duplicates or experienced lineage-wide losses of otherwise conserved gene families. These are of particular interest as they represent changes that could have substantially impacted the functional modules governing the moulting process. Potential ancient duplications were therefore carefully examined employing gene-tree species-tree reconciliation approaches, where feasible, to complement and support the inferences from ancestral state reconstructions. It is becoming increasingly evident that in addition to gene duplications, gene losses can represent a mechanism that generates genetic diversity driving lineage-specific evolution (Albalat and Cañestro 2016; Fernández and Gabaldón 2020). Taking advantage of publicly available unannotated assemblies to predict specific sets of genes and confirm lineage-specific events fully exploits the increasing amount of data from understudied species (Langschied et al. 2024). Thus, to corroborate potential losses, gene searches were performed using unannotated genomes for species from clades where orthologues appeared to be missing, and for which at least several assemblies were available for species from the clade. These investigations focused on the chitin deacetylase (CDA) genes *serpentine* (*serp*), *vermiform* (*verm*), and *Chitin deacetylase-like 4* (*Cda4*), the chitin synthase (CHS) genes *krotzkopf verkehrt* (*kkv*) and *Chitin synthase 2* (*Chs2*), the cuticle organising *knickkopf* (*knk*) gene, a set of genes encoding prophenoloxidases, the larval stage maintaining *chinmo* gene, the adulthood determinant *E93* gene, and the Rieske-type oxygenase encoded by the *neverland* (*nvd*) gene (see Materials and Methods).

The chitin deacetylases and the chitin synthases are key effectors involved in chitin fibre formation that form a major structural component of arthropod exoskeletons. The cuticle is a complex multi-layer structure where proper orientation of individual chitin fibres is fundamental to achieve a helicoidal super-organisation, mainly responsible for the overall exoskeleton resistance (Roer et al. 2015; Politi et al. 2021). In insects, the exoskeleton is built by formation of the apolysial space from its innermost edge, the endocuticle (Moussian 2010). There, a monolayer of epidermal cells produces chitin, which is then transported to the apical membrane, secreted, and assembled into the extracellular matrix. Chitin is made either by *de novo* synthesis from trehalose, mediated by chitin synthases 1 and 2 (*Chs*, *Chs1/kkv* and *Chs2*) or by recycling of the former fibres into chitin monomers, digested by Chitinases (*Chts*) and deacetylated by Chitin deacetylases (CDAs, *CDA1/serp* and *CDA2/verm*). After extrusion, chitin binds cuticle organisation and maturation effectors such as the GPI-anchored protein Knickkopf (*knk*) and arranges into organised sheets.

Reconciliation analysis of these key factors for exoskeleton remodelling identifies duplications at the LCAs of Mandibulata, Araneae, Chelicerata, and Pancrustacea (Figure 5A). The inclusion of myriapods in the dataset pinpoints the duplication of the single ancestral *CDA1/serp-CDA2/verm* chitin deacetylase gene to the Mandibulata LCA, with both copies well-maintained across the descendant lineages (Supplementary Figure 9). An independent duplication of *CDA1/serp-CDA2/verm* is also evident at the LCA of spiders, with two copies maintained in each of the five species included in the analysis. The spider LCA is also the location of the duplication of *Cda4*, another chitin deacetylase gene gain which is also maintained across the five species (Supplementary Figure 10). Both the chitin synthases *kkv* and *Chs2* are widely present across Pancrustacea species, and chelicerates also generally harbour two copies, it could therefore be assumed that both these genes were present in the Arthropoda LCA. However, the myriapods have only one copy and the reconciliation instead favours the evolutionary scenario of two independent ancestral duplications, one at the Chelicerata LCA and the other at the Pancrustacea LCA (Supplementary Figure 11). In the case of the *knk* gene family the orthology dataset identifies more than two copies in all but two of the 35 orders, suggesting the possibility of multiple ancestral duplications. The reconciliation analysis instead confirms the presence of three copper-dependent monooxygenase homologues in the arthropod ancestor, where the family is characterised by differential gene losses rather than ancient gains (Supplementary Figure 12).

**Figure 5.**
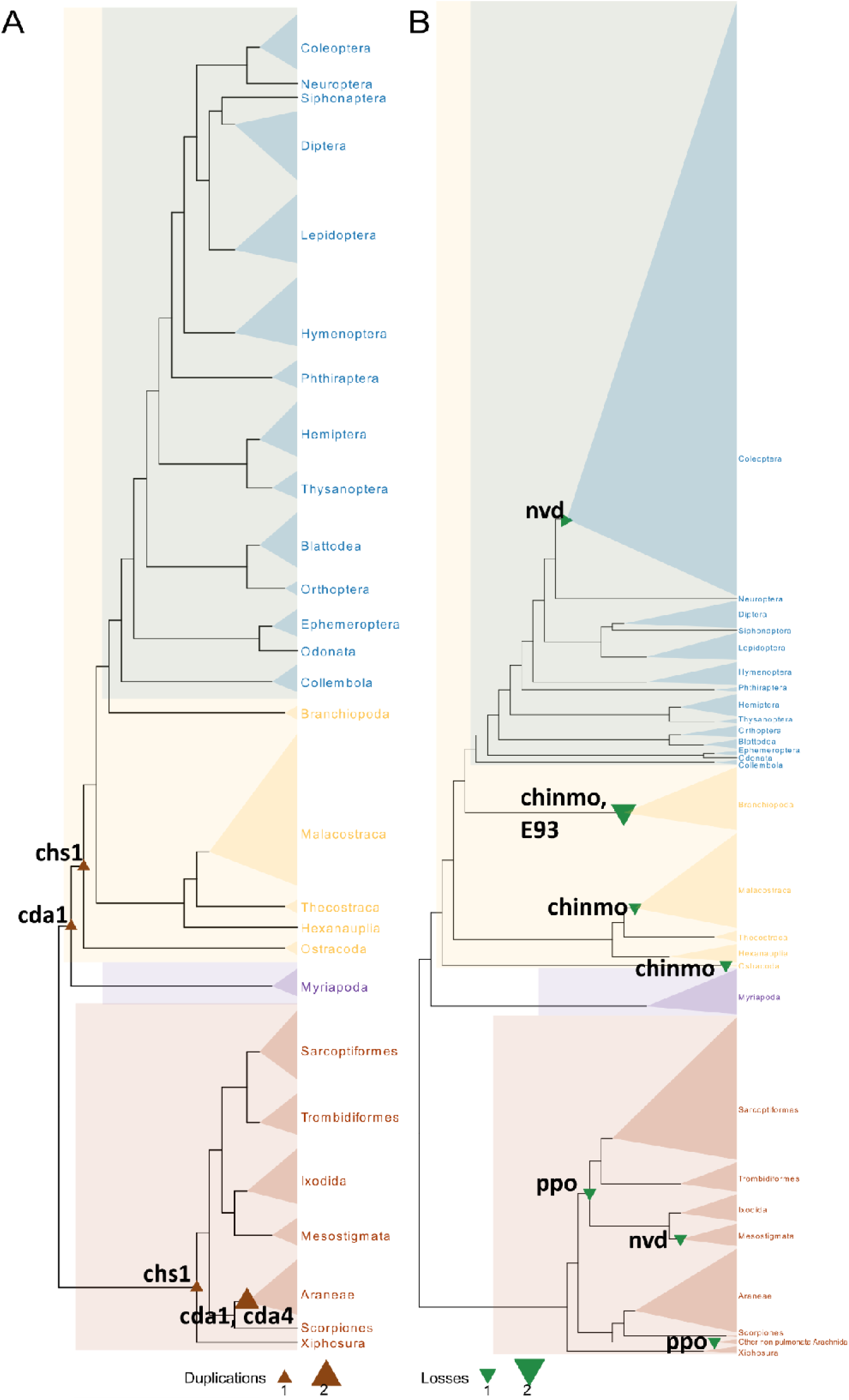
Confirming ancient gene duplications and lineage-wide losses. Candidates identified from gene copy-number profiling as representing likely ancient duplications or lineage-wide losses were further investigated to confirm events and map them on the arthropod phylogeny. Brown and green triangles pinpoint duplication and loss events, respectively, on the phylogenies for the named gene families. **(A)** Summary of ancient and maintained duplications of key factors for exoskeleton remodelling confirmed by gene-tree species-tree reconciliations. The collapsed nodes are depicted with triangles proportional in size to the number of species used for the reconciliation analysis. **(B)** Summary of lineage-wide losses confirmed by genome searches. The collapsed nodes are depicted with triangles proportional in size to the number of genomes searched, e.g. Coleoptera (n=251), Sarcoptiformes (n=57), Branchiopoda (n=22), Myriapoda (n=18), Mesostigmata (n=7).

The Mandibulata LCA gene gain event leading to the *CDA1/serp*-*CDA2/verm* pair of chitin deacetylase genes represents a likely case of functional specialisation after duplication. Both the CDA1/serp and the CDA2/verm proteins, as well as the proteins from the chelicerates, display a PFAM domain architecture that starts with an N-terminal chitin binding Peritrophin-A domain (PF01607) followed by a low-density lipoprotein receptor domain class A (PF00057). However, a C-terminal NodB polysaccharide deacetylase domain (PF01522) is only identified in the chelicerate and CDA1/serp proteins and the myriapod CDA2/verm proteins, despite the pancrustacean CDA2/verm proteins having similar protein lengths (Supplementary Figure 13). Conserved Domain Database (CDD) and SUPERFAMILY domain annotations reveal a CDA-like domain spanning this region even in the pancrustacean CDA2/verm proteins, indicating that it is still generally recognisable while not specifically identified by the PF01522 profile. The CDD annotations reveal a key difference: within the CDA-like domain of the chelicerate and CDA1/serp proteins and the myriapod CDA2/verm proteins is a Zn-binding site with a conserved Asp-His-His triad, but this is not identified in the pancrustacean CDA2/verm proteins. This suggests that after the duplication event in the Mandibulata LCA, the capacity to bind Zn was maintained in both copies in myriapods but lost in the CDA2/verm copy in the pancrustacean ancestor. CDA2/verm catalytic activity has been challenged in a few species, such as *D. melanogaster*, *L. migratoria*, and the prawn *P. monodon*, where a role in mediating chitin fiber organisation has been suggested instead (Dixit et al. 2008; Sarmiento et al. 2016; M. Zhang et al. 2019; Zhang et al. 2021). Therefore, CDA2/verm might have functionally diverged from CDA1/serp and additional experimental evidence may further support an evolutionary scenario of duplication followed by functional specialisation. The pair of *CDA1/serp*-*CDA2/verm* genes display a highly constrained genomic organisation as their tandem position as neighbouring genes is conserved in 65% of pancrustaceans with chromosome-level genome assemblies (and as near-neighbours allowing +/- 5 genes in 88%). Thus despite their putative functional divergence, conserved genomic proximity suggests that coordinated transcriptional control has constrained their evolutionary histories.

Amongst the losses highlighted by the gene copy-number profiling, the pro-phenoloxidases (PPOs) appeared to be absent from several crustacean orders as well as from the arachnid orders Acariformes and Sarcoptiformes (Supplementary Figure 7). PPOs encode the precursor zymogens that are activated via proteolytic cleavage to form the phenoloxidases that catalyse catechol oxidation as part of sclerotization and melanisation mediating cuticle hardening. These enzymes exhibit a complex evolutionary history and are related to the oxygen-carrying hemocyanins and the hexamerin storage proteins of insects and crustaceans (Burmester 2002; Rehm et al. 2012; Scherbaum et al. 2018). Loss patterns across Crustacea suggest multiple independent losses in Podocopida (ostracods), Harpacticoida (copepods), and the Pollicipedomorpha-Balanomorpha (barnacles) ancestor, however genomic sampling of these taxa remains too limited to test for lineage-wide losses. The arachnids, however, are better represented with available genome assemblies, and no PPO-like or hemocyanin-like genes could be recovered in any tick or mite species nor in any other arachnid (83 species) other than for Araneae and Scorpiones. Therefore, across Chelicerata, while Tetrapulmonata species and horseshoe crabs might potentially rely on phenoloxidase activity, others may employ mechanisms different from catechol oxidation for mediating cuticle hardening.

In a few model insects, there is recent evidence for an important role of *chinmo* as a gatekeeper of the immature phenotype (Campli et al. 2024). While *chinmo* was identified in the orthology dataset in all hexapod orders except Odonata (Supplementary Figure 5), a putative orthologue was identified in only a single non-hexapod species, the copepod *T. californicus*. Using the *D. melanogaster* protein sequence as the search seed failed to identify any candidate orthologues in non-hexapod genomes (Crustacea: n=63, Myriapoda: n=18, Chelicerata: n=118). However, chinmo includes stretches of intrinsically disordered regions that can hinder effective sequence searches. Using the *T. californicus* sequence to search non-hexapod genomes identified putative orthologues, that form best reciprocal hits to the query sequence and can be annotated with both the BTB/POZ and double zinc finger domains, in seven out of eight copepods. This tentatively dates the emergence of *chinmo* to the copepod-hexapod LCA, with subsequent losses in other crustaceans. Whether these genes play similar gatekeeper roles in copepods remains to be determined.

While the DNA-binding E93 protein has been established as a gatekeeper of metamorphosis and adulthood fate in Holometabola (Ureña et al. 2016), phylogenetic profiling indicates its near-universal single-copy presence across arthropod orders (Supplementary Figure 5). However, there appear to have been secondary losses in species characterised by direct development from the class Branchiopoda and the order Sarcoptiformes. Genome searches using E93 from *D. melanogaster* or orthologues from the decapod *P. monodon* and the amphipod *H. azteca* revealed no candidate *E93* orthologues in any member of the branchiopod crustacean orders Anostraca, Diplostraca, or Notostraca (22 species). For the Sarcoptiformes, only one out of the seven species in the orthology dataset harboured a putative *E93* orthologue. These widespread losses were further supported by genome searches with sequences from the spider *P. tepidariorum* and the mite *T. urticae* that recovered putative *E93* orthologues in only 28% of the examined Sarcoptiformes (16/57 species). These repeated independent losses of *E93* across different subphyla may have played a role in the regulatory rewiring of toolkit components controlling the progression of development through progressive moults.

The Rieske monooxygenase encoded by the gene *neverland* is near-universally present across the investigated orders, yet no copies were found for Coleoptera or Mesotigmata, which are represented by 11 and four species, respectively (Supplementary Figure 2). Extensive searches of available genomes failed to reliably predict copies of *nvd* in any of the 251 assemblies for Coleoptera, spanning 41 different taxonomic families, or any of the seven Mesostigmata species with available assemblies. For beetles this presents a strong case for the order-wide loss of *nvd* in Coleoptera, while within Mesostigmata additional representative species are needed to confirm this potential loss and if confirmed, to better understand its extent. The genome searches of the seven Mesostigmata species using the *nvd* protein sequences from the spider *P. tepidarioum* and the mite *T. urticae* also returned no reliable gene predictions.

While the orthology dataset identified no beetle orthologues, the genome searches using *D. melanogaster nvd* as the search-seed returned three potential hits in *Anisosticta novemdecimpunctata* (Coccinellidae, ladybirds), *Liopterus haemorrhoidalis* (Dytiscidae, diving beetles), and *Sceptobius lativentris* (Staphylidae, rove beetles). Using the *nvd* gene from *A. mellifera* (Hymenoptera) as the search-seed identified two hits within Chrysomelidae (leaf beetles), in *Octodonta nipae* and *Galerucella tenella*. The predictions from the assemblies of *A. novemdecimpunctata* and *L. haemorrhoidalis* had low sequence similarity to any arthropod protein and showed no domain annotations expected of *nvd* orthologues. The candidate *S. lativentris nvd* gene is found on a 1258 bp contig and the protein exhibits the characteristic Rieske domain, however, the sequence is almost identical to a protein from the Sarcoptiformes mite, *Tyrophagus putrescentiae* (99%). Originally thought to be mycetophagous, this mite feeds on insects including coleopterans such as the tobacco beetle *Lasioderma serricorne* (Papadopoulou 2006), suggesting that mite tissue may have been present in the sample used to sequence the *S. lativentris* genome. The *O. nipae* candidate also exhibits the expected domains, however, it shows greatest sequence identity (>50%) with nvd proteins from Sarcoptiformes mites. The candidate *G. tenella nvd* gene is found on an 88 Kbp contig and the protein exhibits the characteristic Rieske domain, however, it has greatest sequence identity with proteins from the endoparasitoid wasps *Microctonus hyperodae* (66%) and *M. aethiopoides* (59%). Endoparasitoids of the genus *Microctonus* are known to target beetle hosts from the Chrysomelidae family such as cabbage stem flea beetle, *Psylliodes chrysocephala* (Jordan et al. 2020), suggesting that the sample used to sequence the *G. tenella* genome may have been harbouring wasp eggs or larvae.

The low-identity hits without Rieske domains are likely spurious predictions and the other *nvd* gene predictions probably represent sample contaminations with plausible biological explanations. Therefore, considering the extensive genome sampling across Coleoptera, the evidence suggests that the beetle ancestor likely lost the *nvd* gene encoding the key ecdysteroid pathway Rieske monooxygenase. Based on current biochemical knowledge, loss of the enzyme probably also indicates loss of the ability to transform cholesterol into 7-dehydrocholesterol. No other protein capable of catalysing this reaction (EC:1.14.19.21) was found in beetles, when querying the ENZYME database (Release 24/07/2024) and UniProtKB, even including those with an unreviewed status (Bairoch 2000). Furthermore, no enzyme was identified in beetles when considering the reverse catalytic reaction known from the mammalian DHCR7 reductase (EC:1.3.1.21), which mediates the last step of cholesterol synthesis. Similarly for Mesostigmata, searching for proteins putatively capable of catalysing these reactions returned no candidates for any hypothetical functional replacements. Based on studies in model insects, nvd is localised in the endoplasmic reticulum and its catalytic activity depends on the Rieske [2Fe-2S] motif within the Rieske domain (Interpro entry IPR017941) (Yoshiyama-Yanagawa et al. 2011; Pan et al. 2021). Beetles do possess other genes encoding proteins with this key domain, such as a Cytochrome b-c1 complex subunit located in the mitochondrial inner membrane mediating electron transport (EC:7.1.1.8), and an apoptosis-inducing factor 3 protein with a cytoplasmic localisation. However, the 3-ketosteroid-9-alpha-monooxygenase-like C-terminal domain (Interpro IPR045605) that follows the Rieske domain of the nvd protein, is not detectable in any beetle protein-based searches of the Interpro and UniProt databases.

### Cuticle protein repertoire dynamics vary across arthropod subphyla

The exoskeleton represents a hallmark structure essential to all arthropods that provides physical support and resistance to mechanical and chemical stress, with variations in the structural arrangement of layers and different chemical compositions (Roer et al. 2015; Hensel et al. 2016; Politi et al. 2021; Bello et al. 2022). Exoskeleton assembly and remodelling relies on the building blocks of chitin polymers and CPs, where the superfamily of CP genes is known for its large size and complex evolutionary history (Willis 2010). Therefore, CP gene repertoires were examined by identifying canonical CPs from the CPR family within the annotated proteome of each species based on the presence of their representative Rebers and Riddiford (RR) chitin binding domain (Pfam domain PF00379) (Moussian 2010; Willis 2010; Roer et al. 2015; Mrak et al. 2017). As a key component of the biophysical interface between the arthropod and its immediate environment, the physico-chemical properties (Osorio et al. 2015) of CP repertoire amino acid sequences were also characterised to investigate differences across subphyla.

The species with the smallest CP repertoires included the isopod *A. nasatum*, and the Acariformes *S. scabiei, H. destructor, O. nova, D. farinae, B. tropicalis*, and *D. pteronyssinus*, with 19 to 25 genes, while the largest by far were the three *Penaeus* crustaceans, with more than 600 genes. Comparing across subphyla shows that crustaceans harbour, on average, the largest repertoire of CPs, with a median of 212 CPs (Figure 6A). Median counts for Hexapoda (87), Myriapoda (34), and Chelicerata (70) are all much lower, and they show less within-subphylum variation than the crustaceans. To examine whether larger repertoires necessarily mean more diverse repertoires, proteins from each species with sequence similarities of 80% or more were clustered. The ratio of clusters to counts is a proxy for CP repertoire redundancy, with lower values indicating the presence of more sets of similar CPs. This shows that despite the large counts in many crustaceans, these CPs are often similar in sequence and therefore do not represent more diverse repertoires (Figure 6B).

**Figure 6.**
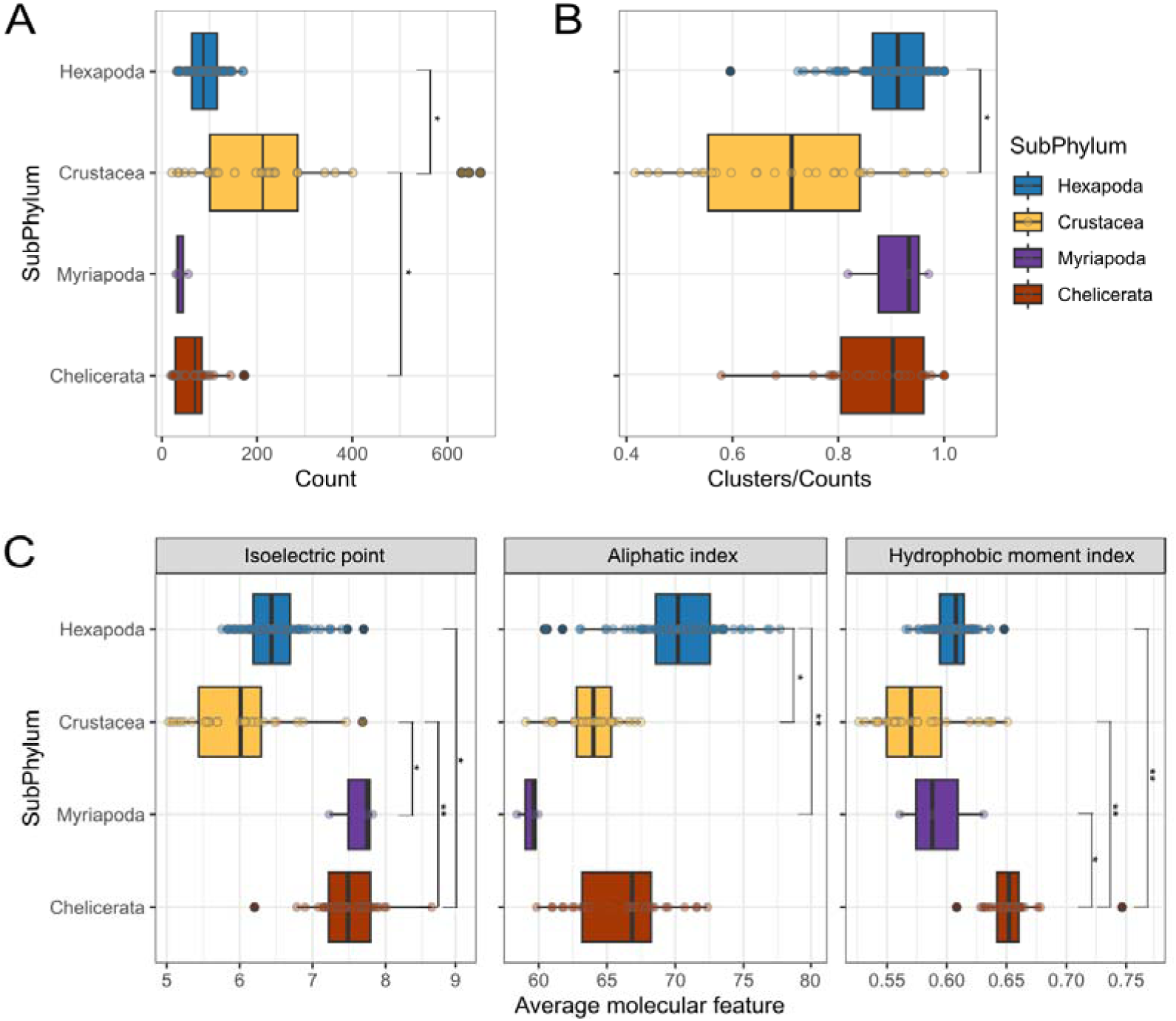
Canonical cuticle protein content across Arthropoda. **(A)** Raw counts of proteins annotated as harbouring the canonical chitin-binding domain across arthropod subphyla (asterisks indicate phylANOVA test p-values of *p<0.05). **(B)** Cuticle Protein (CP) repertoire redundancy. CPs are clustered within-species by 80% sequence similarity, and the ratio of clusters to counts approximates repertoire redundancy, with lower values indicating more sets of CPs with high sequence similarity (asterisks indicate phylANOVA test p-values of *p<0.05) **(C)** CP physico-chemical properties. Each data point represents the average value of all CPs found in a species for the isoelectric point, the aliphatic index, and the hydrophobic moment index (asterisks indicate phylANOVA test p-values of ** p<0.005, and *p<0.05).

Examining their physico-chemical properties shows significant differences between CPs of different subphyla. Crustacean and hexapod CPs are characterised by the lowest predicted isoelectric point (pI) values, contrasting Pancrustacea with the higher values in Myriapoda and Chelicerata (Figure 6C). High-pI CPs are more basic and bind to negatively charged chitin forming more rigid cuticles, while low-pI CPs are more acidic and help buffer pH, regulate hydration, or interact with positive ions resulting in softer or more flexible cuticles. Acidification below the pI might weaken inter-protein interactions, promoting elastic and plastic stretch, suggesting that lower pI values in Pancrustacea may facilitate cuticle expansion capacities (Reynolds 1975; Benoit and Denlinger 2010; Flynn and Kaufman 2015). The CP aliphatic index, where higher values indicate greater proportions of nonpolar amino acids, shows a different contrast among the subphyla, with the highest average values in hexapods, albeit with a wide range across species (Figure 6C). Proteins with an elevated aliphatic index are more hydrophobic and generally more heat-resistant, suggesting that the CP repertoires in many hexapods and some chelicerates are compositionally adapted to withstand higher temperatures. The hydrophobic moment index, where high values indicate clustering of hydrophobic residues to present a clearly defined hydrophobic face and low values mean the residues are evenly spread, is elevated in Chelicerata compared to the three mandibulate subphyla (Figure 6C). As CPs are anchored in a complex matrix containing chitin, other CPs, lipids, cross-linking agents, etc. their hydrophobic moment influences how they are positioned within the matrix and how they interact with other components (Moussian 2013). Low-moment CPs do not have distinct hydrophobic and hydrophilic faces and thus require higher levels of covalent sclerotisation for structural integrity while high-moment CPs may function in anchoring or organising cuticle layers. This suggests that in chelicerates the particular CP localisation and orientation in the extracellular space within the cuticular matrix may be functionally relevant (Politi et al. 2021). Taken together, these results demonstrate that CP repertoires and properties vary across major arthropod groups and likely reflect different evolutionary routes, enabling hypothesis generation about order-specific mechanisms in aquatic Crustacea and adaptation to life on land in Hexapoda and Chelicerata (Van Straalen 2021). Interestingly, lineage-specific patterns of co-expression in moulting among Pancrustacea appear driven in part by cuticle development (Kim et al. 2026).

## Discussion

The broad taxonomic sampling across 35 arthropod orders, including three newly sequenced and annotated myriapods, enables the examination of moulting gene repertoire evolution in deep time with fine granularity. Genomic surveying and gene copy-number profiling highlights the ancient origin of the moulting machinery toolkit, with overall lower levels of protein sequence divergence, and with contrasting stable and dynamic components from across the six functional modules governing moulting processes. The presence of the toolkit in the arthropod LCA is supported by previous studies focused on limited subsets of moulting-related genes describing the ancient origins of key ligand-receptor components (de Oliveira et al. 2019), members of the sesquiterpenoid and ecdysteroid pathways (Qu et al. 2015), the so-called “Halloween genes” (Schumann et al. 2018), or enzymes involved in the final steps of JH synthesis (Smykal and Dolezel 2023). Reconstructing ancestral gene contents across arthropods reveals numerous gain and loss events, with moulting-related gene families experiencing elevated expansions during the early diversification of arthropods. This contrasts the moulting gene repertoire with global gene content evolution where previous arthropod-wide (76 species) gene content analyses pointed to relatively evenly distributed rates of duplications and losses (Thomas et al. 2020). Gene duplications therefore appear to have played an important role in shaping the ancestral moulting machinery of arthropod subphyla, followed by more stable evolutionary trajectories within orders. This relative stability is observed even where substantial changes in moulting processes have occurred, such as in holometabolous insects, and is in contrast to variable and lineage-specific gene expression dynamics (Kim et al. 2026). The majority of protein domain architectures are also stable, with the exception of an over-representation of terminal loss or emergence events amongst the moulting gene set members, that suggests functional elaborations. The conserved and ancient core is nevertheless characterised by dynamic gene repertoire changes observed across all six moulting modules and nearly all 35 examined orders, with the ecdysteroid pathway and late genes showing elevated proportions of duplications. Indeed, global analyses previously identified ecdysteroid metabolism genes amongst the most dynamically changing arthropod gene families, alongside xenobiotic defence, digestion, chemosensation, chitin metabolism, and others (Thomas et al. 2020). Complementing phylogenetic profiling with ancestral state reconstructions of gene contents and protein domain architectures provides a robust approach to characterise the evolutionary dynamics of the arthropod moulting machinery.

One limitation to the characterisation of such patterns is where orders are represented by a single species (43%) or only two species (9%). The lack of “replicates” means that estimates are lacking support from detections across several genomes, and even correctly inferred events may not be representative of the order as a whole. Together, these biases may inflate inferred gains or losses in orders with few sampled species. Nevertheless, in some cases higher observed copy-numbers in single-species orders could reflect true gene gains, possibly arising from WGDs, e.g. in Xiphosura horseshoe crabs (Nong et al. 2021) or in Balanomorpha and Pollicipedomorpha barnacles (J.-H. Kim et al. 2019; Bernot et al. 2022; Yuan et al. 2024), or from high duplication levels reported for the Siphonaptera cat flea, *C. felis* (Driscoll et al. 2020; Feyereisen 2022). Assertions of gene losses across the arthropod phylogeny are particularly challenging where species representation is limited. For example, the representative isopod *A. nasatum* appears to lack several key moulting machinery components, but its genome annotation may be incomplete, so comparisons with other isopods are needed. Where multiple additional unannotated assemblies for a clade are available, targeted genome screening can provide the support needed to corroborate lineage-wide losses, such as the remarkable absence of *nvd* across Coleoptera. Fortunately, ongoing assembly and annotation of genomes from across the tree of life (Mazzoni et al. 2023; Blaxter et al. 2025) will enable future rigorous evaluations of such putative losses of key members of the moulting gene repertoire. This will need to be accompanied by experimental studies in diverse arthropods, to verify functional hypotheses based on phylogenomic comparisons, and to identify new components of the moulting machinery that may be specific to less well studied clades.

The moulting gene repertoire includes both predominantly single-copy and near-universally multi-copy families, reflecting genome-wide patterns in which constrained single-copy genes contrast with relaxed multi-copy families (Waterhouse et al. 2011). Additionally, there are numerous lineage-specific elaborations in the form of gene duplications maintained across descendant taxa (with or without subsequent protein domain changes), and lineage-wide losses of otherwise widely maintained genes. These lineage-specific elaborations, and the capacity for lineage-specific changes in multi-copy families, are likely to be directly or indirectly linked to differences in the variety of life histories across arthropod diversity, with key factors being dietary physiology, developmental modes, and environmental constraints.

With respect to diet, highly variable steroid composition from different food sources might promote diversification of organismal capabilities for hormone biosynthesis with implications for how moulting processes are regulated. This is supported by experimental evidence of a heterogeneous cocktail of ecdysteroid intermediates and analogues in different taxa that are able to elicit moult-inducing biological responses (Campli et al. 2024). For instance, instead of the insect 20E, ponasterone A (25-deoxy-20E) has been suggested as a major bioactive ecdysteroid in some lineages of Crustacea and Chelicerata. Regarding moulting and post-embryonic development, these are inextricably linked because moulting is both the mechanism that makes growth possible and the process that enables transitions. Toolkit and regulatory network differences must underlie the coordinated execution of developmental transitions that fall along a spectrum from gradual to metamorphic. Gradual developers require moults that preserve functional continuity and implement incremental morphological changes. Metamorphic developers undergo large-scale remodelling that enables transitions between ecologically and morphologically disparate life stages. In contrast to different dietary physiologies and developmental modes, environmental constraints may manifest more prominently in variations of the structural components of the moulting machinery rather than through regulatory changes. Here the same underlying exoskeleton renewal process is expected to reflect adaptations to marine and freshwater habitats as well as terrestrialisation events, semi-aquatic life histories, and the innovation of insect flight (Asano et al. 2019; Van Straalen 2021; Vargas et al. 2021; McNamara and Freire 2022).

Our phylogenetically broad and deep analyses of the major components involved in moulting reveal under-appreciated evolutionary dynamics, extend current understanding, and prompt future explorations across taxa and the moulting toolkit functional modules. The arthropod moulting machinery shows developmental flexibility by accommodating lineage and functional variability, within a framework of conserved molecular modules and phenotypic outcomes (True and Haag 2001; McColgan and DiFrisco 2024).

## Materials and Methods

### Myriapod sample collection, DNA and RNA sequencing

Animals were collected between February and April 2022-2023 in Givat HaArbaa, Jerusalem, Israel (GPS coordinates: 31°44′00″N 35°12′48″E). *Scolopendra cingulata* and *Scutigera coleoptrata* were kept in individual transparent plastic boxes with a substrate and large pieces of bark and fed weekly with *Blattella germanica* nymphs. *Archispirostreptus syriacus* individuals were kept together in a large transparent plastic container filled with substrate and egg cartons and were fed weekly with small pieces of vegetables. The substrate was obtained from the collection site. Animals were kept in a climate-controlled room with the following set parameters: temperature (25 ± 1°C), humidity (∼ 40%), and light:darkness regime (14h:10h). Additionally, the substrate was moistened weekly with double-distilled water. No special approvals, such as ethical or collection permits, were required for handling these species. The genomic vouchers are deposited in The National Nature Collections, The Hebrew University of Jerusalem under the following museum voucher numbers: HUJ-INV-CHIL764 (*S. cingulata*), HUJ-INV-CHIL763 (*S. coleoptrata*), HUJ-INV-DIP113 (female *A. syriacus*), and HUJ-INV-DIP114 (male *A. syriacus*).

Animals were anesthetised using carbon dioxide. Substrate particles were gently removed by washing with molecular biology-grade water. Animals were dissected, flash-frozen in liquid nitrogen, and ground into powder using a mortar and pestle. The DNA sample of *S. cingulata* was obtained from the head and several anterior body segments of the female individual. The gut was dissected out before flash-freezing to minimise bacterial contamination. The whole-body sample, excluding the gut, was used to extract the DNA sample of female *S. coleoptrata* specimen. The genome assembly of *A. syriacus* is based on two independent DNA extractions and sequencing procedures, one was obtained from the head and several anterior trunk segments of the female individual, and another sample was derived from the trunk segments of the male. The gut and legs were excluded from the extraction reaction in both cases. The optimal tissue amount for each species was determined experimentally and proved to be a crucial factor in the successful extraction of long DNA fragments. DNA extraction and RNAse treatment were performed using Monarch Genomic DNA Purification Kit (New England Biolabs). The samples of *S. cingulata* and *S. coleoptrata* were extracted following the manufacturer’s protocol. The body *A. syriacus* has a well-developed parietal fat body which complicates the extraction of clean DNA samples, requiring a modified protocol. After the lysis reaction and centrifugation, the supernatant was slowly mixed with phenol:chloroform;isoamyl alcohol (25:24:1) saturated with 10 mM Tris, pH 8.0, 1 mM EDTA (Sigma) for 3 hours, and the DNA from the aqueous phase was then precipitated with ethanol to obtain a crude DNA sample. This sample then underwent RNase treatment and column purification, following the manufacturer’s protocol. The purity and the concentration of DNA samples were assessed using NanoDrop 1000 (Thermo Fisher Scientific), the concentration was additionally confirmed on Qubit Flex Fluorometer using Qubit 1X dsDNA HS Assay Kit (Thermo Fisher Scientific). Samples were stored at -80°C and transported in dry ice to the Lausanne Genomic Technologies Facility for further processing. 5200 Fragment Analyzer system (Agilent) was used to evaluate the fragmentation level of the samples. For the genome annotation, RNA was extracted from several body parts, including whole-body samples without gut, head, ovaries, and testicles. The Quick-RNA MiniPrep Kit (Zymo Research, R1055) was used for RNA extraction, following the manufacturer’s instructions. The purity and the concentration of the obtained RNA samples were estimated with NanoDrop 1000 (Thermo Fisher Scientific). RNA integrity was analysed using a Bioanalyzer system (Eukaryote Total RNA Nano kit, Agilent). RNA libraries were prepared using Illumina Stranded mRNA Prep kit and sequenced on a PE150 AVITI (Element Biosciences) at the Lausanne Genomic Technologies Facility.

### Myriapod genome assembly and annotation

High-quality genomic DNA was sequenced using PacBio HiFi sequencing on a PACMAN instrument, generating highly accurate long reads. The sequencing coverage was approximately 46× for *S. cingulata*, 40× for *S. coleoptrata*, and 30× for *A. syriacus*. The quality of the reads was assessed using LongQC (v1.2.0c) (Fukasawa et al. 2020), while genome characteristics, including ploidy and heterozygosity, were estimated using Smudgeplot (v0.2.5) and GenomeScope2 (v2.0) (Ranallo-Benavidez et al. 2020). Genome assembly was performed using Hifiasm (v0.19.2) (Cheng et al. 2021) and/or Purge_dups (v1.2.6) (Guan et al. 2020), applying different purging strategies depending on the species. For *S. cingulata*, assembly was performed exclusively with Hifiasm using its strongest purging level (-l3). For *A. syriacus* and *S. coleoptrata*, Hifiasm was first applied with the strongest purging setting (-l3) and further refined using Purge_dups. Assembled genomes were screened for contaminants using fcs_gx (v0.5.0) (Astashyn et al. 2024) before final quality assessment. Genome completeness and accuracy were evaluated using the Benchmarking Universal Single-Copy Orthologues (BUSCO) (v5.4.3) assessment tool with the arthropoda_odb10 database (Manni et al. 2021). Additional assembly statistics were obtained using gfastats (v1.3.6) (Formenti et al. 2022), and k-mer-based quality assessment was performed using Merqury (v1.3) (Rhie et al. 2020). The genome assemblies exhibited high completeness and contiguity, with BUSCO scores and N50 values reflecting well-assembled genomes across all three species. The *S. cingulata* assembly is composed of 600 contigs, an N50 of 63,664,107 bp, and a BUSCO completeness of C:97.6% [S:95.2%, D:2.4%], F:0.2%, M:2.2%, n:1013. *S. coleoptrata* was assembled into 344 contigs, with an N50 of 20,554,343 bp and BUSCO scores of C:97.1% [S:96.1%, D:1.0%], F:1.2%, M:1.7%, n:1013. The *A. syriacus* assembly contained 3,887 contigs, with an N50 of 1,010,713 bp and BUSCO completeness of C:96.5% [S:94.2%, D:2.3%], F:1.8%, M:1.7%, n:1013.

Genome annotation was performed by first identifying repetitive elements using RepeatModeler (v2.0.4) (Flynn et al. 2020). Protein-like transposable elements (TEs) were filtered out using BLAST (v2.7.1) (Camacho et al. 2009), transposonPSI (v1.0.0) and ProtExcluder (v1.2) (Campbell et al. 2014). The remaining repetitive elements were then masked in the genome using RepeatMasker (v4.1.5) (Smit et al. 2013). To facilitate gene prediction, RNA-seq data were generated for each species and supplemented with publicly available datasets from the Sequence Read Archive (SRA). For *S. cingulata* and *A. syriacus*, RNA was extracted from an adult female’s head and a few anterior body rings, excluding the gut. For *S. coleoptrata*, RNA was extracted from the head and the whole body of an adult female, also excluding the gut. The RNA samples were sequenced using the Illumina NovaSeq platform, generating strand-specific mRNA libraries with paired-end reads. To enhance the comprehensiveness of the transcriptome data, publicly available RNA-seq datasets from SRA were also used. For *S. cingulata*, SRR1653235 was included, which contains RNA extracted from the anterior half of the trunk, excluding the head and poison claws. For *S. coleoptrata* these included, SRR1653237, derived from a whole organism excluding the head and poison claw, SRR8998264, extracted from the forcipule, and SRR1158078, for which the tissue of origin is unknown. RNA-seq data quality was assessed with FastQC (v0.12.1) (Andrews 2010), and sequencing adaptors and low-quality reads were removed using Atropos (v1.1.28) (Didion et al. 2017). The filtered RNA-seq reads were aligned to the genome using HISAT2 (v2.2.1) (D. Kim et al. 2019), and mapping statistics were generated using SAMtools (v1.17) (Li et al. 2009) and QualiMap (v2.2.2d) (Okonechnikov et al. 2016). Protein-coding gene annotation was performed using BRAKER3 (v3.0.8) (Gabriel et al. 2024), integrating the previously aligned RNA-seq data along with curated protein evidence from OrthoDB (Tegenfeldt et al. 2025). The completeness of the predicted annotation was evaluated using BUSCO (v5.4.3), provided in Supplementary Table 1.

### Publicly sourced genome assemblies and annotations

To obtain a broad representation of arthropod diversity for orthology delineation the completeness of the arthropod genome assemblies available from the United States National Center for Biotechnology Information (NCBI) was evaluated using the Arthropod Assembly Assessment Catalogue (A3Cat, release 2024-07-01) (Feron and Waterhouse 2022b). Assemblies were required to have an annotated proteome with the number protein-coding genes ranging from 8’000 to 50’000. These were then filtered based on the A3Cat BUSCO scores: the annotated assembly having the highest Arthropoda Complete BUSCO score was retained where more than one assembly was available for individual species; all selected assemblies were required to have scores of less than 40% Arthropoda Duplicated BUSCO and at least 75% Arthropoda Complete BUSCO. These thresholds were applied specifically to avoid the exclusion of some non-insect arthropods from key underrepresented lineages required to ensure a broad representation of arthropod diversity. Subsequently, to produce a more balanced phylogenetic diversity, insect taxonomic orders with more than 12 species were downsampled, by retaining within each order one representative for each taxonomic family, having the highest Arthropoda Complete BUSCO score among the species of that family. Additionally, representative species for the families Drosophilidae (Diptera), Tenebrionidae (Coleoptera), and Bombycidae (Lepidoptera) were manually selected: *D. melanogaster* (Release_6_plus_ISO1_MT), *T. castaneum* (GCF_031307605.1), and *B. mori* (GCF_030269925.1), respectively, given their well-established history as model systems and the accumulated knowledge of genes involved in moulting process. Proteomes and annotation feature tables from the selected assemblies were downloaded from NCBI repositories GenBank and RefSeq (2024-07-12) and assessed for completeness using BUSCO v5.4.3 and the arthropoda_odb10 dataset (option: -mode protein) (Manni et al. 2021). All annotated proteomes with a score of at least 75% Arthropoda Complete BUSCO were retained and filtered for the protein-coding longest isoform based on their corresponding annotation feature table. This resulted in the selection of the annotated proteomes of 145 arthropod species representing 35 orders from the four major subphyla, with Arthropoda BUSCO Complete scores of median 96.2% (mean 91.4%, standard deviation 6.0%) and Arthropoda BUSCO Duplicated scores of median 1.9% (mean 3.8%, standard deviation 5.8%), and protein-coding gene counts of median 15’649 (mean 17’603, standard deviation 6’723%). Assembly accessions, protein-coding gene counts, and BUSCO assessments are provided in Supplementary Table 1.

### Species phylogeny reconstruction

The species phylogeny was computed from single-copy orthologues found in at least 95% of the total set of 145 species, detected by the BUSCO assessments performed at A3Cat on the genome assemblies using the arthropoda_odb10 dataset. Protein sequences were aligned using MUSCLE v3.8.1551 (Edgar 2004) with default parameters and trimmed using the automated method for threshold optimization in TrimAl v1.4.1 (option -strictplus) (Capella-Gutiérrez et al. 2009). A concatenated super-alignment was built from the individual alignments (76’471 columns, 71’014 distinct patterns, 57’142 parsimony-informative, 9’412 singleton sites, 9’917 constant sites; other metrics computed by Alistat v1.15 (Wong et al. 2020) are available as Supplementary Table 2) for the phylogeny reconstruction performed using ModelFinder as implemented in IQ-TREE v2.2.0-beta with 1’000 ultrafast bootstrap samples (option -msub nuclear -B 1000 -bnni) (Nguyen et al. 2015; Kalyaanamoorthy et al. 2017; Hoang et al. 2018; Minh et al. 2020). The resulting phylogeny was inspected and manually rooted in NJ-plot using chelicerate species as the outgroup (Perrière and Gouy 1996). The paraphyletic crustaceans, comprising Hexapoda and Crustacea within the monophyletic Pancrustacea were recovered, whereas three orders each represented by only a single species were observed to be in disagreement with previous phylogenomics findings: placement of the horseshoe crab *L. polyphemus* (Xiphosura, Chelicerata) within Arachnida, positioning of the copepod *T. californicus* (Harpacticoida, Crustacea) as sister to species of the Branchiopoda class (namely Diplostraca and Anostraca), and the dragonfly *L. fulva* (Odonata, Hexapoda) not clustering with Ephemeroptera species. Therefore, a second round of phylogeny reconstruction in IQ-TREE was performed applying constraints to achieve the correct topology as follows: early branching of *L. polyphemus* before the Arachnida LCA, diverging of *T. californicus* from the LCA of Malacostraca and Thecostraca, and Ephemeroptera species sharing the most recent common ancestor with *L. fulva* (Misof et al. 2014; Schwentner et al. 2018; Howard et al. 2020). The molecular species tree was time-calibrated using ten calibration nodes (Supplementary Table 3), supplied to the functions makeChronosCalib() and chronos() from the ape v5.8 R package (Paradis and Schliep 2019). Divergence time estimates were sourced from the TimeTree v5 database (Kumar et al. 2022). The resulting ultrametric species phylogeny representing 600 million years of arthropod evolution with 144 internal nodes was used for ancestral state reconstructions and gene-tree-species-tree reconciliation analyses.

### Orthology delineation across 145 arthropod species

Orthology was delineated at the level of the Last Common Ancestor (LCA) of all the 145 selected arthropod species, where an orthologous group (OG) represents the set of genes descended from a single gene in the arthropod LCA. OGs were reconstructed from 142 publicly available annotated proteomes and three myriapod species newly assembled and annotated in this study. The OrthoDB standalone pipeline OrthoLoger v3.0.2 was run with default parameters (Kuznetsov et al. 2023), after filtering the proteomes for the single longest protein-coding isoform of each gene. The OrthoLoger algorithm uses all-against-all pairwise protein sequence alignments to identify best reciprocal hits (BRHs) between all genes from each species pair in the dataset. Subsequently, a graph-based clustering procedure starting with BRH triangulation progressively builds OGs aiming to identify and include all genes descended from a single gene in the LCA. The resulting orthology dataset comprised a total of 74’214 OGs and 2’047’671 genes, representing 80% of the input set of 2’552’371 genes from the annotated proteomes of 145 species. OG phyletic ages were computed as the LCA of all the species included in the OG, simplified to stratifications of Arthropoda, Mandibulata (Pancrustacea + Myriapoda), the four subphyla, and younger than subphyla. OG evolutionary rates were computed using OrthoLoger ancillary tools: pairwise protein alignments are used to calculate average of inter-species identities for each OG, normalised over the average identity of all inter-species BRH, as defined by OrthoDB (Waterhouse et al. 2013).

### Identification of moulting-related genes from the orthology dataset

The lists of genes with evidence of involvement in various moulting processes was curated based on an extensive literature review of the accumulated knowledge of the genetic toolkit governing arthropod moulting (Campli et al., 2024) and manual searches of the FlyBase database (Öztürk-Çolak et al. 2024). The curated list of gene names and NCBI protein identifiers comprises primarily genes from the well-studied *D. melanogaster* fruit fly (n=74), complemented by a further 48 genes from other relatively well-studied insects *B. mori* and *T. castaneum*, decapods *L. vannamei* and *P. trituberculatus*, and the common house spider *P. tepidariorum* (Supplementary Table 4). Considering evidence from the literature, these genes collectively represent 76 moulting-related genes, or targeted gene families of interest for this genomic survey. The protein identifiers were collated to match the versions of the annotated proteomes used for orthology delineation so the gene lists could be used as “seeds” to search the orthology dataset and identify relevant OGs representing the gene families of interest. The dataset was manually curated to reflect current knowledge: in order to reflect the shared history of *stw* with *mco1*, and *chs2* with *kkv* (Arakane et al. 2005; Moussian 2010; Asano et al. 2019), their individual OGs were merged together, respectively. Profiling of orthologue copy-numbers per species for each gene family of interest across the main functional categories/modules and taxonomic groups was performed in R v4.3.2 and plotted using the ggplot2 package (Wickham 2016).

### Gene-tree-species-tree reconciliation

The process of reconciling a gene tree with a species tree aims to map gene duplication events, gene loss events, and speciation events, given a model of gene family evolution, *i.e.* to trace the evolutionary history of a gene family. Confidence in the inferred evolutionary scenarios requires robust input gene trees, which can be challenging to achieve with large sets of diverse species. Given the focus on characterising deep-time events (*i.e.* mainly order-level and older), and that analysing fewer species produces more confident gene trees, a subsampling of the full orthology dataset was performed before gene-tree-species-tree reconciliation. For the insect orders of Diptera, Lepidoptera, Hymenoptera, Coleoptera, and Hemiptera, species sets were downsampled by choosing for each order the subset of species having the five highest Arthropoda Complete BUSCO scores computed on the proteome. Additionally, for the other arthropod orders, where the same genus was present more than once, a single species for each genus was selected using the same criteria, overall resulting in a subsampled dataset of 97 species. The species tree used for the reconciliations was obtained by pruning the 145-species phylogeny to retain these 97 species. The gene families analysed consisted of the moulting-related genes identified from the orthology dataset. Protein sequences from each gene family were aligned using the iterative refinement L-INS-i algorithm from MAFFT v7.520 (Katoh and Standley 2013). Alignments were trimmed using TrimAl v1.4.rev15, firstly with the heuristic automatic method (option: -automated1) and further by removing fragmented protein sequences that could represent incomplete gene annotations (option -resoverlap 0.7 -seqoverlap 70) (Capella-Gutiérrez et al. 2009). Gene trees were reconstructed using maximum likelihood estimates as implemented in RAxML v8.2.12, choosing the GAMMA model and JTT substitution matrix and performing 1000 rapid bootstrap samples (option -f a -N 1000 -m PROTGAMMAJTT) (Stamatakis 2014). Gene trees were rooted in the Python ete3 module, using the chelicerate sequences as outgroup when monophyletic, otherwise when polyphyletic or not present in the tree, midpoint rooting was applied (Huerta-Cepas et al. 2016). For analysing only the most robust input gene trees, only those with more than 50% of the nodes supported by bootstrap values higher than 50% were then used for reconciliations with the species tree. Gene-tree-species-tree reconciliation was performed using the parsimony approach of NOTUNG v.2.9.1.5, under a Duplication and Loss model with default costs and allowing the rearrangement of branches at nodes with less than 90% bootstrap support (option --rearrange --threshold 90 --infertransfers false) (Stolzer et al. 2012). The summary of deep time events from selected gene families is presented in Supplementary Table 5.

### Gene count ancestral state reconstruction

Ancestral state reconstruction aims to estimate the likely state (in this case gene copy numbers) at the internal nodes of the species tree given the observed state at the terminal branches and a model of gene family evolution. To estimate ancestral gene copy numbers, and thus map gene family expansion and contraction events on the species tree, a maximum likelihood approach to the birth-death model of gene family size evolutionary rate was employed, as implemented in Computational Analysis of gene Family Evolution (CAFE) v5 (Mendes et al. 2021). Firstly, the phyletic age of each OG was calculated as the LCA of all the species represented in the OG, considering the full species phylogeny. Then, only OGs with a phyletic age of Arthropoda LCA and with species included in subsampled dataset (97 species set) were used for the input gene count table provided to CAFE. Two separate sets of CAFE runs were performed, (1) for the foreground (all the proteins identified in moulting-related OGs as old as the Arthropoda LCA), and (2) for the background (all the proteins belonging to OGs as old as the Arthropoda LCA, excluding the foreground). The background was further filtered to remove gene families with more than 100 gene copies in one or more species, following CAFE guidelines (using the provided Python script, clade_and_size_filter.py). To ensure model convergence, each set was run independently four times under different parameter settings, always estimating lambda with one k rate category. For the background set, Poisson and normal root family size distributions, and different lambda rates up to four (one for each subphyla) were tested. Using the poisson distribution and two lambda rates (one for Chelicerata and one for Mandibulata), resulted in estimation of lambda parameters respectively as 0.0017 and 0.0018, supported by a maximum likelihood score of -lnL 1.30946e+06. For the foreground set, gene count ancestral state reconstruction was performed using the output from the background set run, resulting in lambda parameter estimates of 0.0009, 0.0018 with a maximum likelihood score of -lnL 7921.63.

### Protein domain evolutionary history analyses

Domain annotation of the longest protein isoforms was performed using PfamScan (Mistry et al. 2007). Consecutive repeated annotations were collapsed (ie. A-B-B-B-C -> A-B-C) as indicated for ancestral domain rearrangement reconstruction in DomRates (Dohmen et al. 2020). Two separate runs using the set of Chelicerata species as the outgroup were performed for the foreground (all the proteins involved in the moulting process in OGs as old as the Arthropoda LCA) and the background (all the proteins belonging to OGs as old as the Arthropoda LCA, excluding the foreground and the proteins without an ortholog), as for the ancestral state count reconstruction. Fisher tests were performed in R stats package v4.3.2 (R Core Team 2023).

### Support for clade-wide gene losses using *de novo* gene finding with MetaEuk

To further assess potential gene losses observed from the proteome-based orthology analysis, the A3cat resource was used to identify genome assemblies without protein-coding annotations available from the NCBI. When multiple assemblies were available for a single species, the assembly with the highest Arthropoda Complete BUSCO score was selected. Only assemblies with at least 75% BUSCO score were retained to perform *de novo* gene finding with MetaEuk (Levy Karin et al. 2020). Genome-wide searches were carried out using the longest protein isoforms of the genes of interest (seed protein identifiers in Supplementary Table 6) as input to MetaEuk (option: --exhaustive-search).

### Annotation and analysis of cuticle protein gene families

Results from PfamScan were filtered for the domain “PF00379.27” (“insect cuticle protein”, “chitin_bind_4”) to retrieve proteins annotated as canonical cuticle proteins (CPs) harbouring the Rebers and Riddiford (RR) chitin binding consensus motif, as indicated in Willis, 2010 (Willis 2010). CPs of each subtype, in each species, were clustered by 80% sequence similarity using CD-HIT (v4.8.1) (Li and Godzik 2006). Molecular properties such as aliphatic index, hydrophobic moment index, and isoelectric point of each CP were computed using the Peptides package (Osorio et al. 2015).

## Data Availability

The genomic data underlying this article are available from the European Nucleotide Archive (ENA) at https://www.ebi.ac.uk/ena/browser/home, and can be accessed with the accession numbers provided in Supplementary Table 1. The orthology dataset generated as part of this study is available at figshare DOI: 10.6084/m9.figshare.31082677. The genome assemblies and annotations for the three myriapod species produced as part of this study are available at figshare DOI: 10.6084/m9.figshare.31077730 and DOI: 10.6084/m9.figshare.31077829, respectively, and are being processed for submission to the ENA.

## Acknowledgements

The authors thank all members of the Sinergia project “An interdisciplinary study of arthropod moulting: linking genotype, phenotype and life history evolution” for useful discussions that supported the development of this manuscript. We are also grateful to Rosa Fernández for making available RNAseq data from *Scutigera coleoptrata* to assist with genome annotation, and to Romain Ferron for assistance with building the arthropod species phylogeny.

