## Supplementary Figures for "Evolutionary dynamics of the arthropod moulting machinery"

### CONTENTS

Supplementary Figure 1: Species tree and orthology 2
Legend: Supplementary Figures 2-7. Gene family copy-number profiling of the moulting machinery. 4
Supplementary Figure 2: Ecdysteroid pathway module 5
Supplementary Figure 3: Sesquiterpenoid pathway module 6
Supplementary Figure 4: Early gene module 7
Supplementary Figure 5: Fate gene module 8
Supplementary Figure 6: Neurostimuli reception module 9
Supplementary Figure 7: Late gene module 10
Supplementary Figure 8: Ancestral reconstruction of protein domain architecture rearrangements 11
Legend: Supplementary Figures 9-12. Species-tree gene-tree reconciliations of gene families with deep time duplications 11
Supplementary Figure 9: Species-tree gene-tree reconciliation for serp/verm family 12
Supplementary Figure 10: Species-tree gene-tree reconciliation for Cda4 family 13
Supplementary Figure 11: Species-tree gene-tree reconciliation for kkv/Chs2 family 14
Supplementary Figure 12: Species-tree gene-tree reconciliation for knk family 15
Supplementary Figure 13: Ancestral loss of the deacetylase domain in the CDA1/serp-CDA2/verm family 16

**Supplementary Figure 1: Species tree and orthology**

**(A)** The species phylogeny shows the relationships and estimated divergence times of the complete dataset of 145 arthropod species. The tree is annotated with the 35 represented orders. The phylogeny was estimated using a super-alignment of protein sequences from single-copy orthologues and time-calibrated with ten calibration nodes. The four major arthropod lineages are highlighted: Chelicerata (pink), Myriapoda (purple), Crustacea (yellow), and Hexapoda (green), where hexapods and the paraphyletic crustaceans together form the monophyletic Pancrustacea. **(B)** Orthology delineation at the level of the last common ancestor (LCA) of the complete dataset of 145 arthropod species resulted in the orthologous group (OG) dataset comprising a total of 74'214 OGs and 2'047'671 genes. The bars show the counts of genes in OGs classified as single-copy (blue) or multi-copy (green, yellow) orthologues, or for which no orthologues could be identified (grey).

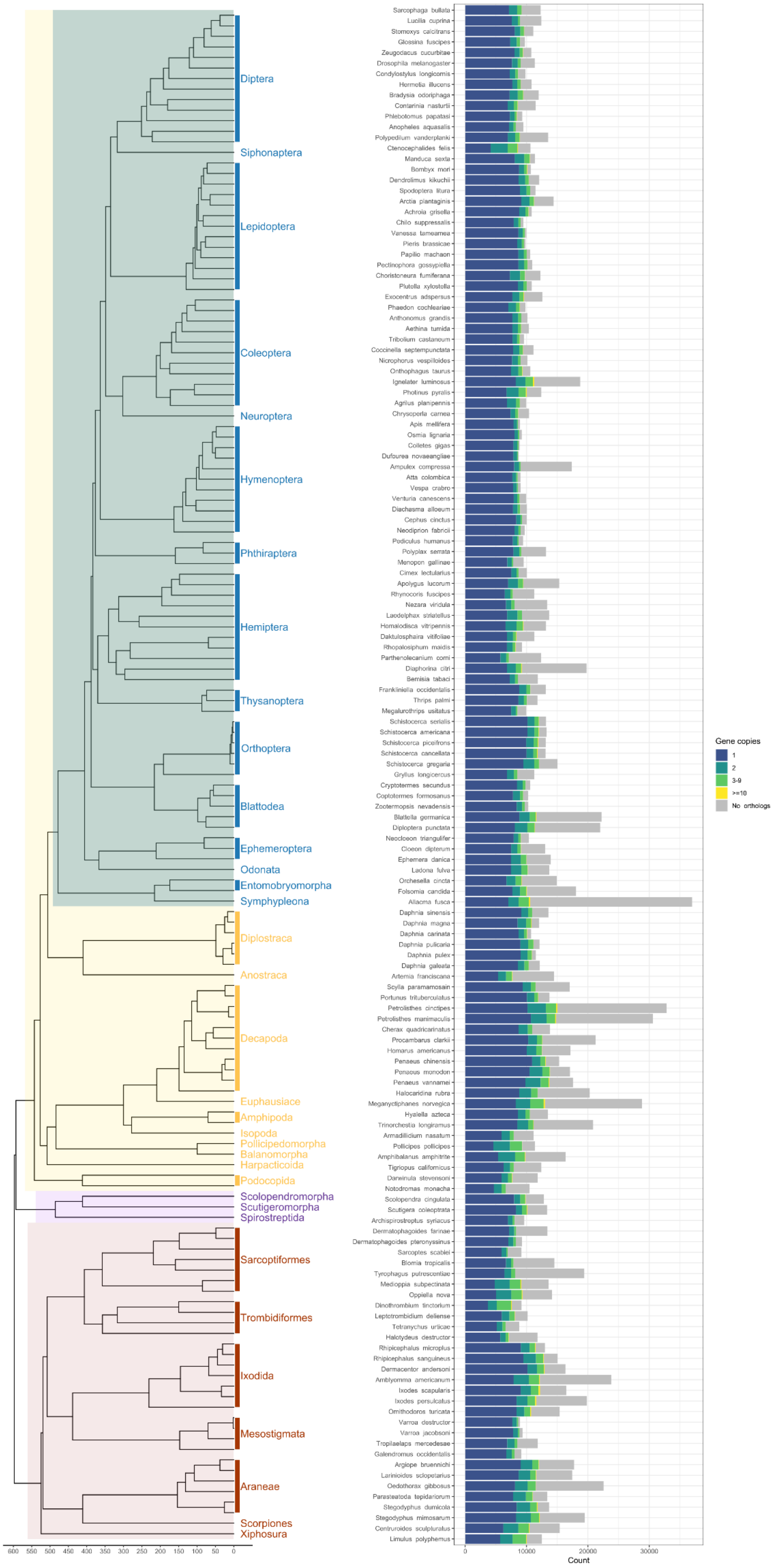
Diptera
Siphonaptera
Lepidoptera
Coleoptera
Neuroptera
Hymenoptera
Phthiraptera
Hemiptera
Thysanoptera
Orthoptera
Blattodea
Ephemeroptera
Odonata
Entomobryomorpha
Symphypleona
Diplostraca
Anostraca
Decapoda
Euphausiacea
Amphipoda
Isopoda
Pollicipedomorpha
Balanomorpha
Harpacticoida
Podocopida
Scolopendromorpha
Scutigeromorpha
Spirostreptida
Sarcoptiformes
Trombidiformes
Ixodida
Mesostigmata
Araneae
Scorpiones
Xiphosura
600 550 500 450 400 350 300 250 200 150 100 50 0
Sarcophaga bullata
Lucilia cuprina
Stomoxys calcitrans
Glossina fuscipes
Zeugodacus cucurbitae
Drosophila melanogaster
Condylostylus longicornis
Hermetia illucens
Bradysia odoriphaga
Contarinia nasturtii
Phlebotomus papatasi
Anopheles aquasalis
Polypedilum vanderplanki
Ctenocephalides felis
Manduca sexta
Bombyx mori
Dendrolimus kikuchii
Spodoptera litura
Arctia plantaginis
Achroia grisella
Chilo suppressalis
Vanessa tameamea
Pieris brassicae
Papilio machaon
Pectinophora gossypiella
Choristoneura fumiferana
Plutella xylostella
Exocentrus adspersus
Phaedon cochleariae
Anthonomus grandis
Aethina tumida
Tribolium castaneum
Coccinella septempunctata
Nicrophorus vespilloides
Onthophagus taurus
Ignelater luminosus
Photinus pyralis
Agrilus planipennis
Chrysoperla carnea
Apis mellifera
Osmia lignaria
Colletes gigas
Dufourea novaeangliae
Ampulex compressa
Atta colombica
Vespa crabro
Venturia canescens
Diachasma alloeum
Cephus cinctus
Neodiprion fabricii
Pediculus humanus
Polyplax serrata
Menopon gallinae
Cimex lectularius
Apolygus lucorum
Rhynocoris fuscipes
Nezara viridula
Laodelphax striatellus
Homalodisca vitripennis
Daktulosphaira vitifoliae
Rhopalosiphum maidis
Parthenolecanium corni
Diaphorina citri
Bemisia tabaci
Frankliniella occidentalis
Thrips palmi
Megalurothrips usitatus
Schistocerca serialis
Schistocerca americana
Schistocerca piceifrons
Schistocerca cancellata
Schistocerca gregaria
Gryllus longicercus
Cryptotermes secundus
Coptotermes formosanus
Zootermopsis nevadensis
Blattella germanica
Diploptera punctata
Neocloeon triangulifer
Cloeon dipterum
Ephemera danica
Ladona fulva
Orchesella cincta
Folsomia candida
Allacma fusca
Daphnia sinensis
Daphnia magna
Daphnia carinata
Daphnia pulicaria
Daphnia pulex
Daphnia galeata
Artemia franciscana
Scylla paramamosain
Portunus trituberculatus
Petrolisthes cinctipes
Petrolisthes manimaculis
Cherax quadricarinatus
Procambarus clarkii
Homarus americanus
Penaeus chinensis
Penaeus monodon
Penaeus vannamei
Halocaridina rubra
Meganyctiphanes norvegica
Hyalella azteca
Trinorchestia longiramus
Armadillidium nasatum
Pollicipes pollicipes
Amphibalanus amphitrite
Tigriopus californicus
Darwinula stevensoni
Notodromas monacha
Scolopendra cingulata
Scutigera coleoptrata
Archispirostreptus syriacus
Dermatophagoides farinae
Dermatophagoides pteronyssinus
Sarcoptes scabiei
Blomia tropicalis
Tyrophagus putrescentiae
Medioppia subpectinata
Oppiella nova
Dinothrombium tinctorium
Leptotrombidium deliense
Tetranychus urticae
Halotydeus destructor
Rhipicephalus microplus
Rhipicephalus sanguineus
Dermacentor andersoni
Amblyomma americanum
Ixodes scapularis
Ixodes persulcatus
Ornithodoros turicata
Varroa destructor
Varroa jacobsoni
Tropilaelaps mercedesae
Galendromus occidentalis
Argiope bruennichi
Larinioides sclopetarius
Oedothorax gibbosus
Parasteatoda tepidariorum
Stegodyphus dumicola
Stegodyphus mimosarum
Centruroides sculpturatus
Limulus polyphemus
0
10000
20000
30000
Count
Gene copies
1
2
3-9
>=10
No orthologs

**Legend: Supplementary Figures 2-7. Gene family copy-number profiling of the moulting machinery.**

Phylogenetic profiling of gene copy numbers of individual families (upper panel) with evidence of involvement in various moulting processes quantified counts of identified orthologues across the 145-species dataset, partitioned by the 35 represented orders (rows) and six functional modules of the moulting machinery: the ecdysteroid pathway (Supplementary Figure 2) and sesquiterpenoid pathway (Supplementary Figure 3) for hormone synthesis and signalling, the early genes (Supplementary Figure 4) and fate genes (Supplementary Figure 5) for activation of the moult-inducing programme; the toolkit of receptors of neuropeptides (Supplementary Figure 6) switching on the moulting process, and downstream late gene effectors for exoskeleton renewal (Supplementary Figure 7). The bars show the average proportional counts across taxonomy orders as single-copy (blue) or multi-copy (green, yellow) orthologues, or for which no orthologues could be identified (deep purple). Cells show the counts of genes in orthologous groups (OGs) classified as single-copy (blue) or multi-copy (green, yellow) orthologues, or for which no orthologues could be identified (deep purple). The ecdysteroid pathway includes neverland (nvd), shroud (sro), spook and spookier (spo;spok), phantom and cytochrome P450 18a1 and 15 (phm;cyp18a1;cyp15), disembodied (dib), shadow (sad), ecdysone importer (EcI), ATP-binding cassette transporter expressed in trachea (Atet), shade (shd), membrane steroid binding protein (MSBP), sterol regulatory element binding protein (SREBP), ecdysoneless (ecd), Gcn5 acetyltransferase (gcn5), ecdysone receptor (EcR) and ultraspiracle (Usp); the sesquiterpenoid pathway includes Juvenile Hormone acid methyl-transferase (JHAMT), epoxide hydrolase (JHEH), esterase binding protein (JHEBP), Juvenile Hormone binding protein (JHBP), FK506-binding protein 39kD (FKBP39), Chd64, methoprene tolerant (met) and taiman (Tai); the early genes include Ecdysone-inducible 74 (E74), 75 and 78 (E75;E78), beta Fushi-tarazu transcription factor 1 (βftz-F1), hormone receptor 3 (HR3), 4 (HR4), 38 (HR38), 39 (HR39), 78 (HR78), 96 (HR96), Hepatocyte nuclear factor 4 (Hnf4), Blimp-1, cryptocephal (crc) and tailess (tll), ; the fate genes include chronologically inappropriate morphogenesis (chinmo), kruppel homologue-1 (Kr-h1), Broad and Ecdysone-inducible 93 (E93); the toolkit of receptors of neuropeptides torso, ecdysis triggering hormone receptor (ETHR), eclosion hormone receptor (EHR), crustacean cardioactive peptide receptor (CCAP-R) and rickets (rk); and the late gene effectors include chitin synthase 1 krotzkopf verkehrt and chitin synthase 2 (kkv;Chs2), chitinase 3 (Cht3) 5, (Cht5), 6 (Cht6), 7 (Cht7), 11 (Cht11), 8 and 2 and 4 and 9 (Cht8;Cht2;Cht4;Cht9), serpentine and vermiform (serp;verm), chitin deacetylase 3 (Cda3), 4 (Cda4), 5 (Cda5), 9 (Cda9), farnesoic acid binding protein (FABP), knickkopf (knk), retroactive (rtv), resilin, straw and multi-copper protein 1 (stw;mco1) and phenoloxidase 1 and 2 and 3 (PPO1;PPO2;PPO3) .

**Supplementary Figure 2: Ecdysteroid pathway module**

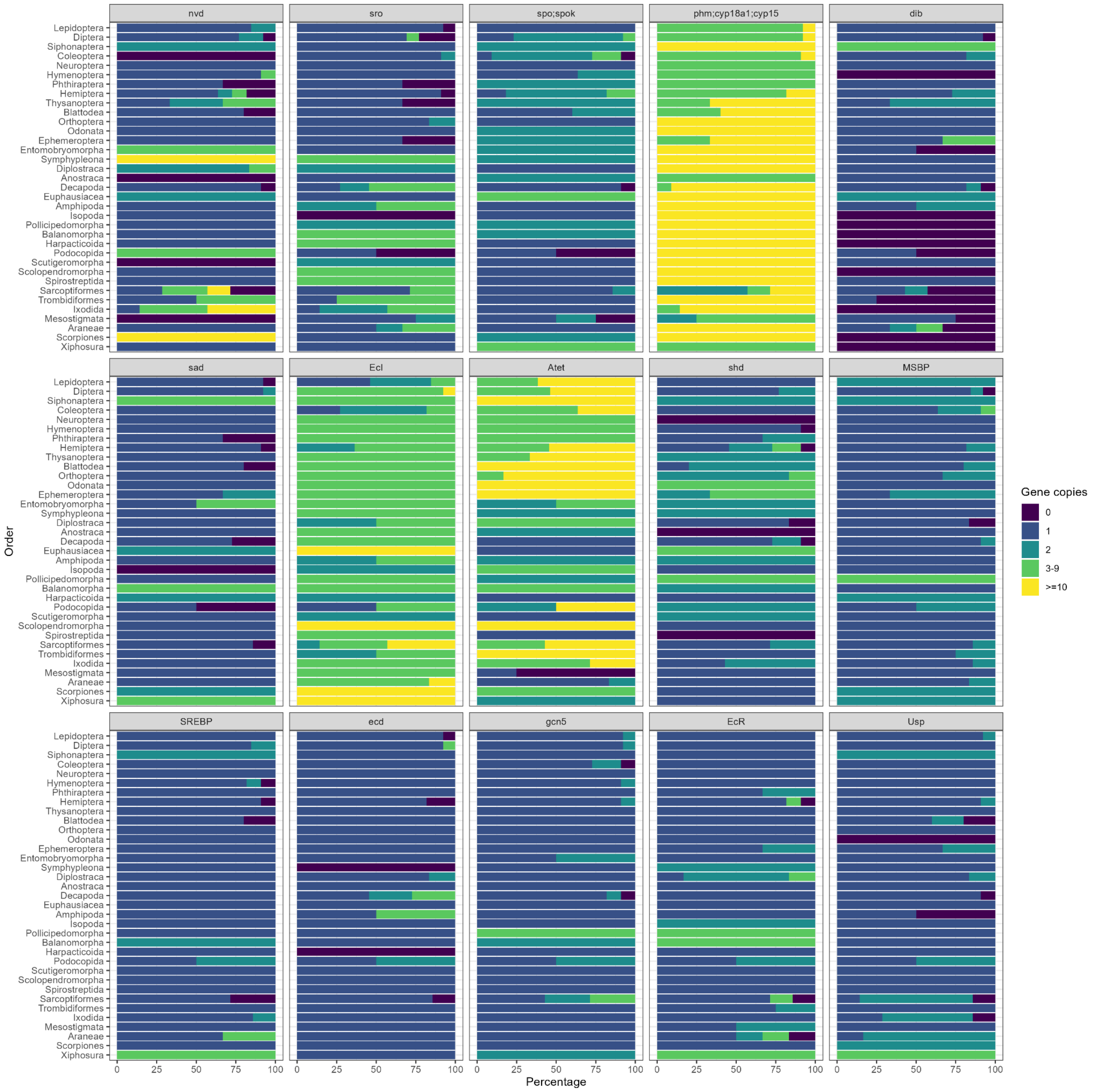


Notes: For the *phm-cyp18a1-cyp15* OG, the cyp15 members should cluster separately, so the expansions are inflated by these CYP2 clan P450s that are abundant especially in non-insect arthropods, where differential gene gains and losses have resulted in the orthology delineation co-clustering cyp15 genes with *phm* and *cyp18a1*. For the *disembodied* (*dib*) OG, evidence from the Arthropod P450 Enchiridion (http://arthropodp450.eu/), indicate it is present in Hymenoptera, Isopoda, Harpacticoida, Ixodida, Scorpiones, and Xiphosura, and that it is absent from barnacles.

### Supplementary Figure 3: Sesquiterpenoid pathway module

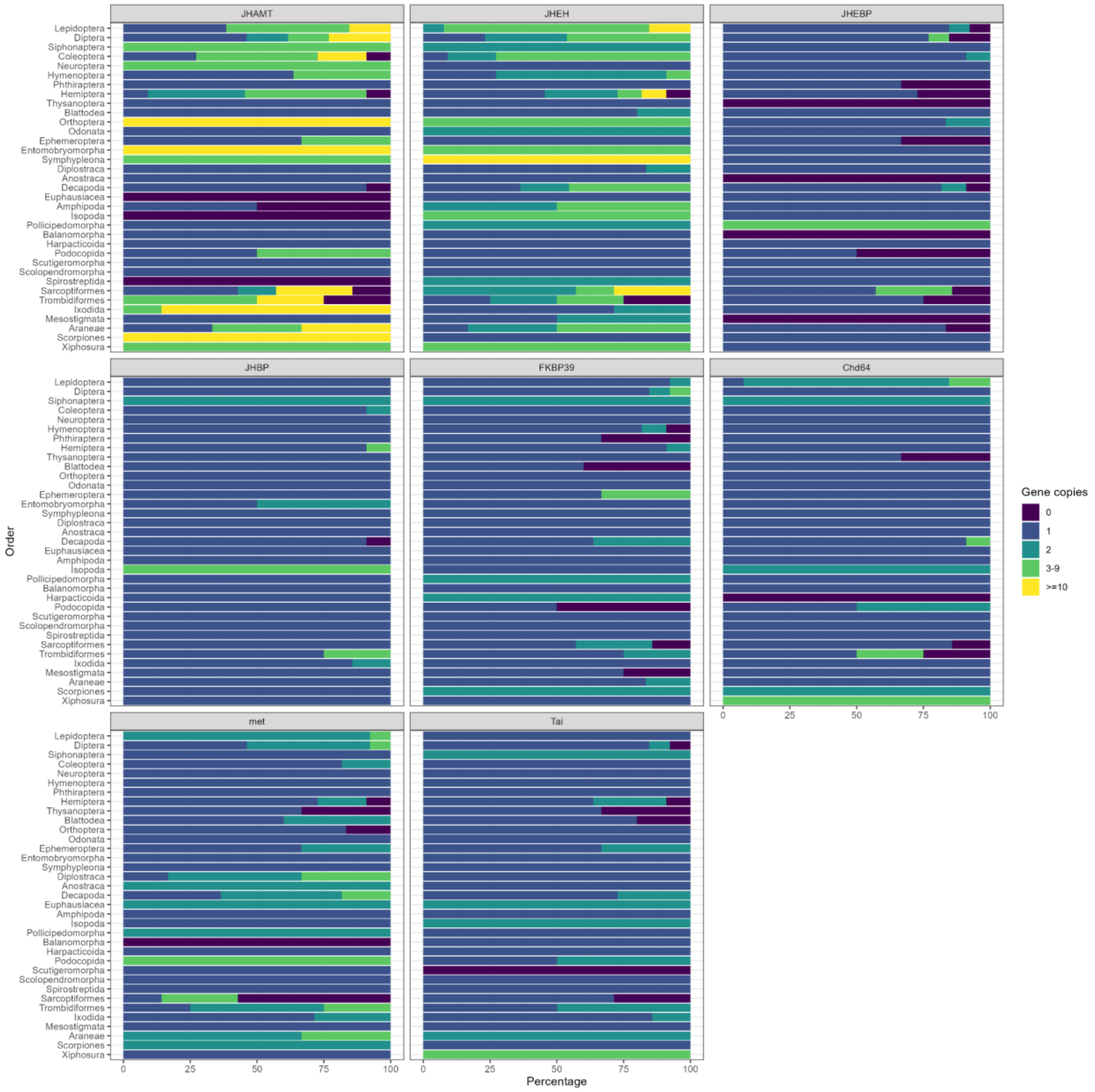

### Supplementary Figure 4: Early gene module

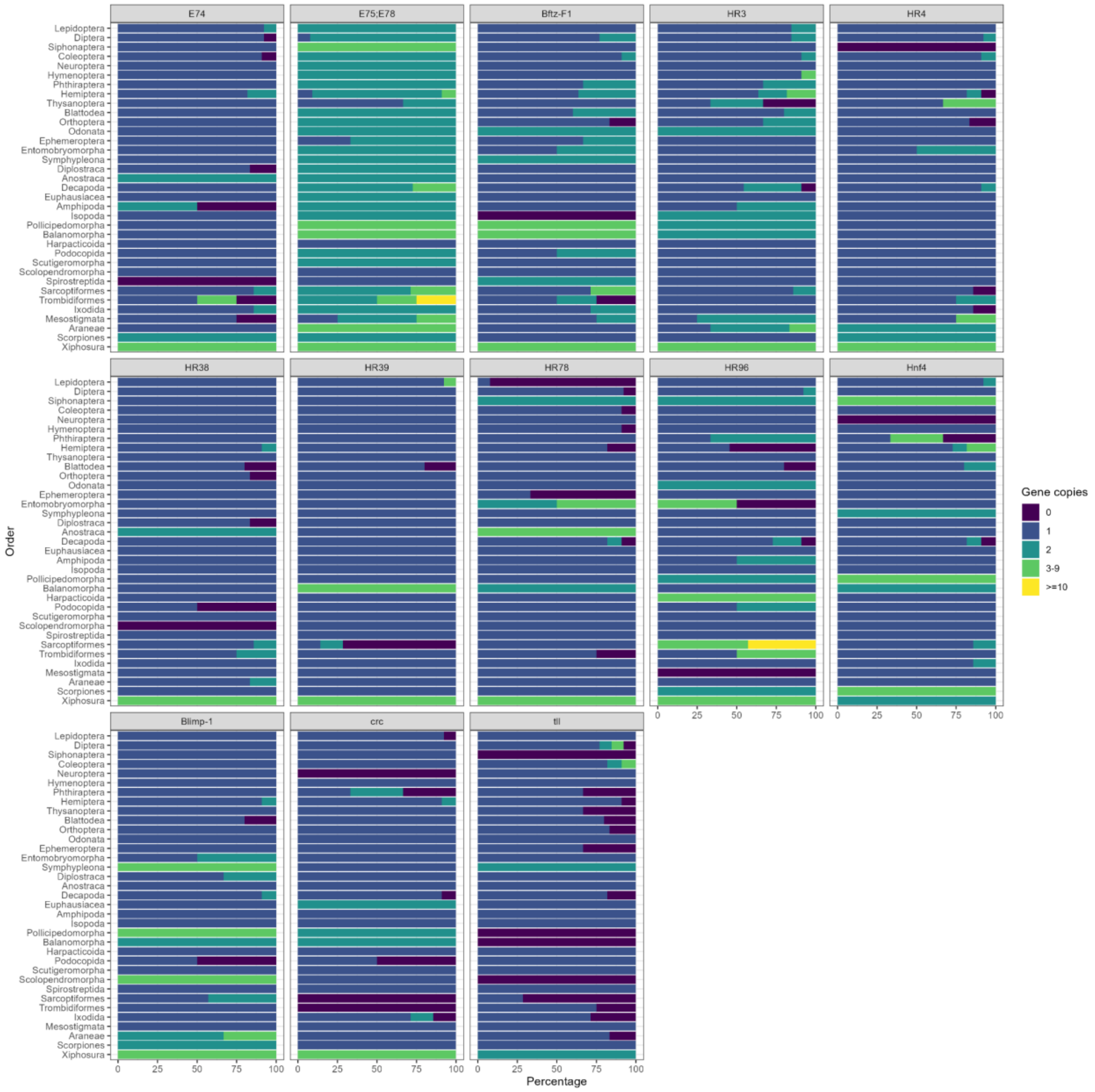

### Supplementary Figure 5: Fate gene module

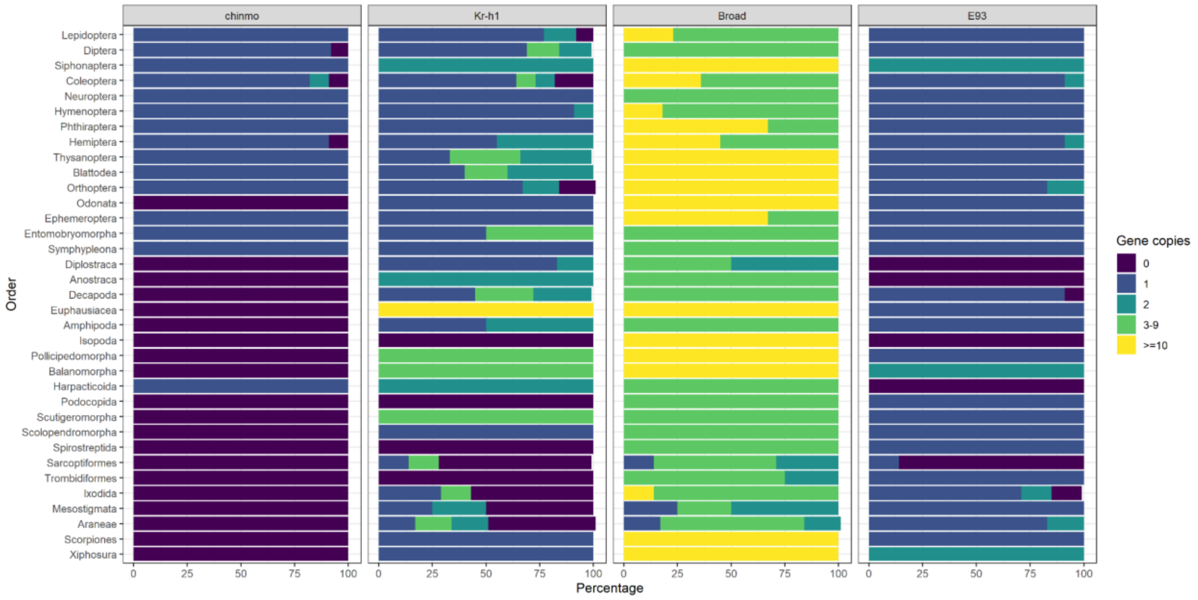

### Supplementary Figure 6: Neurostimuli reception module

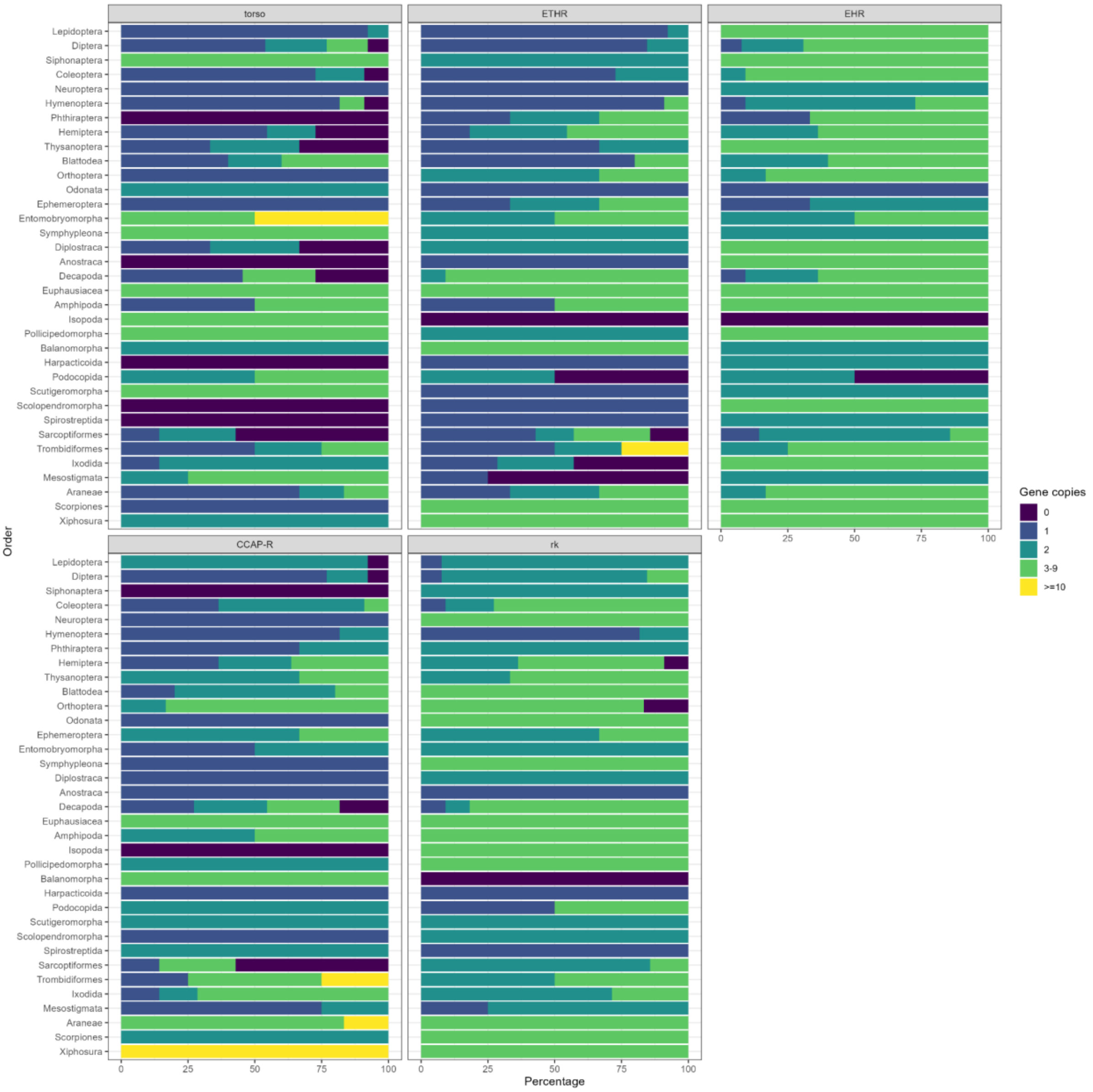

### Supplementary Figure 7: Late gene module

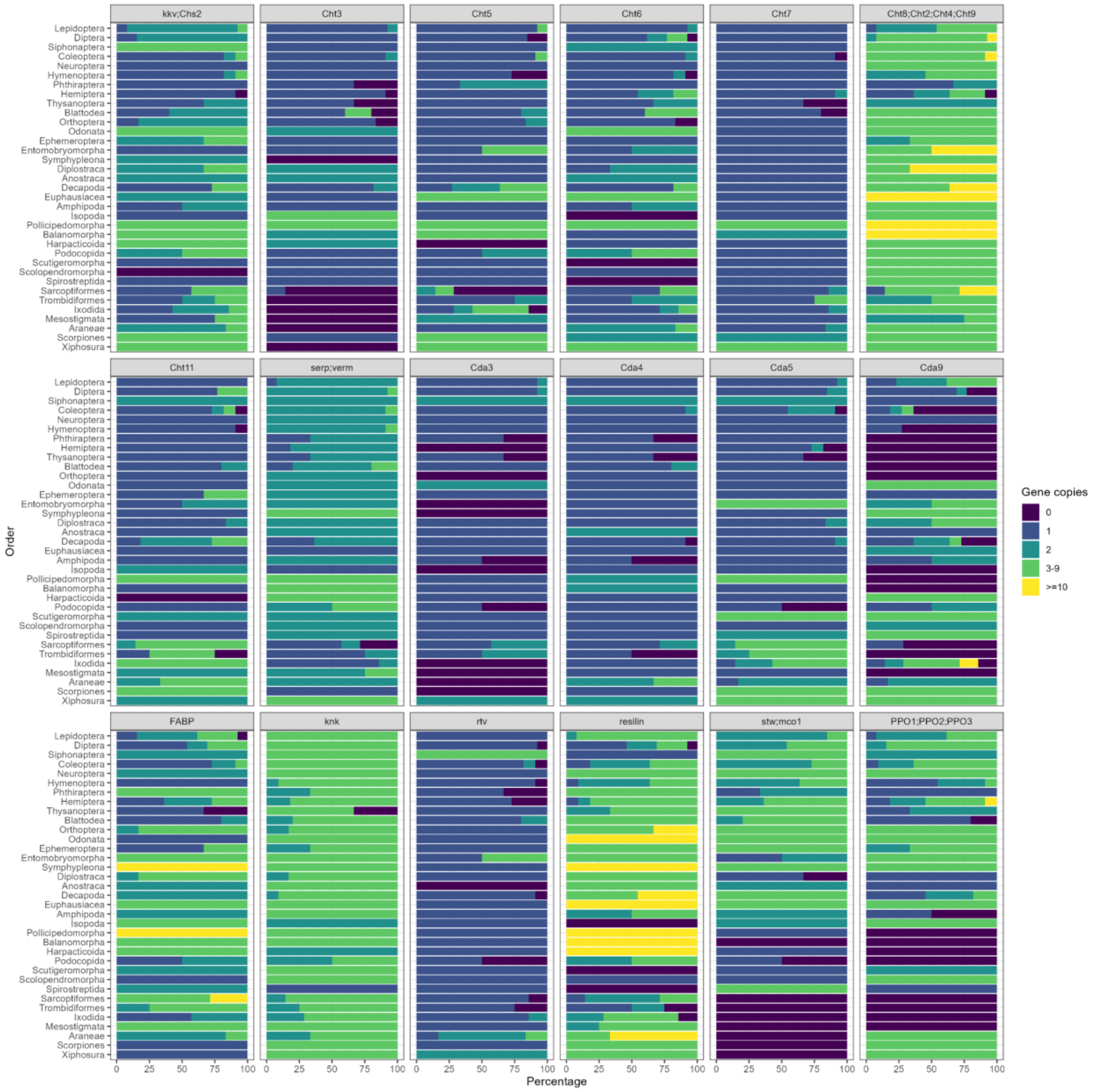

**Supplementary Figure 8: Ancestral reconstruction of protein domain architecture rearrangements**

**A)** Proportions of inferred domain architecture rearrangement are shown (y-axis) with each of the solution types in the two sets coloured according to the legend, based on Dohmen et al. 2020. Conserved unchanged domain architectures are categorised as maintained, observed rearrangements can be classified as complex solutions, where an explanation cannot be inferred, ambiguous solutions where different event types can explain the new rearrangement, non-ambiguous solutions classifies rearrangements from the same event type and exact solutions infers univocal event types. **B)** Cartoon modified from DomRates User Manual found at https://zivgitlab.uni-muenster.de/domain-world/DomRates/, depicting changes in domain architecture in the child node from the parental node, under each of the different six domain rearrangement scenarios (fission, fusion, terminal loss, terminal emergence, single domain loss and single domain emergence).

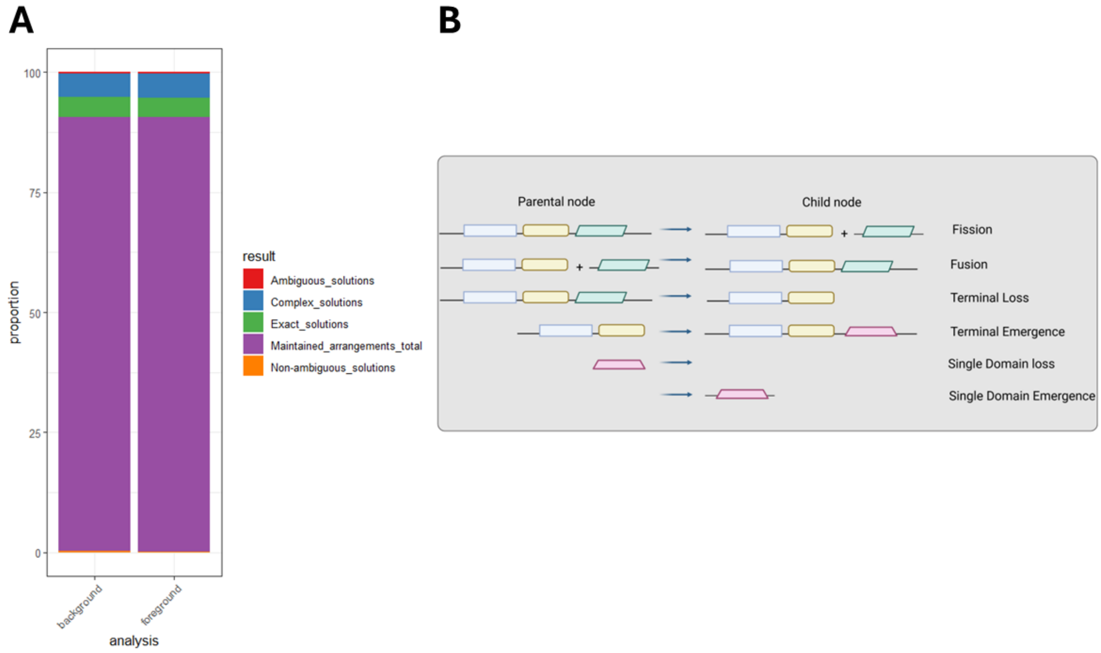


**Legend: Supplementary Figures 9-12. Species-tree gene-tree reconciliations of gene families with deep time duplications**

Dendrograms represent species-tree gene-tree reconciliations for examined gene families of interest, such as serpentine;vermiform (serp;verm, Supplementary Figure 9), chitin deacetylases 4 (Cda4, Supplementary Figure 10), krotzkopf verkehrt and chitin synthase 2 (kkv;chs2, Supplementary Figure 11), knickkopf (knk, Supplementary Figure 12). Bootstrap support values larger than 90% from the original gene tree are reported as node labels. Mapped evolutionary events are coloured according to legend: when in grey, branches, nodes and labels refer to losses, while when in black, they indicate sequences retained after speciation, orange nodes represent duplication events.

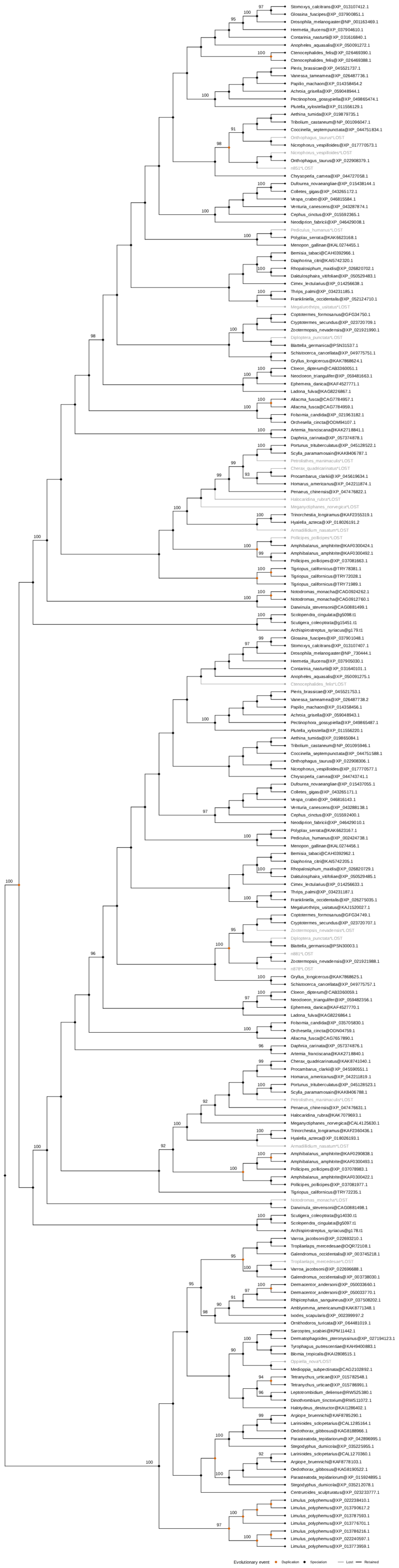

Stomoxys_calcitrans@XP_013107412.1
Glossina_fuscipes@XP_037900851.1
Drosophila_melanogaster@NP_001163469.1
Hermetia_illucens@XP_037904610.1
Contarinia_nasturtii@XP_031616840.1
Anopheles_aquasalis@XP_050091272.1
Ctenocephalides_felis@XP_026469390.1
Ctenocephalides_felis@XP_026469388.1
Pieris_brassicae@XP_045521737.1
Vanessa_tameamea@XP_026487736.1
Papilio_machaon@XP_014358454.2
Achroia_grisella@XP_059048944.1
Pectinophora_gossypiella@XP_049865474.1
Plutella_xylostella@XP_011556129.1
Aethina_tumida@XP_019879735.1
Tribolium_castaneum@NP_001096047.1
Coccinella_septempunctata@XP_044751834.1
Onthophagus_taurus*LOST
Nicrophorus_vespilloides@XP_017770573.1
Nicrophorus_vespilloides*LOST
Onthophagus_taurus@XP_022908379.1
n851*LOST
Chrysoperla_carnea@XP_044727058.1
Dufourea_novaeangliae@XP_015438144.1
Colletes_gigas@XP_043265172.1
Vespa_crabro@XP_046815584.1
Venturia_canescens@XP_043287874.1
Cephus_cinctus@XP_015592365.1
Neodiprion_fabricii@XP_046429008.1
Pediculus_humanus*LOST
Polyplax_serrata@KAK6623168.1
Menopon_gallinae@KAL0274455.1
Bemisia_tabaci@CAH0392966.1
Diaphorina_citri@KAI5742320.1
Rhopalosiphum_maidis@XP_026820702.1
Daktulosphaira_vitifoliae@XP_050529483.1
Cimex_lectularius@XP_014256638.1
Thrips_palmi@XP_034231185.1
Frankliniella_occidentalis@XP_052124710.1
Megalurothrips_usitatus*LOST
Coptotermes_formosanus@GFG34750.1
Cryptotermes_secundus@XP_023720709.1
Zootermopsis_nevadensis@XP_021921990.1
Diploptera_punctata*LOST
Blattella_germanica@PSN31537.1
Schistocerca_cancellata@XP_049775751.1
Gryllus_longicercus@KAK7868624.1
Cloeon_dipterum@CAB3360051.1
Neocloeon_triangulifer@XP_059481663.1
Ephemera_danica@KAF4527771.1
Ladona_fulva@KAG8226867.1
Allacma_fusca@CAG7784957.1
Allacma_fusca@CAG7784959.1
Folsomia_candida@XP_021963182.1
Orchesella_cincta@ODM94107.1
Artemia_franciscana@KAK2718841.1
Daphnia_carinata@XP_057374878.1
Portunus_trituberculatus@XP_045128522.1
Scylla_paramamosain@KAK8406787.1
Petrolisthes_manimaculis*LOST
Cherax_quadricarinatus*LOST
Procambarus_clarkii@XP_045619634.1
Homarus_americanus@XP_042211874.1
Penaeus_chinensis@XP_047476822.1
Halocaridina_rubra*LOST
Meganyctiphanes_norvegica*LOST
Trinorchestia_longiramus@KAF2355319.1
Hyalella_azteca@XP_018026191.2
Armadillidium_nasatum*LOST
Pollicipes_pollicipes*LOST
Amphibalanus_amphitrite@KAF0300424.1
Amphibalanus_amphitrite@KAF0300492.1
Pollicipes_pollicipes@XP_037081663.1
Tigriopus_californicus@TRY78381.1
Tigriopus_californicus@TRY72028.1
Tigriopus_californicus@TRY71989.1
Notodromas_monacha@CAG0924262.1
Notodromas_monacha@CAG0912760.1
Darwinula_stevensoni@CAG0881499.1
Scolopendra_cingulata@g5098.t1
Scutigera_coleoptrata@g15451.t1
Archispirostreptus_syriacus@g179.t1
Glossina_fuscipes@XP_037901048.1
Stomoxys_calcitrans@XP_013107407.1
Drosophila_melanogaster@NP_730444.1
Hermetia_illucens@XP_037905030.1
Contarinia_nasturtii@XP_031640101.1
Anopheles_aquasalis@XP_050091275.1
Ctenocephalides_felis*LOST
Pieris_brassicae@XP_045521753.1
Vanessa_tameamea@XP_026487738.2
Papilio_machaon@XP_014358456.1
Achroia_grisella@XP_059048943.1
Pectinophora_gossypiella@XP_049865487.1
Plutella_xylostella@XP_011556220.1
Aethina_tumida@XP_019865084.1
Tribolium_castaneum@NP_001095946.1
Coccinella_septempunctata@XP_044751588.1
Onthophagus_taurus@XP_022908306.1
Nicrophorus_vespilloides@XP_017770577.1
Chrysoperla_carnea@XP_044743741.1
Dufourea_novaeangliae@XP_015437055.1
Colletes_gigas@XP_043265171.1
Vespa_crabro@XP_046816143.1
Venturia_canescens@XP_043288138.1
Cephus_cinctus@XP_015592400.1
Neodiprion_fabricii@XP_046429010.1
Polyplax_serrata@KAK6623167.1
Pediculus_humanus@XP_002424738.1
Menopon_gallinae@KAL0274456.1
Bemisia_tabaci@CAH0392962.1
Diaphorina_citri@KAI5742205.1
Rhopalosiphum_maidis@XP_026820729.1
Daktulosphaira_vitifoliae@XP_050529485.1
Cimex_lectularius@XP_014256633.1
Thrips_palmi@XP_034231187.1
Frankliniella_occidentalis@XP_026275035.1
Megalurothrips_usitatus@KAJ1520027.1
Coptotermes_formosanus@GFG34749.1
Cryptotermes_secundus@XP_023720707.1
Zootermopsis_nevadensis*LOST
Diploptera_punctata*LOST
Blattella_germanica@PSN30003.1
n881*LOST
Zootermopsis_nevadensis@XP_021921988.1
n878*LOST
Gryllus_longicercus@KAK7868625.1
Schistocerca_cancellata@XP_049775757.1
Cloeon_dipterum@CAB3360059.1
Neocloeon_triangulifer@XP_059482356.1
Ephemera_danica@KAF4527770.1
Ladona_fulva@KAG8226864.1
Folsomia_candida@XP_035705830.1
Orchesella_cincta@ODN04759.1
Allacma_fusca@CAG7657890.1
Daphnia_carinata@XP_057374876.1
Artemia_franciscana@KAK2718840.1
Cherax_quadricarinatus@KAK8741040.1
Procambarus_clarkii@XP_045590551.1
Homarus_americanus@XP_042211819.1
Portunus_trituberculatus@XP_045128523.1
Scylla_paramamosain@KAK8406788.1
Petrolisthes_manimaculis*LOST
Penaeus_chinensis@XP_047476631.1
Halocaridina_rubra@KAK7079693.1
Meganyctiphanes_norvegica@CAL4125630.1
Trinorchestia_longiramus@KAF2360436.1
Hyalella_azteca@XP_018026193.1
Armadillidium_nasatum*LOST
Amphibalanus_amphitrite@KAF0290838.1
Amphibalanus_amphitrite@KAF0300493.1
Pollicipes_pollicipes@XP_037078983.1
Amphibalanus_amphitrite@KAF0300422.1
Pollicipes_pollicipes@XP_037081977.1
Tigriopus_californicus@TRY72235.1
Notodromas_monacha*LOST
Darwinula_stevensoni@CAG0881498.1
Scutigera_coleoptrata@g14030.t1
Scolopendra_cingulata@g5097.t1
Archispirostreptus_syriacus@g178.t1
Varroa_jacobsoni@XP_022693210.1
Tropilaelaps_mercedesae@OQR72108.1
Galendromus_occidentalis@XP_003745218.1
Tropilaelaps_mercedesae*LOST
Varroa_jacobsoni@XP_022696688.1
Galendromus_occidentalis@XP_003738030.1
Dermacentor_andersoni@XP_050033660.1
Dermacentor_andersoni@XP_050033770.1
Rhipicephalus_sanguineus@XP_037508202.1
Amblyomma_americanum@KAK8771348.1
Ixodes_scapularis@XP_002399997.2
Ornithodoros_turicata@XP_064481019.1
Sarcoptes_scabiei@KPM11442.1
Dermatophagoides_pteronyssinus@XP_027194123.1
Tyrophagus_putrescentiae@KAH9400883.1
Blomia_tropicalis@KAI2808515.1
Oppiella_nova*LOST
Medioppia_subpectinata@CAG2102892.1
Tetranychus_urticae@XP_015782548.1
Tetranychus_urticae@XP_015786991.1
Leptotrombidium_deliense@RWS25380.1
Dinothrombium_tinctorium@RWS11072.1
Halotydeus_destructor@KAI1286402.1
Argiope_bruennichi@KAF8785290.1
Larinioides_sclopetarius@CAL1285164.1
Oedothorax_gibbosus@KAG8188966.1
Parasteatoda_tepidariorum@XP_042896995.1
Stegodyphus_dumicola@XP_035225955.1
Larinioides_sclopetarius@CAL1270360.1
Argiope_bruennichi@KAF8778103.1
Oedothorax_gibbosus@KAG8190522.1
Parasteatoda_tepidariorum@XP_015924895.1
Stegodyphus_dumicola@XP_035212078.1
Centruroides_sculpturatus@XP_023233777.1
Limulus_polyphemus@XP_022238410.1
Limulus_polyphemus@XP_013790617.2
Limulus_polyphemus@XP_013787593.1
Limulus_polyphemus@XP_013776701.1
Limulus_polyphemus@XP_013786216.1
Limulus_polyphemus@XP_022240597.1
Limulus_polyphemus@XP_013773959.1
Evolutionary event
Duplication
Speciation
Lost
Retained

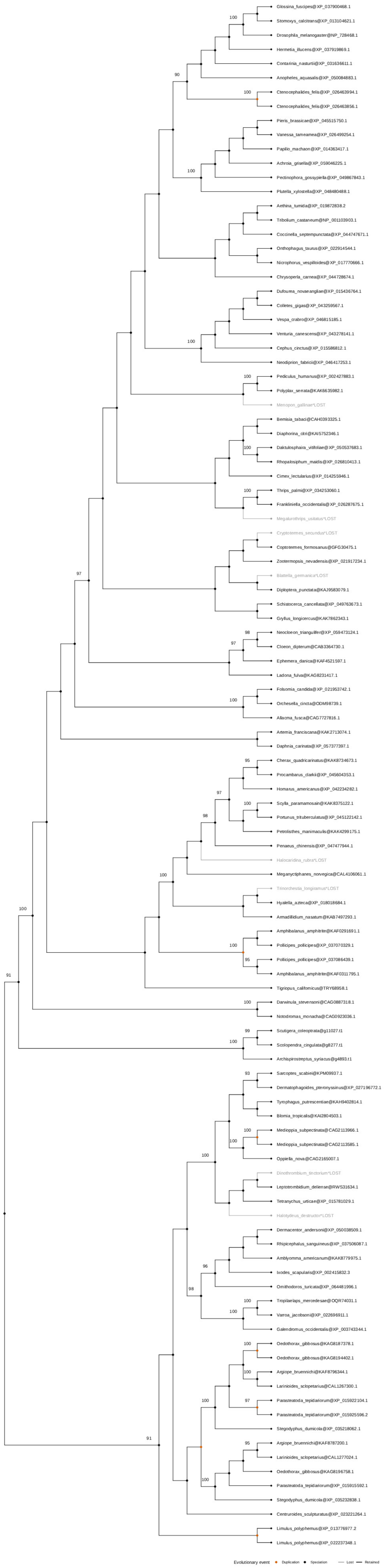
Glossina_fuscipes@XP_037900468.1
Stomoxys_calcitrans@XP_013104621.1
Drosophila_melanogaster@NP_728468.1
Hermetia_illucens@XP_037919869.1
Contarinia_nasturtii@XP_031636611.1
Anopheles_aquasalis@XP_050084883.1
Ctenocephalides_felis@XP_026463994.1
Ctenocephalides_felis@XP_026463856.1
Pieris_brassicae@XP_045515750.1
Vanessa_tameamea@XP_026499254.1
Papilio_machaon@XP_014363417.1
Achroia_grisella@XP_059046225.1
Pectinophora_gossypiella@XP_049867843.1
Plutella_xylostella@XP_048480488.1
Aethina_tumida@XP_019872838.2
Tribolium_castaneum@NP_001103903.1
Coccinella_septempunctata@XP_044747671.1
Onthophagus_taurus@XP_022914544.1
Nicrophorus_vespilloides@XP_017770666.1
Chrysoperla_carnea@XP_044728674.1
Dufourea_novaeangliae@XP_015436764.1
Colletes_gigas@XP_043259567.1
Vespa_crabro@XP_046815185.1
Venturia_canescens@XP_043278141.1
Cephus_cinctus@XP_015586812.1
Neodiprion_fabricii@XP_046417253.1
Pediculus_humanus@XP_002427883.1
Polyplax_serrata@KAK6635982.1
Menopon_gallinae*LOST
Bemisia_tabaci@CAH0393325.1
Diaphorina_citri@KAI5752346.1
Daktulosphaira_vitifoliae@XP_050537683.1
Rhopalosiphum_maidis@XP_026810413.1
Cimex_lectularius@XP_014255946.1
Thrips_palmi@XP_034253060.1
Frankliniella_occidentalis@XP_026287675.1
Megalurothrips_usitatus*LOST
Cryptotermes_secundus*LOST
Coptotermes_formosanus@GFG30475.1
Zootermopsis_nevadensis@XP_021917234.1
Blattella_germanica*LOST
Diploptera_punctata@KAJ9583079.1
Schistocerca_cancellata@XP_049763673.1
Gryllus_longicercus@KAK7862343.1
Neocloeon_triangulifer@XP_059473124.1
Cloeon_dipterum@CAB3364730.1
Ephemera_danica@KAF4521597.1
Ladona_fulva@KAG8231417.1
Folsomia_candida@XP_021953742.1
Orchesella_cincta@ODM98739.1
Allacma_fusca@CAG7727816.1
Artemia_franciscana@KAK2713074.1
Daphnia_carinata@XP_057377397.1
Cherax_quadricarinatus@KAK8734673.1
Procambarus_clarkii@XP_045604353.1
Homarus_americanus@XP_042234282.1
Scylla_paramamosain@KAK8375122.1
Portunus_trituberculatus@XP_045122142.1
Petrolisthes_manimaculis@KAK4299175.1
Penaeus_chinensis@XP_047477944.1
Halocaridina_rubra*LOST
Meganyctiphanes_norvegica@CAL4106061.1
Trinorchestia_longiramus*LOST
Hyalella_azteca@XP_018018684.1
Armadillidium_nasatum@KAB7497293.1
Amphibalanus_amphitrite@KAF0291691.1
Pollicipes_pollicipes@XP_037070329.1
Pollicipes_pollicipes@XP_037086439.1
Amphibalanus_amphitrite@KAF0311795.1
Tigriopus_californicus@TRY68958.1
Darwinula_stevensoni@CAG0887318.1
Notodromas_monacha@CAG0923036.1
Scutigera_coleoptrata@g11027.t1
Scolopendra_cingulata@g8277.t1
Archispirostreptus_syriacus@g4893.t1
Sarcoptes_scabiei@KPM09937.1
Dermatophagoides_pteronyssinus@XP_027196772.1
Tyrophagus_putrescentiae@KAH9402814.1
Blomia_tropicalis@KAI2804503.1
Medioppia_subpectinata@CAG2113966.1
Medioppia_subpectinata@CAG2113585.1
Oppiella_nova@CAG2165007.1
Dinothrombium_tinctorium*LOST
Leptotrombidium_deliense@RWS31634.1
Tetranychus_urticae@XP_015781029.1
Halotydeus_destructor*LOST
Dermacentor_andersoni@XP_050038509.1
Rhipicephalus_sanguineus@XP_037506087.1
Amblyomma_americanum@KAK8779975.1
Ixodes_scapularis@XP_002415832.3
Ornithodoros_turicata@XP_064481996.1
Tropilaelaps_mercedesae@OQR74031.1
Varroa_jacobsoni@XP_022696911.1
Galendromus_occidentalis@XP_003743344.1
Oedothorax_gibbosus@KAG8187378.1
Oedothorax_gibbosus@KAG8194402.1
Argiope_bruennichi@KAF8796344.1
Larinioides_sclopetarius@CAL1267300.1
Parasteatoda_tepidariorum@XP_015922104.1
Parasteatoda_tepidariorum@XP_015925596.2
Stegodyphus_dumicola@XP_035218062.1
Argiope_bruennichi@KAF8787200.1
Larinioides_sclopetarius@CAL1277024.1
Oedothorax_gibbosus@KAG8196758.1
Parasteatoda_tepidariorum@XP_015915592.1
Stegodyphus_dumicola@XP_035232838.1
Centruroides_sculpturatus@XP_023221264.1
Limulus_polyphemus@XP_013776977.2
Limulus_polyphemus@XP_022237348.1
Evolutionary event
Duplication
Speciation
Lost
Retained

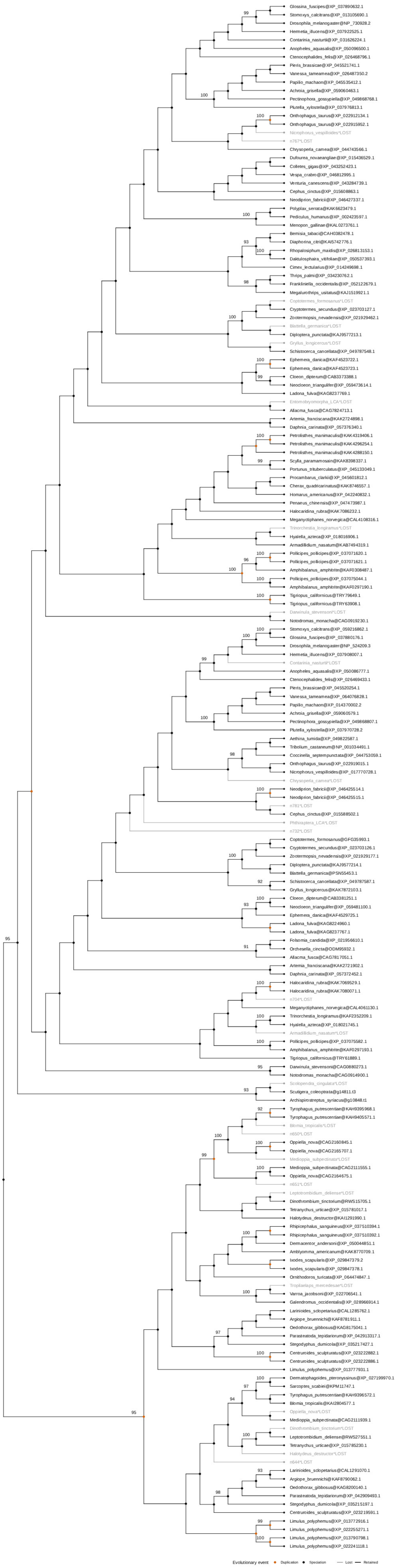
Glossina_fuscipes@XP_037890632.1
Stomoxys_calcitrans@XP_013105690.1
Drosophila_melanogaster@NP_730928.2
Hermetia_illucens@XP_037922525.1
Contarinia_nasturtii@XP_031626224.1
Anopheles_aquasalis@XP_050096500.1
Ctenocephalides_felis@XP_026468796.1
Pieris_brassicae@XP_045521741.1
Vanessa_tameamea@XP_026487350.2
Papilio_machaon@XP_045535412.1
Achroia_grisella@XP_059060463.1
Pectinophora_gossypiella@XP_049868768.1
Plutella_xylostella@XP_037976813.1
Onthophagus_taurus@XP_022912134.1
Onthophagus_taurus@XP_022915952.1
Nicrophorus_vespilloides*LOST
n767*LOST
Chrysoperla_carnea@XP_044743566.1
Dufourea_novaeangliae@XP_015436529.1
Colletes_gigas@XP_043252423.1
Vespa_crabro@XP_046812995.1
Venturia_canescens@XP_043284739.1
Cephus_cinctus@XP_015608863.1
Neodiprion_fabricii@XP_046427337.1
Polyplax_serrata@KAK6623479.1
Pediculus_humanus@XP_002423597.1
Menopon_gallinae@KAL0273761.1
Bemisia_tabaci@CAH0382478.1
Diaphorina_citri@KAI5742776.1
Rhopalosiphum_maidis@XP_026813153.1
Daktulosphaira_vitifoliae@XP_050537393.1
Cimex_lectularius@XP_014249698.1
Thrips_palmi@XP_034230762.1
Frankliniella_occidentalis@XP_052122679.1
Megalurothrips_usitatus@KAJ1519921.1
Coptotermes_formosanus*LOST
Cryptotermes_secundus@XP_023703127.1
Zootermopsis_nevadensis@XP_021929462.1
Blattella_germanica*LOST
Diploptera_punctata@KAJ9577213.1
Gryllus_longicercus*LOST
Schistocerca_cancellata@XP_049787548.1
Ephemera_danica@KAF4523722.1
Ephemera_danica@KAF4523723.1
Cloeon_dipterum@CAB3373388.1
Neocloeon_triangulifer@XP_059473614.1
Ladona_fulva@KAG8237769.1
Entomobryomorpha_LCA*LOST
Allacma_fusca@CAG7824713.1
Artemia_franciscana@KAK2724898.1
Daphnia_carinata@XP_057376340.1
Petrolisthes_manimaculis@KAK4319406.1
Petrolisthes_manimaculis@KAK4296254.1
Petrolisthes_manimaculis@KAK4288150.1
Scylla_paramamosain@KAK8398337.1
Portunus_trituberculatus@XP_045133049.1
Procambarus_clarkii@XP_045601812.1
Cherax_quadricarinatus@KAK8746557.1
Homarus_americanus@XP_042240832.1
Penaeus_chinensis@XP_047473987.1
Halocaridina_rubra@KAK7086232.1
Meganyctiphanes_norvegica@CAL4108316.1
Trinorchestia_longiramus*LOST
Hyalella_azteca@XP_018016906.1
Armadillidium_nasatum@KAB7494319.1
Pollicipes_pollicipes@XP_037071620.1
Pollicipes_pollicipes@XP_037071621.1
Amphibalanus_amphitrite@KAF0308487.1
Pollicipes_pollicipes@XP_037075044.1
Amphibalanus_amphitrite@KAF0297190.1
Tigriopus_californicus@TRY79649.1
Tigriopus_californicus@TRY63908.1
Darwinula_stevensoni*LOST
Notodromas_monacha@CAG0919230.1
Stomoxys_calcitrans@XP_059216862.1
Glossina_fuscipes@XP_037880176.1
Drosophila_melanogaster@NP_524209.3
Hermetia_illucens@XP_037908007.1
Contarinia_nasturtii*LOST
Anopheles_aquasalis@XP_050086777.1
Ctenocephalides_felis@XP_026469433.1
Pieris_brassicae@XP_045520254.1
Vanessa_tameamea@XP_064076828.1
Papilio_machaon@XP_014370002.2
Achroia_grisella@XP_059060579.1
Pectinophora_gossypiella@XP_049868807.1
Plutella_xylostella@XP_037970728.2
Aethina_tumida@XP_049822587.1
Tribolium_castaneum@NP_001034491.1
Coccinella_septempunctata@XP_044753059.1
Onthophagus_taurus@XP_022919015.1
Nicrophorus_vespilloides@XP_017770728.1
Chrysoperla_carnea*LOST
Neodiprion_fabricii@XP_046425514.1
Neodiprion_fabricii@XP_046425515.1
n781*LOST
Cephus_cinctus@XP_015588502.1
Phthiraptera_LCA*LOST
n732*LOST
Coptotermes_formosanus@GFG35993.1
Cryptotermes_secundus@XP_023703126.1
Zootermopsis_nevadensis@XP_021929177.1
Diploptera_punctata@KAJ9577214.1
Blattella_germanica@PSN55453.1
Schistocerca_cancellata@XP_049787587.1
Gryllus_longicercus@KAK7872103.1
Cloeon_dipterum@CAB3381251.1
Neocloeon_triangulifer@XP_059481100.1
Ephemera_danica@KAF4529725.1
Ladona_fulva@KAG8224960.1
Ladona_fulva@KAG8237767.1
Folsomia_candida@XP_021956610.1
Orchesella_cincta@ODM95932.1
Allacma_fusca@CAG7817051.1
Artemia_franciscana@KAK2721902.1
Daphnia_carinata@XP_057372452.1
Halocaridina_rubra@KAK7069529.1
Halocaridina_rubra@KAK7080071.1
n704*LOST
Meganyctiphanes_norvegica@CAL4061130.1
Trinorchestia_longiramus@KAF2352209.1
Hyalella_azteca@XP_018021745.1
Armadillidium_nasatum*LOST
Pollicipes_pollicipes@XP_037075582.1
Amphibalanus_amphitrite@KAF0297193.1
Tigriopus_californicus@TRY61889.1
Darwinula_stevensoni@CAG0880273.1
Notodromas_monacha@CAG0914900.1
Scolopendra_cingulata*LOST
Scutigera_coleoptrata@g14811.t3
Archispirostreptus_syriacus@g10848.t1
Tyrophagus_putrescentiae@KAH9395968.1
Tyrophagus_putrescentiae@KAH9405571.1
Blomia_tropicalis*LOST
n650*LOST
Oppiella_nova@CAG2160845.1
Oppiella_nova@CAG2165707.1
Medioppia_subpectinata*LOST
Medioppia_subpectinata@CAG2111555.1
Oppiella_nova@CAG2164675.1
n651*LOST
Leptotrombidium_deliense*LOST
Dinothrombium_tinctorium@RWS15705.1
Tetranychus_urticae@XP_015781017.1
Halotydeus_destructor@KAI1291990.1
Rhipicephalus_sanguineus@XP_037510394.1
Rhipicephalus_sanguineus@XP_037510392.1
Dermacentor_andersoni@XP_050044851.1
Amblyomma_americanum@KAK8770709.1
Ixodes_scapularis@XP_029847379.2
Ixodes_scapularis@XP_029847378.1
Ornithodoros_turicata@XP_064474847.1
Tropilaelaps_mercedesae*LOST
Varroa_jacobsoni@XP_022706541.1
Galendromus_occidentalis@XP_028966914.1
Larinioides_sclopetarius@CAL1285762.1
Argiope_bruennichi@KAF8781911.1
Oedothorax_gibbosus@KAG8175041.1
Parasteatoda_tepidariorum@XP_042913317.1
Stegodyphus_dumicola@XP_035217427.1
Centruroides_sculpturatus@XP_023222882.1
Centruroides_sculpturatus@XP_023222886.1
Limulus_polyphemus@XP_013777931.1
Dermatophagoides_pteronyssinus@XP_027199970.1
Sarcoptes_scabiei@KPM11747.1
Tyrophagus_putrescentiae@KAH9396572.1
Blomia_tropicalis@KAI2804577.1
Oppiella_nova*LOST
Medioppia_subpectinata@CAG2111939.1
Dinothrombium_tinctorium*LOST
Leptotrombidium_deliense@RWS27551.1
Tetranychus_urticae@XP_015785230.1
Halotydeus_destructor*LOST
n644*LOST
Larinioides_sclopetarius@CAL1291070.1
Argiope_bruennichi@KAF8790062.1
Oedothorax_gibbosus@KAG8200140.1
Parasteatoda_tepidariorum@XP_042909493.1
Stegodyphus_dumicola@XP_035215197.1
Centruroides_sculpturatus@XP_023219591.1
Limulus_polyphemus@XP_013772916.1
Limulus_polyphemus@XP_022255271.1
Limulus_polyphemus@XP_013790798.1
Limulus_polyphemus@XP_022241118.1
Evolutionary event
Duplication
Speciation
Lost
Retained

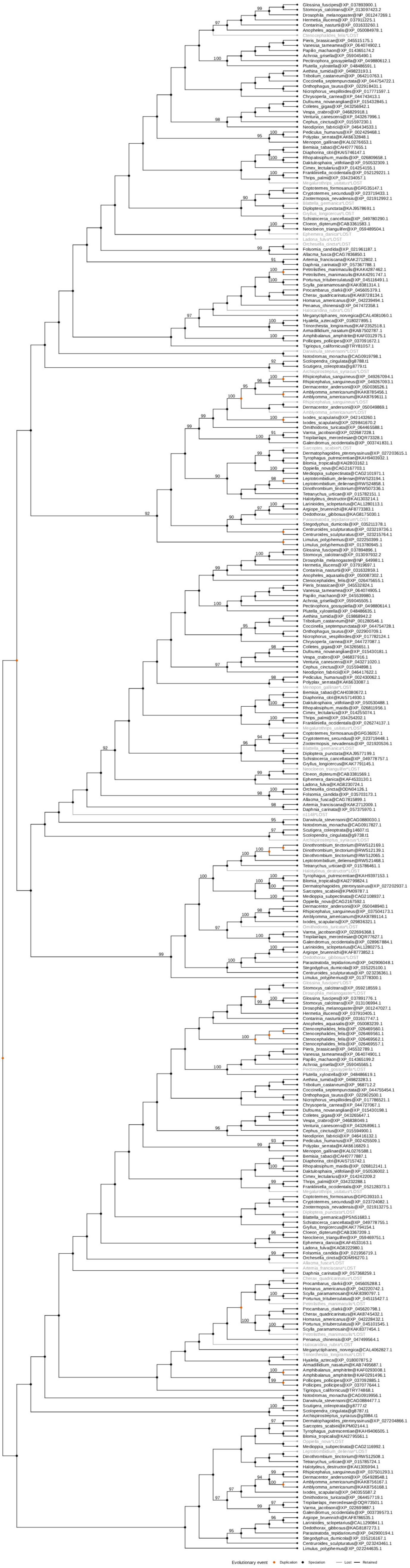
Evolutionary event
Duplication
Speciation
Lost
Retained

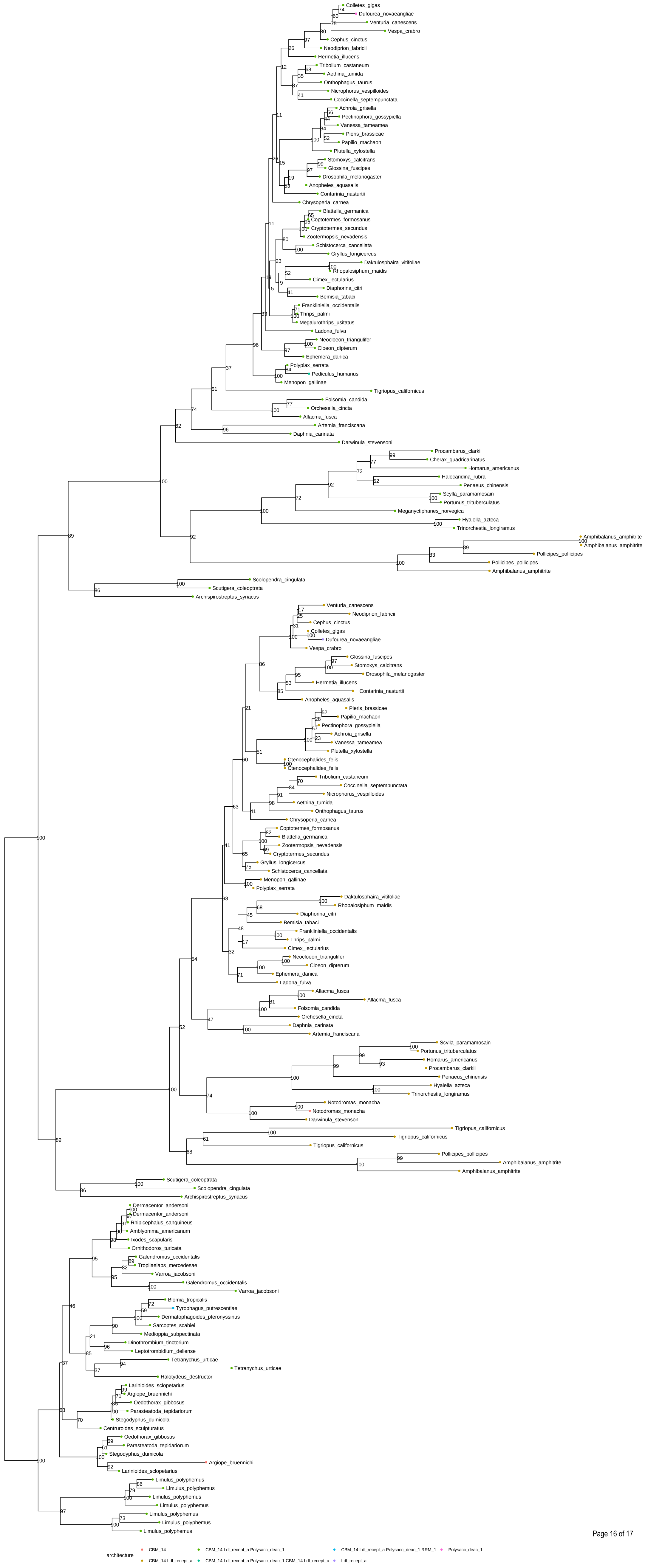
Colletes_gigas
Dufourea_novaeangliae
Venturia_canescens
Vespa_crabro
Cephus_cinctus
Neodiprion_fabricii
Hermetia_illucens
Tribolium_castaneum
Aethina_tumida
Onthophagus_taurus
Nicrophorus_vespilloides
Coccinella_septempunctata
Achroia_grisella
Pectinophora_gossypiella
Vanessa_tameamea
Pieris_brassicae
Papilio_machaon
Plutella_xylostella
Stomoxys_calcitrans
Glossina_fuscipes
Drosophila_melanogaster
Anopheles_aquasalis
Contarinia_nasturtii
Chrysoperla_carnea
Blattella_germanica
Coptotermes_formosanus
Cryptotermes_secundus
Zootermopsis_nevadensis
Schistocerca_cancellata
Gryllus_longicercus
Daktulosphaira_vitifoliae
Rhopalosiphum_maidis
Cimex_lectularius
Diaphorina_citri
Bemisia_tabaci
Frankliniella_occidentalis
Thrips_palmi
Megalurothrips_usitatus
Ladona_fulva
Neocloeon_triangulifer
Cloeon_dipterum
Ephemera_danica
Polyplax_serrata
Pediculus_humanus
Menopon_gallinae
Tigriopus_californicus
Folsomia_candida
Orchesella_cincta
Allacma_fusca
Artemia_franciscana
Daphnia_carinata
Darwinula_stevensoni
Procambarus_clarkii
Cherax_quadricarinatus
Homarus_americanus
Halocaridina_rubra
Penaeus_chinensis
Scylla_paramamosain
Portunus_trituberculatus
Meganyctiphanes_norvegica
Hyalella_azteca
Trinorchestia_longiramus
Amphibalanus_amphitrite
Amphibalanus_amphitrite
Pollicipes_pollicipes
Pollicipes_pollicipes
Amphibalanus_amphitrite
Scolopendra_cingulata
Scutigera_coleoptrata
Archispirostreptus_syriacus
Venturia_canescens
Neodiprion_fabricii
Cephus_cinctus
Colletes_gigas
Dufourea_novaeangliae
Vespa_crabro
Glossina_fuscipes
Stomoxys_calcitrans
Drosophila_melanogaster
Hermetia_illucens
Contarinia_nasturtii
Anopheles_aquasalis
Pieris_brassicae
Papilio_machaon
Pectinophora_gossypiella
Achroia_grisella
Vanessa_tameamea
Plutella_xylostella
Ctenocephalides_felis
Ctenocephalides_felis
Tribolium_castaneum
Coccinella_septempunctata
Nicrophorus_vespilloides
Aethina_tumida
Onthophagus_taurus
Chrysoperla_carnea
Coptotermes_formosanus
Blattella_germanica
Zootermopsis_nevadensis
Cryptotermes_secundus
Gryllus_longicercus
Schistocerca_cancellata
Menopon_gallinae
Polyplax_serrata
Daktulosphaira_vitifoliae
Rhopalosiphum_maidis
Diaphorina_citri
Bemisia_tabaci
Frankliniella_occidentalis
Thrips_palmi
Cimex_lectularius
Neocloeon_triangulifer
Cloeon_dipterum
Ephemera_danica
Ladona_fulva
Allacma_fusca
Allacma_fusca
Folsomia_candida
Orchesella_cincta
Daphnia_carinata
Artemia_franciscana
Scylla_paramamosain
Portunus_trituberculatus
Homarus_americanus
Procambarus_clarkii
Penaeus_chinensis
Hyalella_azteca
Trinorchestia_longiramus
Notodromas_monacha
Notodromas_monacha
Darwinula_stevensoni
Tigriopus_californicus
Tigriopus_californicus
Tigriopus_californicus
Pollicipes_pollicipes
Amphibalanus_amphitrite
Amphibalanus_amphitrite
Scutigera_coleoptrata
Scolopendra_cingulata
Archispirostreptus_syriacus
Dermacentor_andersoni
Dermacentor_andersoni
Rhipicephalus_sanguineus
Amblyomma_americanum
Ixodes_scapularis
Ornithodoros_turicata
Galendromus_occidentalis
Tropilaelaps_mercedesae
Varroa_jacobsoni
Galendromus_occidentalis
Varroa_jacobsoni
Blomia_tropicalis
Tyrophagus_putrescentiae
Dermatophagoides_pteronyssinus
Sarcoptes_scabiei
Medioppia_subpectinata
Dinothrombium_tinctorium
Leptotrombidium_deliense
Tetranychus_urticae
Tetranychus_urticae
Halotydeus_destructor
Larinioides_sclopetarius
Argiope_bruennichi
Oedothorax_gibbosus
Parasteatoda_tepidariorum
Stegodyphus_dumicola
Centruroides_sculpturatus
Oedothorax_gibbosus
Parasteatoda_tepidariorum
Stegodyphus_dumicola
Argiope_bruennichi
Larinioides_sclopetarius
Limulus_polyphemus
Limulus_polyphemus
Limulus_polyphemus
Limulus_polyphemus
Limulus_polyphemus
Limulus_polyphemus
Limulus_polyphemus
architecture
CBM_14
CBM_14 Ldl_recept_a
CBM_14 Ldl_recept_a Polysacc_deac_1
CBM_14 Ldl_recept_a Polysacc_deac_1 CBM_14 Ldl_recept_a
CBM_14 Ldl_recept_a Polysacc_deac_1 RRM_1
Ldl_recept_a
Polysacc_deac_1

### Supplementary Figure 13: Ancestral loss of the deacetylase domain in the CDA1/serp-CDA2/verm family

The gene tree reports bootstrap support values as internal node labels and domain architecture as leaf node colours, according to the legend. A complete domain architecture (CBM_14 , Ldl_recept_a and Polysacc_deac_1, in light green) has the chitin binding Peritrophin-A domain at the N-terminal (CBM_14, PF01607), followed by the low-density lipoprotein receptor domain class A (Ldl_recept_a, PF00057) and it includes the polysaccharide deacetylase domain at the C-terminal (Polysacc_deac_1, PF01522); this is only identified in the chelicerate species, in the CDA1/serp proteins and in the CDA2/verm myriapod, but it is not annotated in the proteins from the pancrustacean CDA2/verm (CBM_14, Ldl_recept_a in sand yellow).
