## Supplementary Tables for "Evolutionary dynamics of the arthropod moulting machinery"

| Assembly Accession | TaxId | Species | Arthropoda Complete BUSCO | Arthropoda Single BUSCO | Arthropoda Duplicated BUSCO | Number of protein coding genes | Arthropoda Complete BUSCO Proteomes | Arthropoda Single BUSCO Proteomes | Arthropoda Duplicated BUSCO Proteomes |
|---|---|---|---|---|---|---|---|---|---|
| GCA_000376725.2 | 123851 | Ladona fulva | 93.7 | 92.3 | 1.4 | 18816 | 78.2 | 76.6 | 1.6 |
| GCA_000507165.2 | 1049336 | Ephemera danica | 94.5 | 91.6 | 2.9 | 18562 | 85.8 | 84.3 | 1.5 |
| GCA_000611955.2 | 407821 | Stegodyphus mimosarum | 93.7 | 90.2 | 3.5 | 27135 | 80 | 78.1 | 1.9 |
| GCA_000767015.2 | 6954 | Dermatophagoides farinae | 84.8 | 83.6 | 1.2 | 15208 | 90 | 88.4 | 1.6 |
| GCA_000828355.1 | 52283 | Sarcoptes scabiei | 84.9 | 84.7 | 0.2 | 10473 | 80.9 | 80.5 | 0.4 |
| GCA_001187945.1 | 7375 | Lucilia cuprina | 98.6 | 97.6 | 1 | 14452 | 91 | 90 | 1 |
| GCA_001718145.1 | 48709 | Orchesella cincta | 93 | 88.4 | 4.6 | 20247 | 90 | 85.2 | 4.8 |
| GCA_002081605.1 | 418985 | Tropilaelaps mercedesae | 89.6 | 87.9 | 1.7 | 14303 | 85.9 | 84.1 | 1.8 |
| GCA_003018175.1 | 6973 | Blattella germanica | 96.5 | 94 | 2.5 | 28676 | 84.9 | 82.9 | 2 |
| GCA_003335185.2 | 195883 | Laodelphax striatellus | 97 | 93.4 | 3.6 | 17512 | 95 | 90.9 | 4.1 |
| GCA_003675905.2 | 299467 | Leptotrombidium deliense | 86 | 84.9 | 1.1 | 14667 | 83.2 | 82.2 | 1 |
| GCA_003675995.1 | 1965070 | Dinothrombium tinctorium | 93.6 | 54.3 | 39.3 | 19024 | 88.5 | 54.4 | 34.1 |
| GCA_005959815.1 | 7385 | Sarcophaga bullata | 96.8 | 94.9 | 1.9 | 15763 | 87.8 | 86.2 | 1.6 |
| GCA_006783055.1 | 1923959 | Trinorchestia longiramus | 86.4 | 83.1 | 3.3 | 26080 | 86.5 | 83 | 3.5 |
| GCA_007210705.1 | 6832 | Tigriopus californicus | 93.5 | 92.8 | 0.7 | 15577 | 91.9 | 90.6 | 1.3 |
| GCA_009176605.1 | 96803 | Armadillidium nasatum | 89.3 | 83.1 | 6.2 | 14636 | 75.3 | 70.5 | 4.8 |
| GCA_009739505.2 | 248454 | Apolygus lucorum | 96.5 | 91.9 | 4.6 | 20111 | 94.3 | 89.8 | 4.5 |
| GCA_009805615.1 | 1232801 | Amphibalanus amphitrite | 92.5 | 64 | 28.5 | 25581 | 85.2 | 61.3 | 23.9 |
| GCA_011009095.1 | 2038154 | Ignelater luminosus | 96.9 | 94.4 | 2.5 | 27558 | 94.5 | 92.1 | 2.4 |
| GCA_013167095.2 | 1820382 | Daphnia sinensis | 98.5 | 96.3 | 2.2 | 16955 | 97.4 | 94.8 | 2.6 |
| GCA_013340265.1 | 36987 | Coptotermes formosanus | 98.5 | 97.3 | 1.2 | 12983 | 90.1 | 88.7 | 1.4 |
| GCA_013358835.2 | 34615 | Ixodes persulcatus | 88.2 | 84.1 | 4.1 | 28396 | 78.2 | 75.9 | 2.3 |
| GCA_015342795.1 | 94029 | Argiope bruennichi | 92.7 | 89.4 | 3.3 | 23259 | 87.3 | 83.8 | 3.5 |
| GCA_016920775.1 | 1564500 | Bradysia odoriphaga | 97.8 | 94.9 | 2.9 | 16206 | 95 | 92.2 | 2.8 |
| GCA_018290095.1 | 319348 | Polypedilum vanderplanki | 98.7 | 96.9 | 1.8 | 17863 | 98.7 | 96.7 | 2 |
| GCA_019049445.1 | 860918 | Ampulex compressa | 99.2 | 99.2 | 0 | 18896 | 97.3 | 97.1 | 0.2 |
| GCA_019343175.1 | 931172 | Oedothorax gibbosus | 95.2 | 91.4 | 3.8 | 29717 | 95.7 | 91.6 | 4.1 |
| GCA_019925095.2 | 765133 | Dendrolimus kikuchii | 97.6 | 96.2 | 1.4 | 15321 | 95.4 | 93.9 | 1.5 |
| GCA_021650745.2 | 40697 | Blomia tropicalis | 90.3 | 88.9 | 1.4 | 16592 | 90.4 | 88.3 | 2.1 |
| GCA_021730765.1 | 59818 | Tyrophagus putrescentiae | 82.3 | 78.5 | 3.8 | 23009 | 86.5 | 82.7 | 3.8 |
| GCA_022750525.1 | 2874060 | Halotydeus destructor | 90.5 | 89 | 1.5 | 14583 | 89.5 | 88.1 | 1.4 |
| GCA_024506315.2 | 121845 | Diaphorina citri | 92.1 | 89.2 | 2.9 | 26564 | 95.4 | 90.8 | 4.6 |
| GCA_025370935.1 | 7141 | Choristoneura fumiferana | 97.4 | 91.2 | 6.2 | 16231 | 83.1 | 77.4 | 5.7 |
| GCA_026979955.1 | 439358 | Megalurothrips usitatus | 97 | 96.2 | 0.8 | 11617 | 90.3 | 89.2 | 1.1 |
| GCA_029955175.1 | 1586481 | Ecocentrus adspersus | 99.5 | 99.2 | 0.3 | 16908 | 88.5 | 87.7 | 0.8 |
| GCA_030143305.2 | 6943 | Amblyomma americanum | 85.5 | 70.9 | 14.6 | 34557 | 79.9 | 71 | 8.9 |
| GCA_030220185.1 | 6984 | Diploptera punctata | 99.2 | 92.6 | 6.6 | 28414 | 86 | 81.5 | 4.5 |
| GCA_032884065.1 | 6661 | Artemia franciscana | 89.4 | 81.5 | 7.9 | 20122 | 81.3 | 75.4 | 5.9 |
| GCA_033782935.1 | 88211 | Petrolisthes cinctipes | 92 | 75.2 | 16.8 | 44511 | 94.4 | 77 | 17.4 |
| GCA_034508575.1 | 1843537 | Petrolisthes manimaculis | 92.4 | 79.7 | 12.7 | 40296 | 95.1 | 82.1 | 13 |
| GCA_037055365.1 | 468196 | Polyplax serrata | 98.7 | 98.7 | 0 | 14991 | 97.3 | 96.9 | 0.4 |
| GCA_037179515.1 | 373956 | Halocaridina rubra | 88.5 | 86.2 | 2.3 | 25341 | 82.5 | 81.4 | 1.1 |
| GCA_038050395.1 | 536013 | Parthenolecanium corni | 95 | 89.7 | 5.3 | 14951 | 86.4 | 82.4 | 4 |
| GCA_038098605.1 | 2509291 | Gryllus longicercus | 99 | 90.3 | 8.7 | 14730 | 95.6 | 87.6 | 8 |
| GCA_038387795.1 | 85552 | Scylla paramamosain | 95.1 | 93.5 | 1.6 | 20946 | 89.9 | 88.4 | 1.5 |
| GCA_038502225.1 | 27406 | Cherax quadricarinatus | 88.8 | 86.9 | 1.9 | 17532 | 88.3 | 86.4 | 1.9 |
| GCA_040020575.1 | 488301 | Rhynocoris fuscipes | 98.5 | 95.1 | 3.4 | 13362 | 86.6 | 82.9 | 3.7 |
| GCA_040285465.1 | 328185 | Menopon gallinae | 98.8 | 98.2 | 0.6 | 10891 | 88.5 | 87.5 | 1 |
| GCA_902825445.1 | 874455 | Arctia plantaginis | 98.8 | 97.9 | 0.9 | 18402 | 98.5 | 97.5 | 1 |
| GCA_902829235.1 | 197152 | Cloeon dipterum | 97.8 | 95.4 | 2.4 | 16299 | 97 | 94.5 | 2.5 |
| GCA_902850365.2 | 168631 | Chilo suppressalis | 98.9 | 98.2 | 0.7 | 11366 | 95.5 | 94.9 | 0.6 |
| GCA_905338385.1 | 69355 | Darwinula stevensoni | 90.3 | 88.1 | 2.2 | 15334 | 84.5 | 82.5 | 2 |
| GCA_905338405.1 | 399045 | Notodromas monacha | 86.2 | 81.1 | 5.1 | 13723 | 80.8 | 76.7 | 4.1 |
| GCA_905368565.1 | 1979941 | Medioppia subpectinata | 85.4 | 75.8 | 9.6 | 23249 | 81.1 | 72.9 | 8.2 |
| GCA_905397405.1 | 334625 | Oppiella nova | 85.1 | 75.4 | 9.7 | 23525 | 82 | 72.8 | 9.2 |
| GCA_910591605.1 | 39272 | Allacma fusca | 91 | 89.2 | 1.8 | 47610 | 91.8 | 89.7 | 2.1 |
| GCA_918026855.4 | 80249 | Phaedon cochleariae | 99.7 | 92.7 | 7 | 13141 | 96 | 89.7 | 6.3 |
| GCA_918697745.1 | 27404 | Daphnia galeata | 95.2 | 94.2 | 1 | 15649 | 93.1 | 91.8 | 1.3 |
| GCA_918797505.1 | 7038 | Bemisia tabaci | 98.5 | 96.1 | 2.4 | 14348 | 96.3 | 94.1 | 2.2 |
| GCA_928085145.1 | 85310 | Nezara viridula | 98.4 | 95.8 | 2.6 | 15971 | 91.3 | 88.9 | 2.4 |
| GCA_964023285.1 | 280406 | Larinioides sclopetarius | 98.5 | 93.8 | 4.7 | 22843 | 95.7 | 91 | 4.7 |
| GCA_964058975.1 | 48144 | Meganyctiphanes norvegica | 75.7 | 71.7 | 4 | 42189 | 85.3 | 82 | 3.3 |
| GCF_000001215.4 | 7227 | Drosophila melanogaster | 99.6 | 99.1 | 0.5 | 13986 | 99.9 | 99 | 0.9 |
| GCF_000006295.1 | 121224 | Pediculus humanus | 97.1 | 96.9 | 0.2 | 10758 | 97 | 96.7 | 0.3 |
| GCF_000239435.1 | 32264 | Tetranychus urticae | 90.1 | 84.2 | 5.9 | 11686 | 91.6 | 85.1 | 6.5 |
| GCF_000255335.2 | 34638 | Galendromus occidentalis | 93.9 | 90.8 | 3.1 | 11566 | 96.1 | 92 | 4.1 |
| GCF_000341935.2 | 211228 | Cephus cinctus | 99.2 | 98.9 | 0.3 | 11511 | 99.6 | 99.2 | 0.4 |
| GCF_000365465.3 | 114398 | Parasteatoda tepidariorum | 97 | 93.2 | 3.8 | 19750 | 98.3 | 93.3 | 5 |
| GCF_000517525.1 | 6850 | Limulus polyphemus | 92.3 | 78.9 | 13.4 | 22873 | 93.9 | 80.4 | 13.5 |
| GCF_000648675.2 | 79782 | Cimex lectularius | 99.4 | 97.8 | 1.6 | 11949 | 99.5 | 98.2 | 1.3 |
| GCF_000648695.1 | 166361 | Onthophagus taurus | 99.6 | 96.7 | 2.9 | 14537 | 99.8 | 96.2 | 3.6 |
| GCF_000671375.1 | 218467 | Centruroides sculpturatus | 93.9 | 89.4 | 4.5 | 24591 | 93.7 | 88.4 | 5.3 |
| GCF_000696155.1 | 136037 | Zootermopsis nevadensis | 98.9 | 98.5 | 0.4 | 12381 | 98.4 | 98.1 | 0.3 |
| GCF_000697945.3 | 133901 | Frankliniella occidentalis | 98.3 | 97.3 | 1 | 16516 | 99.5 | 97.9 | 1.6 |
| GCF_000699045.2 | 224129 | Agrilus planipennis | 97.7 | 94.1 | 3.6 | 13371 | 97.8 | 90.3 | 7.5 |
| GCF_000764305.2 | 294128 | Hyalella azteca | 93.7 | 93 | 0.7 | 17163 | 95.6 | 94.6 | 1 |
| GCF_001272555.1 | 178035 | Dufourea novaeangliae | 99.8 | 99.7 | 0.1 | 9844 | 99.7 | 99.5 | 0.2 |
| GCF_001412225.1 | 110193 | Nicrophorus vespilloides | 99.5 | 98.3 | 1.2 | 12642 | 100 | 98.6 | 1.4 |

| | | | | | | | | | |
|---|---|---|---|---|---|---|---|---|---|
| GCF_001412515.2 | 454923 | Diachasma alloeum | 99.5 | 99 | 0.5 | 12351 | 99.1 | 98.5 | 0.6 |
| GCF_001594045.1 | 520822 | Atta colombica | 99.2 | 98.7 | 0.5 | 10316 | 99.3 | 98.7 | 0.6 |
| GCF_001901225.1 | 6956 | Dermatophagoides pteronyssinus | 86 | 84.8 | 1.2 | 11184 | 92.1 | 90.7 | 1.4 |
| GCF_002217175.1 | 158441 | Folsomia candida | 97.5 | 95.9 | 1.6 | 24221 | 97.9 | 96.1 | 1.8 |
| GCF_002443255.1 | 109461 | Varroa destructor | 94.2 | 92.7 | 1.5 | 10260 | 95.2 | 93.4 | 1.8 |
| GCF_002532875.1 | 62625 | Varroa jacobsoni | 94.2 | 92.8 | 1.4 | 10739 | 95.8 | 94.3 | 1.5 |
| GCF_002706865.2 | 69820 | Spodoptera litura | 98.8 | 97.6 | 1.2 | 14745 | 99 | 97.6 | 1.4 |
| GCF_002891405.2 | 105785 | Cryptotermes secundus | 98.2 | 96.4 | 1.8 | 13170 | 98.3 | 96.8 | 1.5 |
| GCF_003254395.2 | 7460 | Apis mellifera | 99.8 | 99.8 | 0 | 9935 | 99.2 | 99 | 0.2 |
| GCF_003426905.1 | 7515 | Ctenocephalides felis | 96 | 69.4 | 26.6 | 18878 | 95.8 | 58.8 | 37 |
| GCF_003676215.2 | 43146 | Rhopalosiphum maidis | 98.6 | 93.8 | 4.8 | 12060 | 98.6 | 93 | 5.6 |
| GCF_003789085.1 | 6689 | Penaeus vannamei | 85 | 78.1 | 6.9 | 24987 | 93.7 | 82.5 | 11.2 |
| GCF_008802855.1 | 7054 | Photinus pyralis | 99.4 | 89.9 | 9.5 | 20647 | 98.8 | 85.7 | 13.1 |
| GCF_009176525.2 | 265458 | Contarinia nasturtii | 97.6 | 96.2 | 1.4 | 14889 | 99.5 | 97.7 | 1.8 |
| GCF_010614865.2 | 202533 | Stegodyphus dumicola | 84.1 | 80.9 | 3.2 | 20729 | 87.5 | 84.8 | 2.7 |
| GCF_011947565.2 | 41117 | Pollicipes pollicipes | 90.9 | 69.1 | 21.8 | 20444 | 93.3 | 57.3 | 36 |
| GCF_012274295.1 | 473952 | Osmia lignaria | 99.4 | 99.1 | 0.3 | 10556 | 99 | 98.6 | 0.4 |
| GCF_012932325.1 | 161013 | Thrips palmi | 96.9 | 95.9 | 1 | 14332 | 97.5 | 96.4 | 1.1 |
| GCF_013123115.1 | 935657 | Colletes gigas | 99.1 | 97.7 | 1.4 | 10168 | 99.5 | 98.1 | 1.4 |
| GCF_013339695.2 | 34632 | Rhipicephalus sanguineus | 92.5 | 86.7 | 5.8 | 22510 | 97.2 | 89.9 | 7.3 |
| GCF_013339725.1 | 6941 | Rhipicephalus microplus | 90.8 | 87 | 3.8 | 18598 | 95.1 | 90.6 | 4.5 |
| GCF_014805625.1 | 7396 | Glossina fuscipes | 99.4 | 96.1 | 3.3 | 12387 | 99.7 | 96.5 | 3.2 |
| GCF_014839805.1 | 7130 | Manduca sexta | 98.3 | 89.8 | 8.5 | 15967 | 99.3 | 89.1 | 10.2 |
| GCF_015228065.1 | 6687 | Penaeus monodon | 83.7 | 81.6 | 2.1 | 24092 | 91.3 | 87.3 | 4 |
| GCF_016920785.2 | 6945 | Ixodes scapularis | 90.5 | 86 | 4.5 | 26659 | 98.5 | 93.9 | 4.6 |
| GCF_017591435.1 | 210409 | Portunus trituberculatus | 93.5 | 92.9 | 0.6 | 17292 | 97.2 | 96.3 | 0.9 |
| GCF_018991925.1 | 6706 | Homarus americanus | 93.2 | 92.2 | 1 | 22368 | 97.6 | 96.3 | 1.3 |
| GCF_019202785.1 | 139456 | Penaeus chinensis | 90.8 | 89.8 | 1 | 20076 | 95.8 | 94.5 | 1.3 |
| GCF_019457755.1 | 32260 | Venturia canescens | 99.6 | 98.9 | 0.7 | 11831 | 99.5 | 98.2 | 1.3 |
| GCF_020424385.1 | 6728 | Procambarus clarkii | 94.4 | 93.2 | 1.2 | 26417 | 98.3 | 96.5 | 1.8 |
| GCF_020631705.1 | 35525 | Daphnia magna | 98.6 | 94.4 | 4.2 | 16891 | 97.8 | 91.7 | 6.1 |
| GCF_021130785.1 | 197043 | Homalodisca vitripennis | 83.3 | 73.7 | 9.6 | 19904 | 91.9 | 80.1 | 11.8 |
| GCF_021134715.1 | 6669 | Daphnia pulex | 98.1 | 97.9 | 0.2 | 15295 | 97.3 | 96.3 | 1 |
| GCF_021155785.1 | 2872261 | Neodiprion fabricii | 99.4 | 99.1 | 0.3 | 11597 | 99.5 | 99.2 | 0.3 |
| GCF_021234035.1 | 35523 | Daphnia pulicaria | 98.8 | 97.2 | 1.6 | 16865 | 97.8 | 95.2 | 2.6 |
| GCF_021461385.2 | 274613 | Schistocerca piceifrons | 97.5 | 94.1 | 3.4 | 17490 | 98.9 | 95.7 | 3.2 |
| GCF_021461395.2 | 7009 | Schistocerca americana | 98.2 | 96.2 | 2 | 17662 | 99.1 | 96.9 | 2.2 |
| GCF_022539665.1 | 120202 | Daphnia carinata | 98.7 | 97.6 | 1.1 | 13404 | 97.9 | 96.2 | 1.7 |
| GCF_022605725.1 | 2921223 | Anthonomus grandis | 99.8 | 97.7 | 2.1 | 13112 | 99.1 | 96.9 | 2.2 |
| GCF_023375885.1 | 34620 | Dermacentor andersoni | 93.7 | 90.6 | 3.1 | 22938 | 98.6 | 95.8 | 2.8 |
| GCF_023864275.1 | 274614 | Schistocerca cancellata | 98.1 | 95.4 | 2.7 | 16907 | 99.4 | 96.2 | 3.2 |
| GCF_023864345.2 | 2023355 | Schistocerca serialis | 98 | 95.8 | 2.2 | 17237 | 99.1 | 97.2 | 1.9 |
| GCF_023897955.1 | 7010 | Schistocerca gregaria | 98.3 | 94.1 | 4.2 | 19799 | 99.2 | 96 | 3.2 |
| GCF_024362695.1 | 13191 | Pectinophora gossypiella | 98.8 | 97.2 | 1.6 | 14107 | 99.7 | 97.8 | 1.9 |
| GCF_024364675.1 | 116153 | Aethina tumida | 99.8 | 98 | 1.8 | 13131 | 99.5 | 97.6 | 1.9 |
| GCF_024763615.1 | 29031 | Phlebotomus papatasi | 95.6 | 94 | 1.6 | 11610 | 97.6 | 95.9 | 1.7 |
| GCF_025091365.1 | 58002 | Daktulosphaira vitifoliae | 98.7 | 95.8 | 2.9 | 14650 | 98.5 | 95.1 | 3.4 |
| GCF_028554725.1 | 28588 | Zeugodacus cucurbitae | 99.3 | 99 | 0.3 | 13888 | 99.4 | 98.7 | 0.7 |
| GCF_029603195.1 | 2530218 | Condylostylus longicornis | 98.7 | 98 | 0.7 | 12227 | 98.9 | 97.4 | 1.5 |
| GCF_030269925.1 | 7091 | Bombyx mori | 99.4 | 99 | 0.4 | 13459 | 99.2 | 98.5 | 0.7 |
| GCF_030625045.1 | 688607 | Achroia grisella | 98.9 | 95.7 | 3.2 | 13873 | 99.9 | 96.5 | 3.4 |
| GCF_031216515.1 | 2078957 | Neocloeon triangulifer | 97.8 | 96.9 | 0.9 | 12721 | 98.6 | 97 | 1.6 |
| GCF_031307605.1 | 7070 | Tribolium castaneum | 99.8 | 99.4 | 0.4 | 12172 | 99.3 | 98.5 | 0.8 |
| GCF_037043105.1 | 334116 | Vanessa tameamea | 98.9 | 98.4 | 0.5 | 11875 | 99.8 | 99.6 | 0.2 |
| GCF_037126465.1 | 34597 | Ornithodoros turicata | 96.4 | 92.1 | 4.3 | 22416 | 98.3 | 93.1 | 5.2 |
| GCF_905115235.1 | 343691 | Hermetia illucens | 98.8 | 98.4 | 0.4 | 14004 | 99.5 | 99 | 0.5 |
| GCF_905147105.1 | 7116 | Pieris brassicae | 99.1 | 98.7 | 0.4 | 11885 | 99.6 | 99.3 | 0.3 |
| GCF_905475395.1 | 189513 | Chrysoperla carnea | 98 | 97.4 | 0.6 | 12985 | 98 | 97 | 1 |
| GCF_907165205.1 | 41139 | Coccinella septempunctata | 99.6 | 98.8 | 0.8 | 14769 | 99.7 | 98.3 | 1.4 |
| GCF_910589235.1 | 7445 | Vespa crabro | 99.6 | 99.3 | 0.3 | 10204 | 99.5 | 98.9 | 0.6 |
| GCF_912999745.1 | 76193 | Papilio machaon | 99 | 98.1 | 0.9 | 13379 | 99.8 | 98.8 | 1 |
| GCF_932276165.1 | 51655 | Plutella xylostella | 99.3 | 98.8 | 0.5 | 13971 | 99.6 | 98.9 | 0.7 |
| GCF_943734665.1 | 42839 | Anopheles aquasalis | 98.1 | 97.4 | 0.7 | 11511 | 99.6 | 98.4 | 1.2 |
| GCF_963082655.1 | 35570 | Stomoxys calcitrans | 99.2 | 98.7 | 0.5 | 15178 | 99.5 | 99 | 0.5 |
| this_study | 41365 | Scolopendra cingulata | 98.3 | 96.2 | 2.1 | 17955 | 97.6 | 95.2 | 2.4 |
| this_study | 29022 | Scutigera coleoptrata | 97.1 | 96.1 | 1 | 18157 | 98.1 | 97 | 1.1 |
| this_study | 2973175 | Archispirostreptus syriacus | 96.5 | 94.2 | 2.3 | 11437 | 95.1 | 92.6 | 2.5 |

**Supplementary Table 1: Genomic resources.**

Assembly accessions, protein-coding gene counts, and BUSCO assessments from *de novo* sequenced myriapod species and the publicly available annotated proteomes from NCBI.

| Metric | Value |
|---|---|
| Number of sequences in the alignment | 145 |
| Number of sites in the alignment | 76471 |
| Pairs of sequences | 10440 |
| Completeness (C) score for the alignment (Ca) | 0.772771 |
| Maximum C-score for individual sequences (Cr_max) | 0.863412 |
| Minimum C-score for individual sequences (Cr_min) | 0.305554 |
| Maximum C-score for individual sites (Cc_max) | 0.986207 |
| Minimum C-score for individual sites (Cc_min) | 0.006897 |
| Maximum C-score for pairs of sequences (Cij_max, i!=j) | 0.857488 |
| Minimum C-score for pairs of sequences (Cij_min, i!=j) | 0.167057 |

**Supplementary Table 2: Metrics of the concatenated super-alignment used for species phylogeny reconstruction.**

The species phylogeny was computed from single-copy orthologues found in at least 95% of the total set of 145 species, detected by the BUSCO assessments performed for the A3Cat on the genome assemblies using the arthropoda_odb10 lineage dataset. A concatenated super-alignment was built from the trimmed individual multiple alignments (76’471 columns, 71’014 distinct patterns, 57’142 parsimony-informative, 9’412 singleton sites, 9’917 constant sites).

| Last Common Ancestor | Min estimate | Max estimate |
|---|---|---|
| Chelicerata | 511.8 | 632 |
| Hexapoda | 425.4 | 478.1 |
| Insecta | 370.9 | 451.7 |
| PanCrustacea | 501.8 | 580 |
| Myriapoda | 442 | 550.9 |
| Chilopoda | 411.7 | 442.7 |
| Malacostraca | 254.2 | 444 |
| Branchiopoda | 365.1 | 491.7 |
| Hymenoptera,Lepidoptera | 310.7 | 389.7 |
| Root | 536.6 | 600 |

**Supplementary Table 3: Time estimates for species phylogeny tree calibration.**

Divergence time estimates were sourced from the TimeTree v5 database (Kumar et al. 2022) to time-calibrate the molecular species tree.

| TaxId | Gene name | Moulting pathway | NCBI prot ID |
|---|---|---|---|
| 7227 | Bftz-F1 | Early_genes | NP_524143.2 |
| 7227 | Blimp-1 | Early_genes | NP_001261442.1 |
| 7227 | crc | Early_genes | NP_524897.1 |
| 7227 | E74 | Early_genes | NP_730288.1 |
| 7227 | E75 | Early_genes | NP_730321.1 |
| 7227 | E78 | Early_genes | NP_524195.2 |
| 7227 | Hnf4 | Early_genes | NP_723413.1 |
| 7227 | HR3 | Early_genes | NP_001334718.1 |
| 7227 | HR38 | Early_genes | NP_477119.1 |
| 7227 | HR39 | Early_genes | NP_476932.1 |
| 7227 | HR4 | Early_genes | NP_001259161.1 |
| 7227 | HR78 | Early_genes | NP_730636.1 |
| 7227 | HR96 | Early_genes | NP_524493.1 |
| 7227 | tll | Early_genes | NP_524596.1 |
| 7227 | Atet | Ecdysteroid_pathway | NP_001097079.1 |
| 7227 | cyp18a1 | Ecdysteroid_pathway | NP_728191.1 |
| 7227 | dib | Ecdysteroid_pathway | NP_524810.2 |
| 7227 | ecd | Ecdysteroid_pathway | NP_647707.1 |
| 7227 | EcI | Ecdysteroid_pathway | NP_648989.1 |
| 7227 | EcR | Ecdysteroid_pathway | NP_724460.1 |
| 7227 | gcn5 | Ecdysteroid_pathway | NP_648586.2 |
| 7227 | MSBP | Ecdysteroid_pathway | NP_573087.1 |
| 7227 | nvd | Ecdysteroid_pathway | NP_001097670.1 |
| 7227 | phm | Ecdysteroid_pathway | NP_573319.1 |
| 7227 | sad | Ecdysteroid_pathway | NP_650123.1 |
| 7227 | shd | Ecdysteroid_pathway | NP_001261843.1 |
| 7227 | spo | Ecdysteroid_pathway | NP_647975.2 |
| 7227 | spok | Ecdysteroid_pathway | NP_001104460.2 |
| 7227 | SREBP | Ecdysteroid_pathway | NP_730449.1 |
| 7227 | sro | Ecdysteroid_pathway | NP_651725.1 |
| 7227 | Usp | Ecdysteroid_pathway | NP_476781.1 |
| 7227 | Broad | Fate_genes | NP_001162638.2 |
| 7227 | chinmo | Fate_genes | NP_001188680.1 |
| 7227 | E93 | Fate_genes | NP_001097865.1 |
| 7227 | Kr-h1 | Fate_genes | NP_477466.1 |
| 7227 | Cda3 | Late_genes | NP_609806.1 |
| 7227 | Cda4 | Late_genes | NP_728468.1 |
| 7227 | Cda5 | Late_genes | NP_722590.2 |
| 7227 | Cda9 | Late_genes | NP_611192.1 |
| 7227 | Chs2 | Late_genes | NP_524209.3 |
| 7227 | Cht11 | Late_genes | NP_572361.1 |
| 7227 | Cht2 | Late_genes | NP_477298.2 |
| 7227 | Cht3 | Late_genes | NP_001036422.1 |
| 7227 | Cht4 | Late_genes | NP_524962.2 |
| 7227 | Cht5 | Late_genes | NP_650314.1 |
| 7227 | Cht6 | Late_genes | NP_001245598.1 |
| 7227 | Cht7 | Late_genes | NP_647768.3 |
| 7227 | Cht8 | Late_genes | NP_611542.2 |
| 7227 | Cht9 | Late_genes | NP_611543.3 |

| | | | |
|---|---|---|---|
| 7227 | FABP | Late_genes | NP_001027179.1 |
| 7227 | kkv | Late_genes | NP_730928.2 |
| 7227 | knk | Late_genes | NP_649981.1 |
| 7227 | mco1 | Late_genes | NP_609287.3 |
| 7227 | PPO1 | Late_genes | NP_476812.1 |
| 7227 | PPO2 | Late_genes | NP_610443.1 |
| 7227 | PPO3 | Late_genes | NP_524760.1 |
| 7227 | resilin | Late_genes | NP_611157.1 |
| 7227 | rtv | Late_genes | NP_572693.1 |
| 7227 | serp | Late_genes | NP_730444.1 |
| 7227 | stw | Late_genes | NP_610170.2 |
| 7227 | verm | Late_genes | NP_001163469.1 |
| 7227 | CCAP-R | Neurostimuli_reception | NP_001368957.1 |
| 7227 | EHR | Neurostimuli_reception | NP_729905.2 |
| 7227 | ETHR | Neurostimuli_reception | NP_650960.2 |
| 7227 | rk | Neurostimuli_reception | NP_476702.1 |
| 7227 | torso | Neurostimuli_reception | NP_476762.1 |
| 7227 | Chd64 | Sesquiterpenoid_pathway | NP_647860.1 |
| 7227 | FKBP39 | Sesquiterpenoid_pathway | NP_524364.2 |
| 7227 | JHAMT | Sesquiterpenoid_pathway | NP_609793.2 |
| 7227 | JHBP | Sesquiterpenoid_pathway | NP_608420.1 |
| 7227 | JHEBP | Sesquiterpenoid_pathway | NP_611989.1 |
| 7227 | JHEH | Sesquiterpenoid_pathway | NP_611385.1 |
| 7227 | met | Sesquiterpenoid_pathway | NP_511126.2 |
| 7227 | Tai | Sesquiterpenoid_pathway | NP_001245949.1 |
| 6689 | HR3 | Early_genes | XP_027222334.1 |
| 6689 | dib | Ecdysteroid_pathway | XP_027209672.1 |
| 6689 | EcR | Ecdysteroid_pathway | XP_027212061.1 |
| 6689 | E74 | Early_genes | XP_027206747.1 |
| 6689 | E75 | Early_genes | XP_027211093.1 |
| 6689 | Bftz-F1 | Early_genes | XP_027236936.1 |
| 6689 | kkv | Late_genes | XP_027227734.1 |
| 6689 | knk | Late_genes | XP_027210805.1 |
| 6689 | sad | Ecdysteroid_pathway | XP_027238669.1 |
| 6689 | spo | Ecdysteroid_pathway | XP_027213908.1 |
| 6689 | Usp | Ecdysteroid_pathway | XP_027216942.1 |
| 6689 | E78 | Early_genes | XP_027234687.1 |
| 6689 | phm | Ecdysteroid_pathway | XP_027215959.1 |
| 6689 | JHEH | Sesquiterpenoid_pathway | XP_027222530.1 |
| 6689 | cyp18a1 | Ecdysteroid_pathway | XP_027215960.1 |
| 6689 | JHBP | Sesquiterpenoid_pathway | XP_027223816.1 |
| 6689 | Chd64 | Sesquiterpenoid_pathway | XP_027224334.1 |
| 6689 | JHEBP | Sesquiterpenoid_pathway | XP_027208769.1 |
| 6689 | Tai | Sesquiterpenoid_pathway | XP_027215744.1 |
| 6689 | SREBP | Ecdysteroid_pathway | XP_027238775.1 |
| 6689 | sro | Ecdysteroid_pathway | XP_027222633.1 |
| 6689 | HR4 | Early_genes | XP_027207778.1 |
| 6689 | Kr-h1 | Fate_genes | XP_027230918.1 |
| 6689 | nvd | Ecdysteroid_pathway | XP_027231585.1 |
| 210409 | spo | Ecdysteroid_pathway | XP_045129380.1 |

| | | | |
|---|---|---|---|
| 210409 | sad | Ecdysteroid_pathway | XP_045131452.1 |
| 210409 | dib | Ecdysteroid_pathway | XP_045105768.1 |
| 114398 | HR3 | Early_genes | XP_042902546.1 |
| 114398 | EcR | Ecdysteroid_pathway | XP_015926579.1 |
| 114398 | E74 | Early_genes | XP_021001446.2 |
| 114398 | E75 | Early_genes | XP_015930846.1 |
| 114398 | Bftz-F1 | Early_genes | XP_015908511.1 |
| 114398 | met | Sesquiterpenoid_pathway | XP_042896763.1 |
| 114398 | shd | Ecdysteroid_pathway | XP_042903873.1 |
| 114398 | Usp | Ecdysteroid_pathway | XP_015929720.1 |
| 114398 | E78 | Early_genes | XP_015903773.1 |
| 114398 | JHEH | Sesquiterpenoid_pathway | XP_042906018.1 |
| 114398 | cyp18a1 | Ecdysteroid_pathway | XP_015920197.2 |
| 114398 | JHAMT | Sesquiterpenoid_pathway | XP_015910904.1 |
| 114398 | JHEBP | Sesquiterpenoid_pathway | XP_015915401.1 |
| 114398 | CCAP-R | Neurostimuli_reception | XP_015921275.1 |
| 114398 | spo | Ecdysteroid_pathway | XP_015926868.1 |
| 114398 | SREBP | Ecdysteroid_pathway | XP_042907215.1 |
| 114398 | E93 | Fate_genes | XP_042896616.1 |
| 114398 | HR4 | Early_genes | XP_021000290.1 |
| 114398 | nvd | Ecdysteroid_pathway | XP_042901224.1 |
| 7091 | cyp15 | Sesquiterpenoid_pathway | NP_001140197.1 |
| 7070 | cyp15 | Sesquiterpenoid_pathway | NP_001308597.1 |

**Supplementary Table 4: Curated list of genes involved in the moulting process.**

| Gene family | Ancestor | Nr of ancestral family duplications | Nr of ancestral family losses |
|---|---|---|---|
| serp;verm | Mandibulata | 1 | 0 |
| serp;verm | Araneae | 1 | 0 |
| kkv;Chs2 | Chelicerata | 1 | 0 |
| kkv;Chs2 | Pancrustacea | 1 | 0 |
| knk | Arthropoda | 2 | 0 |
| Cda4 | Araneae | 1 | 0 |
| ppos | Non tetrapulmonata | 0 | 1 |
| ppos | Acari | 0 | 1 |
| nvd | Coleoptera | 0 | 1 |
| nvd | Mesostigmata | 0 | 1 |
| chinmo | Ostracoda | 0 | 1 |
| chinmo | Malacostraca | 0 | 1 |
| chinmo | Branchiopoda | 0 | 1 |
| E93 | Branchiopoda | 0 | 1 |

**Supplementary Table 5: Summary of deep time events from selected gene families.**

Total count of duplications inferred by reconciliation and losses inferred by gene prediction assessment are mapped to clade ancestors, as plotted in Figure 5.

| Query NCBI identifier | Species | Gene name |
|---|---|---|
| NP_001097865.1 | Drosophila melanogaster | E93 |
| XP_042896616.1 | Parasteatoda tepidariorum | E93 |
| XP_037795760.1 | Paeneus monodon | E93 |
| TRY61992.1 | Tigriopus californicus | chinmo |
| XP_015835513.1 | Tribolium castaneum | chinmo |
| NP_609287.3 | Drosophila melanogaster | mco1 |
| XP_045026750.1 | Daphnia magna | mco1 |
| XP_042224188.1 | Hommarus americanus | mco1 |
| XP_042229820.1 | Hommarus americanus | ppo1 |
| NP_476812.1 | Drosophila melanogaster | ppo1 |
| XP_042903490.1 | Parasteatoda tepidariorum | hemocyanin chain A |
| NP_001097670.1 | Drosophila melanogaster | nvd |
| XP_026296485.1 | Apis mellifera | nvd |
| XP_015795304.1 | Tetrachynus urticae | nvd |
| XP_042901224.1 | Parasteatoda tepidariorum | nvd |

**Supplementary Table 6: Query genes for gene prediction in unannotated assemblies.**
